# Ångström Resolution with Flow Immunofluorescence Localization Microscopy (FILM)

**DOI:** 10.64898/2025.12.16.694546

**Authors:** Johanna Bartmuß, Anke Leinhaas, Tatjana Frank-Wiebe, Marc Schmidt-Supprian, Ali Kinkhabwala

## Abstract

We present flow immunofluorescence localization microscopy (FILM) as a novel paradigm for single-molecule localization microscopy (SMLM). FILM is based on continuous flow of standard fluorescent binders, including conventional antibodies or nanoscale probes, over a fixed sample along with gentle imaging-to-photobleaching of the stochastically bound probes with an LED on a widefield microscope setup. Gentle illumination yields 10^4^–10^6^ photons per fluorophore corresponding to localization precision at the few nanometer to Ångström scales. We employ FILM to localize diverse markers in cultured cells and immunological synapses down to 9 Å resolution, and for sequential localization of multiple targets in tissue slices. Requiring only minimal hardware and off-the-shelf probes, FILM offers an accessible and versatile route for serial SMLM at molecular resolution.

## Introduction

Precise spatial mapping of proteins and protein complexes within cells is essential for understanding cellular function and organization. Traditionally, light microscopy has been limited in resolution by the diffraction barrier, restricting structural insights to greater than approximately 200 nm (*1*). Over the past three decades, a range of super-resolution techniques have emerged to overcome this limit, including STimulated Emission Depletion (STED) (*2*), which enables down to roughly 50 nm resolution by depletion of excited fluorophore signal at the periphery of a confocal detection volume, and single-molecule localization microscopy (SMLM), which achieves 15–20 nm resolution by the sparse detection and precise localization of isolated fluorescent probes (*3–6*). While these advances marked a major breakthrough in optical imaging, they still fell short of providing the necessary molecular resolution required to resolve individual protein complexes at the scale of a few nanometers.

Molecular-scale resolution was more recently achieved through the integration of SMLM with advanced imaging modalities such as cryogenic microscopy (cryo-SMLM) (*7–9*), structured excitation and probe triangulation (e.g., MINimal photon FLUXes, MINFLUX) (*10*), physical sample expansion (Expansion Microscopy, ExM) (*7*, *11*, *12*), or stochastic labeling and repeated localization of individual targets (Resolution Enhancement by Sequential Imaging, RESI) (*13*). Widespread adoption of these approaches however remains limited due to the need for sophisticated laser-based instrumentation along with specialized fluorophores, complex labeling strategies, demanding expansion protocols, or cryogenic chambers. Moreover, achieving reliable molecular resolution for multiple, densely packed targets within a sample remains a significant challenge for the field (*14*).

One particular set of single molecule approaches relies on the direct, stochastic binding/unbinding of transient probes (*6*, *15*) or binding/photobleaching of stable probes (*16*, *17*) to targets on a specimen from an immersing reservoir. For these approaches, it is critical to limit the photobleaching of the free fluorescent probes in the reservoir. Total Internal Reflection Fluorescence (TIRF) microscopy (*18*) or Highly Inclined and Laminated Optical sheet (HILO) microscopy (*19*) is therefore typically employed to delay the photobleaching of the freely diffusing probes as well as to reduce fluorescence background from the reservoir. However, restricted illumination only slows photobleaching but cannot eliminate it, with a steady accumulation of photobleached probes in the imaging volume nevertheless occurring over time (Fig. S1A–B), which is exacerbated with widefield illumination (Fig. S1C–D). Especially problematic for stable binders, these photobleached probes can bind to and block target sites, thereby reducing the achievable labeling efficiency.

Instead of a static reservoir, we reasoned that a *dynamic* reservoir of fluorescent probes generated by continuous flow, together with gentle illumination for detection and localization of the bound probes, should yield a low level of photobleaching of the free probe during its transit over the illuminated field of view (FOV) to targets within the sample. Continuous flow not only supplies the sample with fresh fluorescent probes at all times from upstream, but also simultaneously removes photobleached free probes to downstream, generating a shallow, steady-state photobleaching gradient for the free probe along the flow direction even for widefield illumination (Fig. S1E–F). Continuous flow as well supplies the sample with fresh buffer and removes reactive oxygen species (ROS) generated by the illumination to downstream, which should reduce the need for oxygen scavengers in the buffer. Unlike TIRF or HILO, widefield microscopy allows for imaging of the sample both homogeneously and at depth, features critical for flexible intracellular and tissue-level imaging. To facilitate the use of widefield microscopy, the probe concentration in the reservoir should be as low as possible to reduce background. The probe concentration also determines the rate of probe binding, which can be balanced with the photobleaching rate determined by the illumination intensity, to create a steady level of sparse labeling of the specimen for single fluorophore detection.

Here we realize this concept as Flow Immunofluorescence Localization Microscopy (FILM). Immunofluorescence based on flow, or *flow immunofluorescence*, can be contrasted with flow cytometry. For flow cytometry, cells flow through an illuminated volume that is continuously sampled by a detector; for flow immunofluorescence, fluorescent probes flow over and bind to targets within an illuminated field of view of a fixed sample that is continuously imaged by a camera. FILM is based on a standard widefield microscope equipped with conventional LED illumination and basic microfluidics for continuous delivery of off-the-shelf affinity probes to the sample through use of a pump or even simply gravity (Fig. 1A). While detection of single fluorescent probes with conventional fluorescence microscopy is not possible due to dense specimen labeling (Fig. 1B), continuous imaging of the gradual binding and photobleaching of isolated probes delivered by flow to the sample enables the recording of a super-resolution movie of all probe binding-to-photobleaching events within the illuminated FOV (Fig. 1C). For probes labeled with a standard dye with conventional photobleaching kinetics, optimal parameters of flow speed and applied illumination intensity can typically be found that ensure a shallow gradient of photobleaching of the free probe along the flow direction, allowing for super-resolved staining of the sample. Homogeneous imaging of the sample both laterally and at few micron depth within the sample requires that the sum of the transit time over the FOV and the diffusion time into the sample be significantly shorter than the photobleaching timescale (see boxed relation in Fig. 1C). The time for diffusive entry, *T*_diffusion_ = *h*/*D*^2^, into a sample of typical height, *h* = 5–10 µm, is at least 0.7–2.7 s for an antibody (hydrodynamic radius, *r* = 5.8 nm; diffusion constant, *D* = 37 µm^2^/s in water (*20*)) and at least 0.17–0.7 s for a hypothetical 3 nm probe (hydrodynamic radius, *r* = 1.5 nm; *D* = 143 µm^2^/s). This diffusion timescale, in particular, sets a fundamental lower limit of a few seconds on the photobleaching half-time, *T*_1/2,_, for FILM. In addition to these considerations, we furthermore optimize FILM for low background and high photon yields of individual probes by tuning the probe concentration and illumination, with consideration of the epitope distribution in the sample, for efficient SMLM. Using fluorescent antibodies, antibody Fab fragments, and phalloidin-Atto643, we report high-density, multiplexed FILM imaging in whole cells, immunological synapses, and tissue samples with high localization precision. Our results demonstrate a performance of FILM *on par* with or even exceeding current state-of-the-art technologies for SMLM, while maintaining a simple widefield imaging regimen. Together, our results highlight FILM-based super-resolution imaging as a straightforward approach for SMLM down to molecular resolution.

**Fig. 1.**
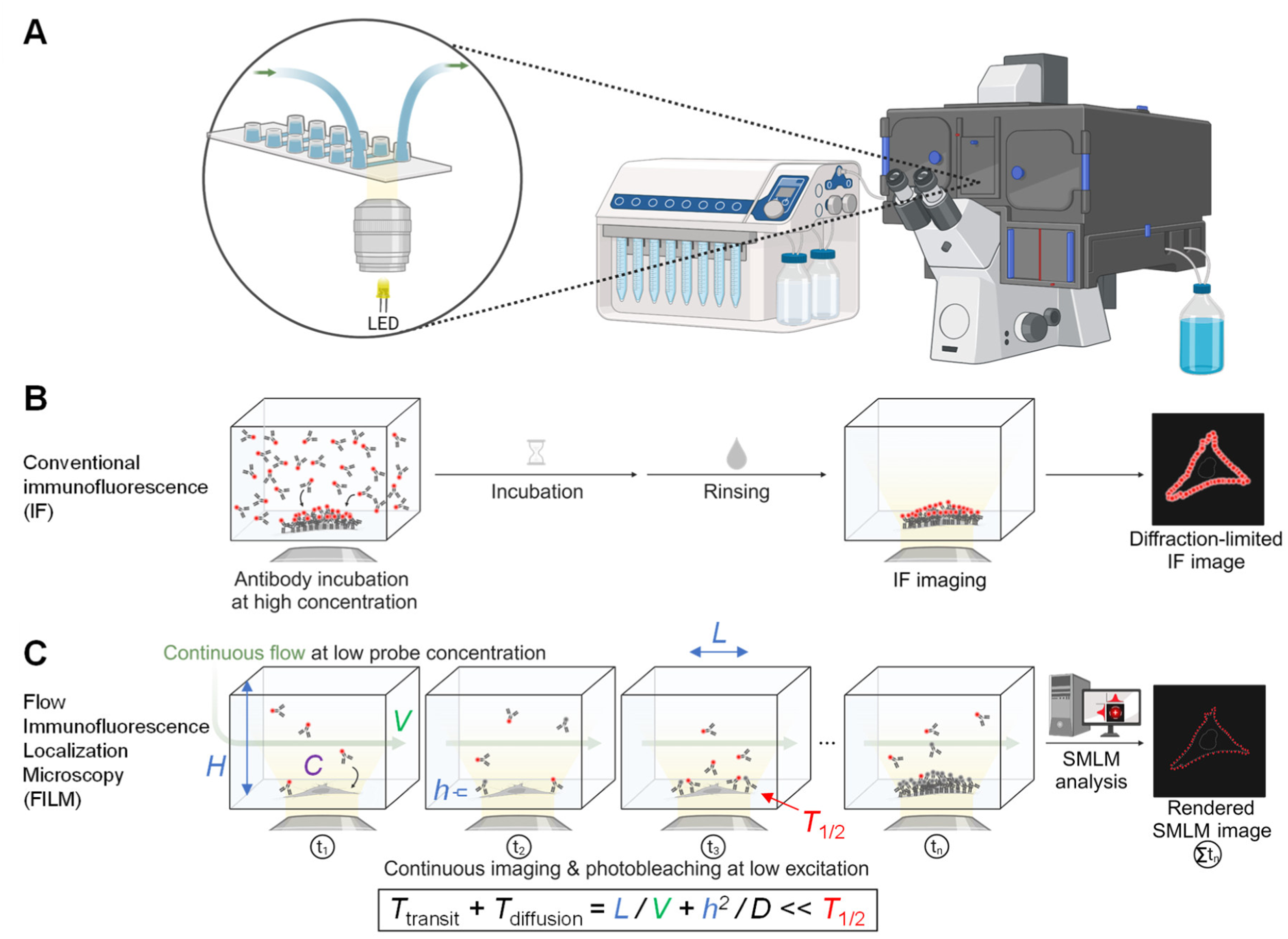
FILM hardware and concept. (A) Instrumentation for FILM. A pump, or even simply gravity, can be used to flow one or more solutions in a serial fashion over a fixed biological sample mounted in a microfluidic imaging chamber (green arrows indicate flow direction). Sample fluorescence is observed on a widefield microscope with LED illumination. (B) Diffraction-limited imaging using conventional immunofluorescence (IF). (C) Schematic representation of FILM. A low concentration of fluorescently labeled binder probes are delivered to the sample under continuous flow and with continuous imaging (yellow cone of excitation light emanating from the objective). Continuous detection to photobleaching of bound probes is balanced by continuous replenishment from upstream of fresh probes that bind the sample (depicted over the different timepoints: t_1_, …, t_n_). A small fraction of free probes will be photobleached during transit. Instead of accumulating near the sample over time, these photobleached probes are continuously flowed downstream to the waste. Probes can be conventional antibody or antibody-like binders used for IF that bind in either a stable or semi-stable fashion (down to binding half-times on the minute timescale). Single molecule binding events are detected and super-resolved using custom analysis software, resulting in a single super-resolution image containing all of the observed localizations. The condition for homogeneous labeling, both laterally and at depth within the sample, is given by the boxed expression. This mandates that the sum of the transit time, *T*_transit_, and diffusion time, *T*_diffusion_, be significantly less than the photobleaching half-time, *T*_1/2_. The transit time is equal to the illuminated FOV of length, *L*, divided by the flow velocity, *V*, and the time for diffusive entry of the probe into the sample can be approximated by the square of the sample height, *h*, divided by the diffusion constant relevant for the probe when in the sample, *D*. The density of fluorescent labels on the sample will be proportional to the applied probe concentration, *C*, with the contribution of background from free probes for widefield detection proportional to both *C* and the chamber height, *H*. *Created in BioRender. Bartmuß, J. (2024)* https://BioRender.com/j10h690

## Results

### Optimization of Photobleaching Half-Time and Photon Collection for FILM

To determine which illumination intensities yield photobleaching half-times on at least the seconds timescale required for homogeneous staining with FILM, we imaged isolated dye-labeled probes in phosphate-buffered saline (PBS, pH 7.4) buffer that were sparsely adsorbed on a coverslip at different illumination intensities until they were photobleached (Fig. S2A). We separately investigated an antibody probe conjugated to allophycocyanin (APC) (*21*) with degree-of-labeling (DOL) equal to unity (one APC per probe), and an antibody Fab fragment conjugated to Vio^®^ 667 with DOL = 1–2 (Materials and Methods). Using illumination intensities spanning 5.3 W cm^−2^ (using an LED at 100%) to 676 W cm^−2^ (100% laser), we determined photobleaching lifetime distributions and half-times for both probes (Figs. S2B–C). A broad distribution of half-times was observed, ranging from <10 ms to 25 s for APC and 0.15 ms to 40 s for Vio^®^ 667, from the highest to the lowest applied intensities. The photobleaching half-times of tens of seconds observed for each fluorophore at the lowest intensity corresponding to 100% LED should be sufficient for homogeneous staining with FILM.

For APC, a 6-bilin exciton-coupled complex, high irradiance promotes triplet formation and accelerates photobleaching (*21*). For organic dyes like Vio^®^ 667, photobleaching also increases nonlinearly with illumination intensity due to the population of long-lived triplet states and multi-photon absorbance (*22*). In both cases, accelerated photobleaching at high irradiance reduces the total number of photons that can be collected per fluorophore before photobleaching. Conversely, at sufficiently low intensities, the photobleaching quantum yield, *ϕ*_*b*_, defined as the probability of irreversible photobleaching upon absorption of a photon, should be constant. At these low intensities, each probe should emit a fixed number of photons before bleaching, independent of the applied illumination intensity. Instead of a constant number of photons collected from each probe, however, we observe an increasing number of photons for decreasing intensity, with a roughly hundred-fold overall increase over the assayed intensity range (Fig. S2D), implying a strong nonlinear dependence of photobleaching on intensity over this range. At the lowest applied intensity corresponding to 100% LED power, we detected 10⁴–10⁶ photons per probe for both the APC- and Vio^®^ 667-labeled probes.

To confirm the consistency of our results with theoretical expectations, we calculate the expected number of detected photons per probe until photobleaching based on the relevant parameters of illumination intensity, fluorophore extinction coefficient, photobleaching half-time, and instrument detection efficiency (adapted from (*22*)). The average number of photons absorbed by a fluorophore before photobleaching is equal to the sum of an infinite geometric series based on the probability of survival, *ϕ*_*s*_ = 1 − *ϕ*_*b*_, upon each successive absorption, or 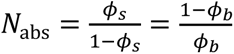. The latter multiplied by the quantum yield, *ϕ_F_*, gives the number of emitted photons, 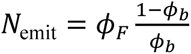. As photobleaching is rare (*ϕ_b_* ≪ 1), this can be simplified to Nemit ≈ *ϕ_F_*/*ϕ_b_*. Relating the rates of absorption and photobleaching allows for an explicit solution of *ϕ*_*b*_ in terms of the photobleaching half-time and other parameters, leading to the following expression of the average number of emitted photons:

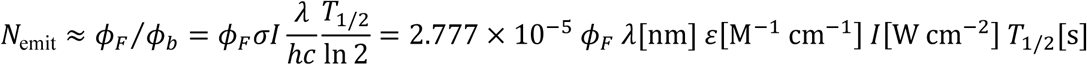

with *σ* the absorption cross-section, *I* the excitation intensity at the sample, *λ* the excitation wavelength, *h* Planck’s constant, *c* the speed of light, *T*_1/2_ the photobleaching half-time, and ε the extinction coefficient.

For the relevant parameters characterizing APC (*21*) of *ϕ*_*F*_ = 0.68 and ε = 7.0 × 10^5^ M^−1^ cm^−1^, along with the observed half-time of *T*_1/2_ = 25 s at 100% LED with *I* = 5.3 W cm^−2^ (applied at *λ* ≈ 640 nm), we obtain an expected average number of emitted photons of *N*_emit_ = 1.1 × 10^6^. The actually detected number of photons will be *N* = *ηN*_emit_, with *η* ≤ 1 the detection efficiency. The latter depends on the fraction of the emitted light, *f* = *Ω*/4*π* = (1 − cos *θ*)/2, collected over the solid angle, *Ω*, corresponding to our objective with *NA* = 1.46 and refractive index for oil immersion of *RI* = 1.53, with sin *θ* = *NA*/*RI* = 0.9542 and therefore cos *θ* = 0.299. The detection efficiency further depends on the ∼85% quantum efficiency of the camera at the peak emission wavelengths of APC (660 nm) and Vio^®^ 667 (667 nm) (Materials and Methods). These together give *η* ≈ 0.30. The theoretically expected average number of photons detected by our camera should then be *N* = 3.4 × 10^5^, agreeing well with the observed average of 3.3 × 10^5^ for APC (Fig. S2D, 100% LED). For Vio^®^ 667, the corresponding values are *ϕ*_*F*_ = 0.28 and ε = 2.15 × 10^5^ M^−1^ cm^−1^, with a range of DOL between 1–2 and an observed half-time of 40 s at 100% LED. These give a theoretically expected average number of photons detected by our camera of *N* = 0.68 − 1.4 × 10^5^, again consistent with the observed average of 1.4 × 10^5^ (Fig. S2D, 100% LED). These high numbers of detected photons for probes in PBS buffer agree well with literature values of ∼10^5^–10^6^ photons for single organic dyes in a more optimal, reduced oxygen buffer (*23*).

For SMLM, localization precision scales inversely with the square root of the detected photons, *N*, as 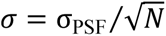, with *σ*_PSF_ the optical resolution (Gaussian width of the point spread function, PSF) (*24*). For far-red emission light, *σ*_PSF_ ≈ 100 nm, with >10,000 photons required to reach sub-nm precision (Materials and Methods). The 10^4^–10^6^ photons that we detected at 100% LED for both probes could be shown to correspond to localization precisions at the few nm to Ångstrom levels (Fig. S2E, Materials and Methods).

### FILM Imaging of Cytological Markers in Cultured Cells and Immunological Synapses at Molecular Resolution

The above experiments prove that gentle LED illumination generates sufficiently long photobleaching half-times (tens of seconds) required for homogeneous staining with FILM, and additionally permits much higher photon collection and corresponding localization precision. How well these results translate to the *in situ* setting of FILM, for which the probes are flowed over a sample to which they bind and are then detected with widefield microscopy, is however unclear. Namely, before each bound probe can be detected, it must first flow through the illuminated field of view some distance and then diffuse into the sample to its target (Fig. 1C). As detection is delayed with respect to the onset of probe illumination, a shorter *apparent* photobleaching half-time will be observed, with fewer photons collected than for the imaging-to-photobleaching of probes adsorbed onto glass assessed above. As well, the successful detection and precise localization of each probe may be hampered by the background from the free probe reservoir.

To resolve these questions, we attempted to image *α*-tubulin in fixed HeLa cells using a conventional monoclonal antibody conjugated with APC. Microtubules are composed of *αβ* tubulin heterodimers that connect end-to-end to form polar protofilaments, with 13 protofilaments arranged side-by-side forming a hollow cylindrical tube of 25 nm diameter (*12*, *25*, *26*) (Fig. 2A). The applied flow speed of 40 µL/min implied a transit time across the FOV of *T* = 0.65 s (Fig. 1C, Materials and Methods), which is significantly shorter than the photobleaching half-time of 25 s obtained for APC adsorbed onto glass at 100% LED (Fig. S2B). For the probe concentration, we settled on a dilution of 1:5,000 from the stock (Table S2), which is 100× lower than the staining concentration recommended for immunofluorescence. To generate the final images shown in Figs. 2B–C (full FOV displayed in Fig. S3) we recorded a movie comprised of 9,564 single plane frames with 5 s exposure each that was interleaved at 1-min intervals with short-exposure brightfield images (Fig. S4). A total of 14,176,305 localizations was recorded over the 13.3 hour acquisition.

**Fig. 2.**
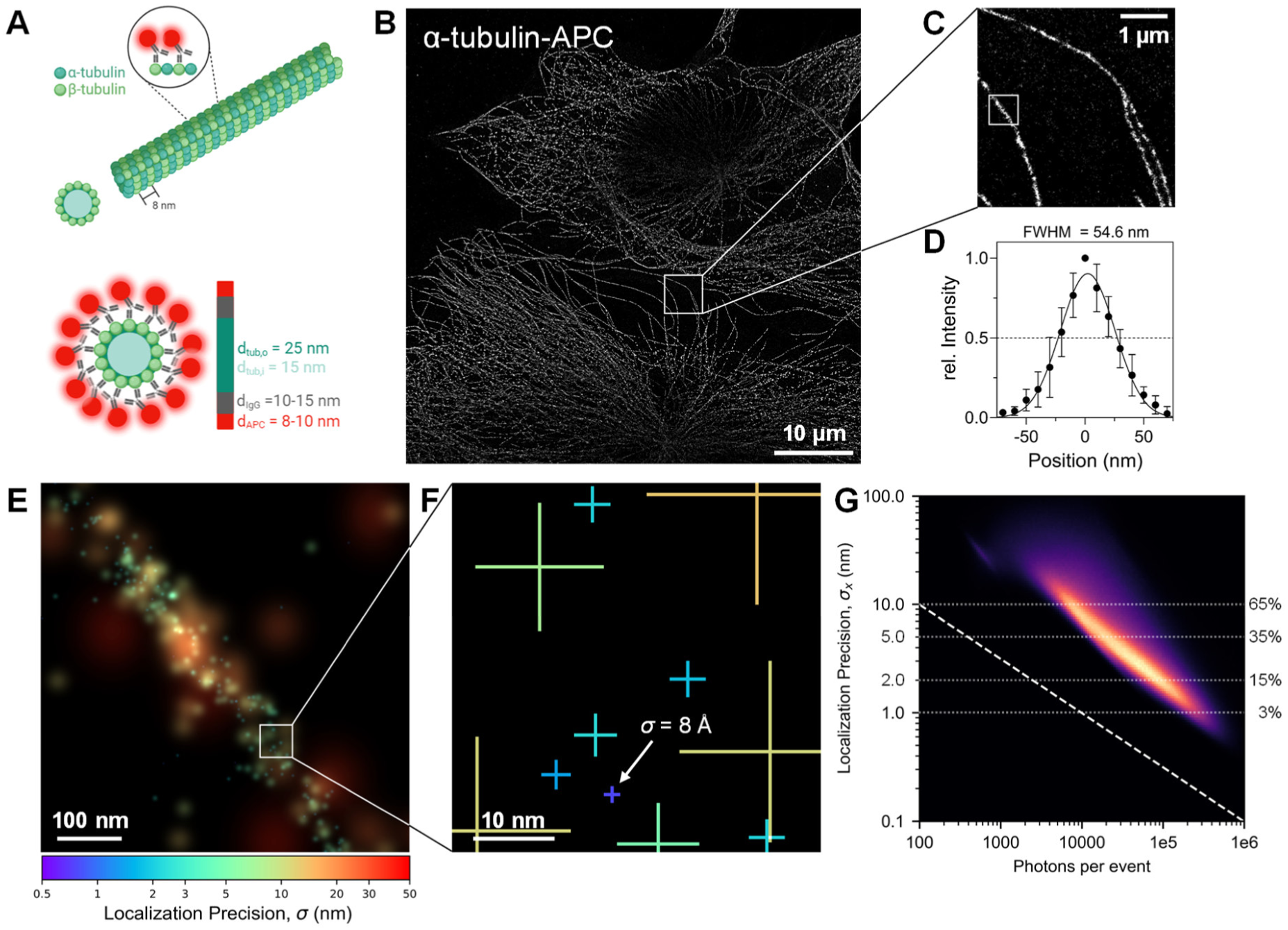
α-tubulin resolved in HeLa cells with FILM. (A) Schematic model of an antibody-decorated microtubule. The true microtubule size of d_Tub,o_ = 25 nm will appear larger due to the labeling. Staining with conventional primary antibodies (d_IgG_ = 10 nm) fluorescently labeled with APC (d_APC_ = 8 nm) results in a total diameter of ∼ 55 nm (= d_Tub,o_ + 2‧d_IgG_ + 2‧r_APC_). *Created in BioRender. Bartmuß, J. (2024)* https://BioRender.com/v24z991. (B) FILM imaging of anti-α-tubulin-APC antibodies labeling PFA-fixed and permeabilized HeLa cells (widefield illumination, 100% LED illumination, 5.3 W cm^−2^). Displayed is a zoom-in (with 103.17 nm pixels) of the full image shown in Fig. S3 over which 14,176,305 probes were detected (Table S2). (C) Zoom-in of B (10 nm pixels with 3-pixel Gaussian blurring). (D) Peak-aligned/-normalized average transverse profile for *n* = 10 single microtubule filaments, with each individual filament measured over a straight segment of ∼3 µm length (calculated from a super-resolved image with 5 nm pixels). Error bars represent ± SD. The black curve shows the Gaussian fit to the mean profile with a FWHM of 54.6 nm. (E) Zoom-in of boxed region in C with events displayed as peak-normalized 2D Gaussians color-coded for their localization precision according to the colorbar. (F) Zoom-in of E with color-coded crosses depicting the 1-*σ* error bars (see colorbar in E). White arrow indicates a localization at Ångström-level precision. (G) 2D histogram representation of localization precision (*σ*_*x*_) versus photons per event for the full dataset (see Fig. S10 for separate depictions of the photon count and localization precision distributions).

For probe localization and reconstruction of the final super-resolution image we developed and validated a general analysis pipeline made available here called PROSPERO, which stands for <u>P</u>recise <u>R</u>esolution <u>O</u>f <u>S</u>parse <u>P</u>robe <u>E</u>vents via <u>R</u>egistration & <u>O</u>ptimization (see Figs. S5–S9; Materials and Methods). PROSPERO first detects events in a small detection window (e.g. 9×9 pixel) by local thresholding. Every thresholded event in each frame is then fit to a model consisting of a 2D Gaussian plus flat background. Unique events are identified and tracked across multiple frames. Sub-nm drift correction is then performed using both the tracked probes and the brightfield images as fiducial markers (“Registration” step). Accounting for drift, each unique event is fit anew, now in a joint manner over its relevant frames, to a 2D Gaussian with globally fixed position (relative to the coordinate system defined by the first frame) with variable normalization for each frame; a variable flat background for each frame is simultaneously fit as well (“Optimization” step). The super-resolution information for each event over the full image, or over a rectangular subregion, can then be represented in various ways as shown below (e.g. as 5 nm pixel representation for the full image, or as color-coded 2D Gaussians or 2D error bars representing localization precision for a subregion).

To assess the obtained resolution, transverse intensity profiles for ten isolated microtubules from the full image (Fig. S3) were measured, with the mean profile displayed in Fig. 2D. Gaussian fitting yielded a full width at half maximum (FWHM) of 54.6 nm, which compared well with the microtubule width of 25 nm plus two flanking antibodies of size ∼10 nm (*27*) each labeled with a single APC of radius 4 nm, or 25 + 20 + 8 = 53 nm.

A detailed analysis using zoomed-in views of an isolated filament (Fig. 2E and 2F) depict the localizations of each antibody probe as 2D Gaussians or error bars, respectively. Collected photons per localization ranged from 10^3^‒10^6^ (Fig. S10A), with 65% of the localizations at <10 nm, 35% at <5 nm, and 3% at <1 nm precision (Fig. S10B). The apparent photobleaching half-time of the probes *in situ* was a few seconds (Fig. S10C), which was shorter than the 25 s measured for the analyzed subset of probes on glass, but still significantly longer than the 0.65 s transit time of the probes over the illuminated FOV (Materials and Methods). The observed uniform staining of the entire FOV (Fig. S3) proved the adequacy of these experimental settings. Furthermore, we analyzed the deviation of the localization precision distribution from the theoretical limit (Fig. 2G; dashed line), which we found could be attributed to a significant background component (present already in the raw data, Table S2) that was necessary to include in our 2D Gaussian fitting of each probe event. Despite this background, the high photon yield obtained at low illumination intensity from the LED still enabled a localization precision below 10 nm for the majority of the detected probes, with a significant fraction detected at better than 2 nm precision.

Further examples demonstrating homogeneous FILM imaging of other important intracellular markers in HeLa cells based on APC-conjugated antibodies are provided as supplemental figures: detection of cytokeratin bundles (Fig. S11) and single filaments (Fig. S12) down to 50 nm and 20 nm widths, respectively; LAMP1-rich vesicles with 50–200 nm widths (Fig. S13); and TOM22 distributed relatively uniformly along the mitochondrial outer membrane (Fig. S14). For these datasets, additional filtering out of radially asymmetric events was performed to remove false events due to off-center detection of highly out-of-focus probes along their diffractive rings (Materials and Methods; Fig. S6; Table S2). This additional filtering was not necessary for tubulin, as most events lied close to the focal plane with no significant evidence of such false detections in the tracked probe data (e.g. see the zoomed-in view of the tracked images displayed in Fig. S5B).

We next analyzed the staining of immune cells with FILM. For these datasets, filtering of radially asymmetric events was also performed. Additionally, filtering out of weak, poorly resolved events was performed to remove sharp optical features of the cells that were backlit by the free probe background (see Fig. S8).

First, we attempted the staining of *α*-tubulin in T and B cell co-cultures under conditions favoring their interactions through the formation of immunological synapses (Fig. S15). In all cells, the microtubule network was concentrated at the cell cortex, as previously observed (e.g. (*28*)). However, several sets of sparse, linearly aligned localizations were also identified in the interior (Fig. S16A). Such linear tubulin “structures” were not discernible in the crowded cytoplasm of the HeLa cells. Close examination of a set of such isolated linear structures (Figs. 3A and S16B–I) revealed nearly all localizations to reside within superimposed strips of 15 nm width, consistent with the resolution limit posed by the ∼10–15 nm antibody probes. Further examples of straight tubulin structures, as well as some that appear curvy and branching, are additionally displayed in isolated cells in Figs. 3B and S17 and in immunological synapses in Fig. S18. We speculate that these structures could represent isolated tubulin protofilaments, which have however never previously been reported *in vivo*, or tubulin monomers recruited to other narrow linear cytoskeletal elements.

**Fig. 3.**
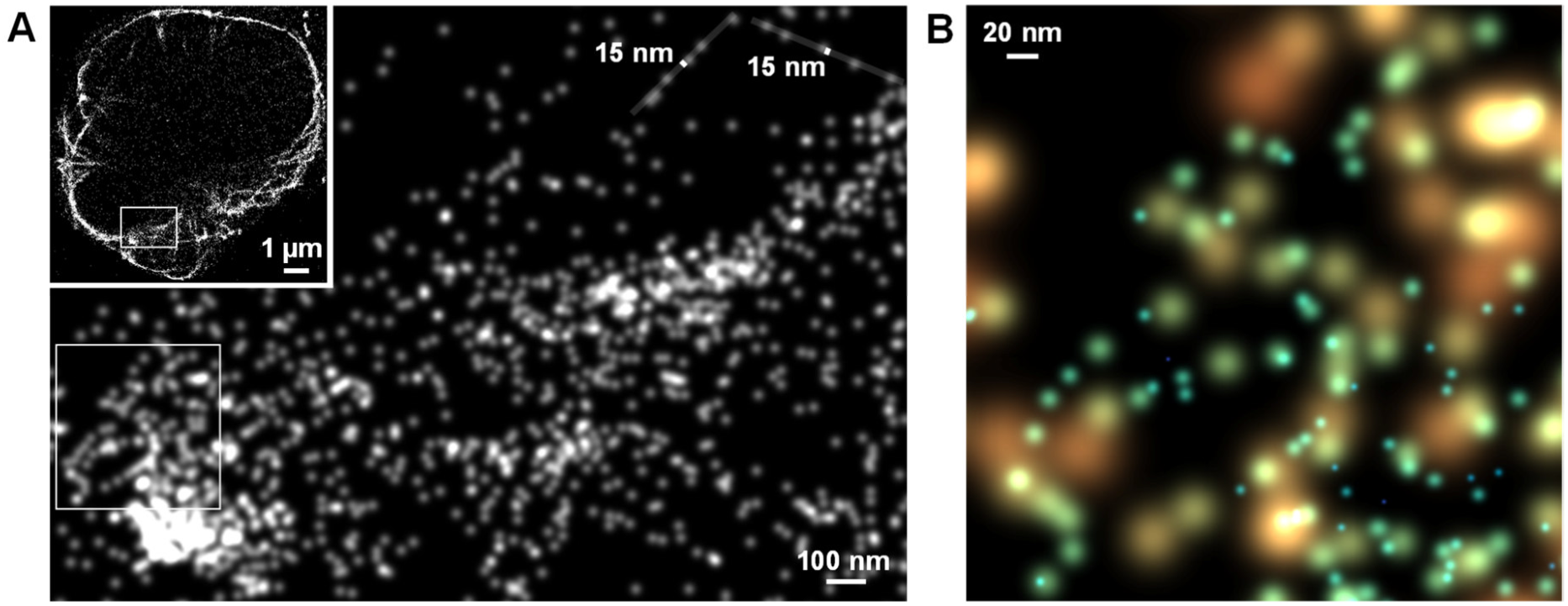
α-tubulin resolved in a T cell with FILM. (A) FILM image of the anti-α-tubulin antibody from Fig. 2 applied now to a PFA-fixed and permeabilized Jurkat T cell (zoom-in of the respective ROI indicated in Fig. S15). Lower right panel is a zoom-in of the boxed region in the upper left panel (5 nm pixels with 2-pixel Gaussian blurring). Two isolated linear tubulin structures with localizations confined to the displayed 15-nm overlaid strips are indicated at the upper right. (B) Zoom-in of the boxed region in the lower right panel of A (color-coded, peak-normalized 2D Gaussians as in Fig. 2E), showing multiple curvy tubulin structures with apparent widths smaller than the displayed 20 nm scalebar.

An important target that would greatly benefit from molecular-resolution imaging is the immunological synapse (*29*), a compact, multi-zonal structure of a few micrometer diameter that is composed of hundreds of distinct proteins, including receptor–ligand pairs that bridge the narrow membrane gap found at the central synapse (down to ∼10 nm) as well as the larger ectodomain spans characteristic of more peripheral regions of the synapse (up to ∼45 nm) (*30*) (Fig. S19). Immunological synapses play central roles in immune cell communication as well as in the killing of target cells by cytotoxic immune cells. Many aspects of their molecular-scale functional structures remain to be determined (*29*). We used APC-conjugated antibodies or a Vio^®^ 667-conjugated Fab fragment (ICAM-1; Materials and Methods) to resolve the substructural organization of the supramolecular activation cluster (SMAC) zones that constitute the “bullseye” structure of a T:B cell immunological synapse (Figs. S19). For the central SMAC (cSMAC) structure, tight central concentrations of TCRα/β, CD28, and the tetraspanin CD81 were found as expected (Figs. S20A–C). For the peripheral SMAC (pSMAC) zone, the markers CD2 and ICAM-1 were less represented at the center of the interface, consistent with exclusion from the cSMAC (Figs. S21A–B). ICAM-1 was more broadly distributed than CD2 across the cell-cell interface, likely reflecting its transition from early pSMAC enrichment to a more uniform distribution at this late stage (∼30 min) of synapse formation (*31*). The distal SMAC (dSMAC) zone marker CD45 exhibited a uniform distribution as well (Fig. S21C), which was also consistent with a late-stage synapse (*32*). Zoom-ins of the markers revealed intricate curvilinear substructures (Figs. S20D–F and S21D–F) consistent with membrane subdomains or cytoskeletal trafficking (e.g. CD2 in Fig. S21), and close (<20 nm) protein-protein associations (Figs. S20G–I and S21G–I). These results provide an important set of validated markers for deeper molecular co-localization studies of immunological synapse structures with FILM.

With these experiments, we have established FILM as an SMLM method that achieves *in situ* molecular resolution. Continuously flowing a probe at low concentration over a sample yields uniform, efficient localization across the full field of view. Gentle illumination — required to limit photobleaching of the free probe — also maximizes signal from bound targets, enabling detection of up to 10^6^ photons per molecule. This photon budget supports molecular-resolution reconstructions in biological specimens observed with widefield microscopy despite substantial background from the free probe.

### FILM Imaging of Multiple Markers on Tissue Slice Sections

With FILM, application of distinct probes over a single sample in a sequential manner should permit the analysis of histological sections. To this end, we serially applied FILM for multiple markers in two different tissue sections. As for the immune cell datasets, filtering out of radially asymmetric events (to remove off-center detections) and weak, poorly resolved localizations (to remove backlit features) was employed (Fig. S6; Fig. S8; Table S2; Materials and Methods). While localization precision was degraded somewhat due to high free probe background, a slight majority of the filtered events was still detected at <10 nm precision (compared to 65% of the events for tubulin in HeLa cells, Fig. 2G).

We first analyzed the intrafollicular region (IFR) of healthy tonsil tissue, where B cells are tightly packed in defined histological structures termed germinal center (GC) and mantle zone (MZ) (Fig. 4A). In cell culture, prior two-color super-resolution imaging revealed that the B cell receptor (BCR) isotypes IgM and IgD are located in distinct nanoclusters (*33*, *34*), organized by nanoclustered tetraspanins including CD81 (*35*). We stained these surface receptors and the BCR co-receptor and rituximab target CD20 using probes conjugated to either APC or phycoerythrin (PE). The super-resolution image that we obtained with FILM of the boxed region in Fig. 4A is displayed as a multicolor overlay in Fig. 4B, with comparison to standard IF on the left (Materials and Methods); the individual images comprising the overlays are additionally provided in Fig. S22. The respective ROI within the MZ is successively zoomed-in on in Fig. 4C–E, demonstrating the power of FILM for *in situ* colocalization analysis at molecular resolution.

**Fig. 4.**
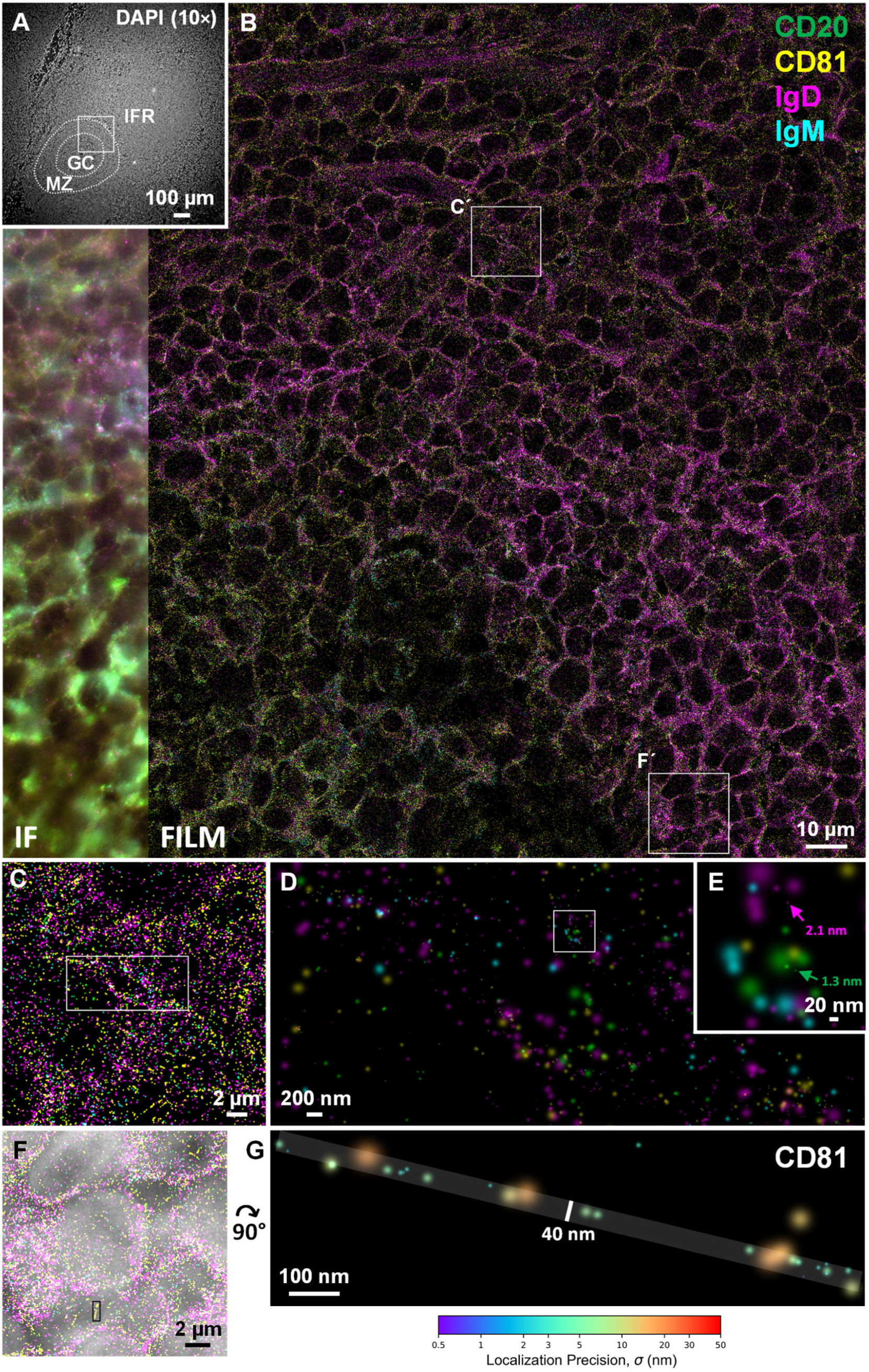
Serial FILM imaging of four markers in a tonsil tissue slice. (A) Overview DAPI image (using a 10× objective) of a PFA-fixed and quenched human palatine tonsil tissue section (4 µm thick) containing a follicle. White borders (dotted lines) are used to indicate the anatomic follicular zones corresponding to the germinal center (GC), mantle zone (MZ) and interfollicular region (IFR). (B) Multicolor overlay of sequential FILM acquisitions of the boxed region in A of PE-labeled IgM at 20% LED power (0.7 W cm^−2^) and APC-labeled IgD, CD81, and CD20 at 100% LED power (5.3 W cm^−2^) (10-nm pixels with 3-pixel Gaussian blurring). The sample was washed extensively with PBS under continuous flow and photobleaching before application of each successive probe. After FILM imaging, IF images were acquired (shown on the left side of the panel for comparison). (C) Zoom-in of the boxed region with Ć label in B (10-nm pixels with 3-pixel Gaussian blurring). (D) Zoom-in of boxed region in C using peak-normalized Gaussian representations for the localizations (color-coded for each marker as in B). (E) Zoom-in of boxed region in D. Colored arrows indicate individual localizations with 2.1 nm and 1.3 nm precision for IgD and CD20, respectively. (F) Zoom-in of the boxed region with F’ label in B (10-nm pixels with 3-pixel Gaussian blurring) overlaid with a conventional DAPI image (grayscale). Boxed region contains a single isolated linear CD81 structure. (G) Zoom-in of the boxed region in F, now displaying only the CD81 localizations as peak-normalized Gaussians with color-coded precision according to the colorbar. The estimated positions (centers of the Gaussians) for all 27 individual localizations lie within the overlaid strip of width 40 nm (two offset localizations toward the upper right were not considered as part of the structure). See Fig. S23 for four additional examples of similar linear CD81 structures.

Our visualization of 4 μm thick tissue sections supports the presence of homotypic protein clusters of varying sizes, especially for CD81 and IgD, which is much higher expressed in MZ B cells than IgM, while GC B cells largely lose IgD and IgM expression. CD81 imaging, in particular, revealed several narrow linear distributions scattered over the field of view of widths <40–50 nm (Figs. 4F–G and S23). These linear distributions correlated with regions of low nuclear staining, suggesting a peripheral localization. These distributions might therefore reflect the known localization of CD81 to peripheral, actin-rich, high membrane curvature structures (*36*), such as microvilli and connecting ridges on B cells (*37*).

Staining of a 4 µm thick section of colorectal cancer tissue with anti-pan-Cytokeratin-APC (Figs. S24) revealed ∼50–100 nm wide cytokeratin bundles (Fig. S25A–D) and narrower branching structures (Fig. S25E–G) consistent with single filaments. Successive FILM staining of the same tissue slice with anti-*β*-catenin-APC (Fig. S26) revealed cytokeratin-deficient, ∼200 nm wide “channels” of *β*-catenin flanked by cytokeratin (Fig. S27). A possible explanation for these structures could be the localization of *β*-catenin to other segregated parts of the cytoskeleton (e.g. its recruitment to microtubules by adenomatous polyposis coli (*38*)). Future co-localization studies using FILM of other cytoskeletal markers and well-characterized *β*-catenin interactors would help to better understand the origin of these structures.

Together, these examples display the power of FILM for the characterization of cellular features and close protein associations below the resolution limit within complex tissue samples.

### FILM Imaging of Single Actin Filaments Down to Ångström Resolution

Although FILM’s intrinsic resolving power is very high (see Fig. 2G), the *in situ* resolution achieved so far has been capped by the ∼10–15 nm size of the antibody labels, with linkage error from the probes broadening apparent features and reducing the effective resolution. To probe cellular structures with higher resolution, we turned to dye-labeled phalloidin of size ∼3 nm (*39*, *40*) to localize F-actin in fixed HeLa cells (Fig. 5A). For this experiment, we restricted the illuminated FOV, which reduced the general background and allowed for roughly two-fold higher resolution (Fig. S28). Inspection of the image by zooming into specific regions revealed an intricate actin network of thick bundles down to single filaments (Fig. S29). By focusing on regions with single peripheral actin bundles, a thickness of <45 nm for a single actin bundle could be observed (Fig. S30), which agreed well with the down to ∼50 nm resolution of actin bundles achieved with “Phalloidin-PAINT” (*41*). In another zoom-in shown in Fig. 5B many aligned localizations consistent with single actin filaments were observed. Further zoom-ins (Fig. 5C–F, Fig. S31) reveal that all localizations (centers of the Gaussians) resided within overlaid strips of 4 nm in width, demonstrating *in situ* attainment of <4 nm resolution, limited by the expected ∼3 nm wide helical arrangement expected for the dye when bound via phalloidin to the actin filament (*39*, *40*). For this background-reduced experiment, 50% of the probes were now localized with <2 nm precision and 20% with <1 nm precision (Fig. 5G, Fig. S28E–F). In Fig. 5H, a zoom-in of the boxed region in Fig. S29 is displayed. This region contains a highly resolved single filament (boxed region), which is shown in Fig. 5I to contain two highly precise localizations (at ∼3–4 Ångström precision) observed at 4.3 nm displacement from each other with 9 Ångström resolution (Materials and Methods). These results demonstrate the ability of FILM to capture substructures at <4 nm resolution over a large field of view.

**Fig. 5.**
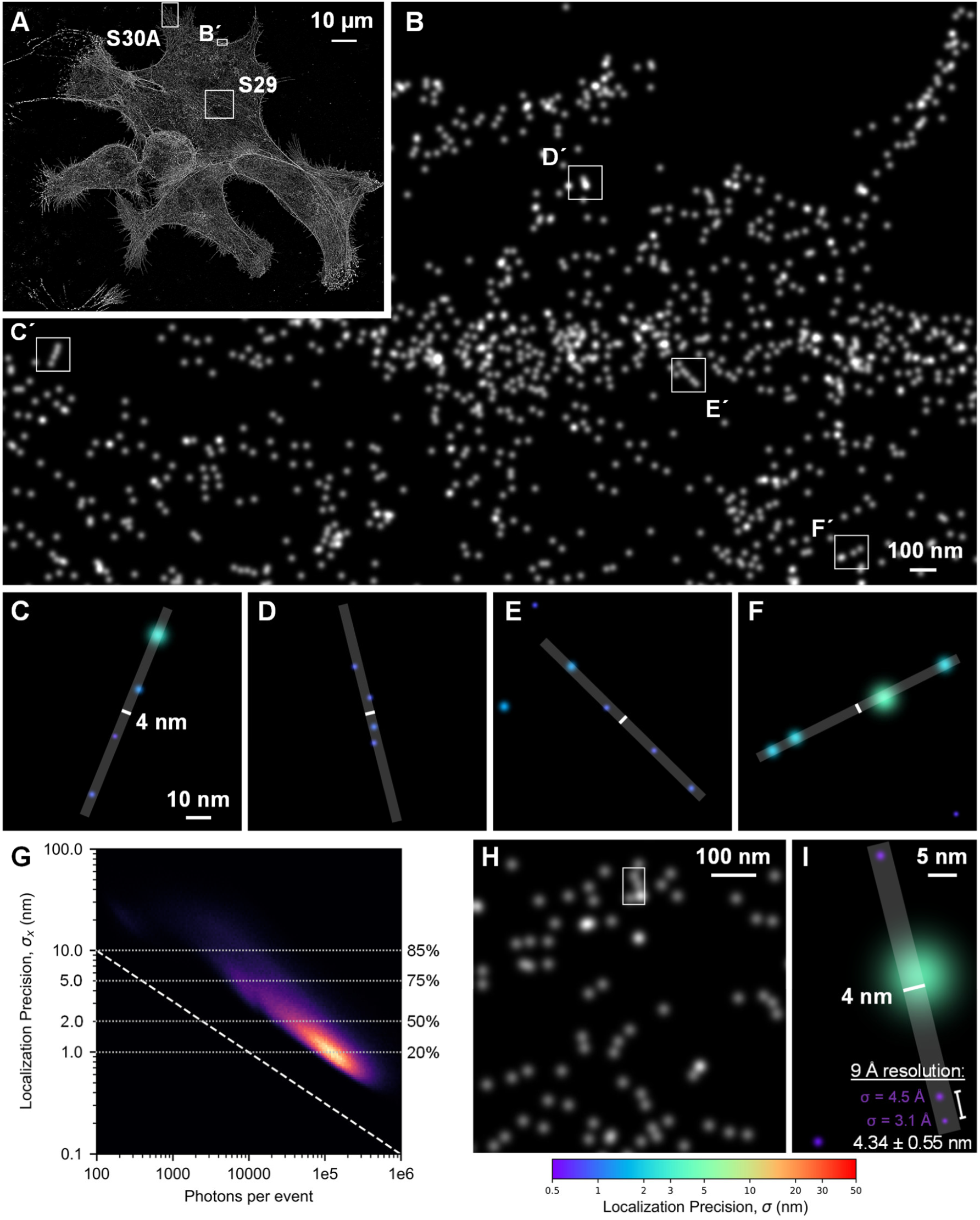
Phalloidin-Atto643 staining of actin in HeLa cells with FILM. (A) Background-reduced FILM imaging of the actin cytoskeleton in fixed HeLa cells using phalloidin-Atto643 and 100% LED (5.4 W cm^−2^) (10 nm pixels with 3-pixel Gaussian blurring). Four-fold reduction of the general, homogeneous background was achieved by reducing the excitation pinhole to 20% (27% of the FOV), allowing for higher localization precision for the same number of photons (see Fig. S28). (B) Zoom-in of the boxed region B’ in panel A (5 nm pixels with 2-pixel Gaussian blurring). (C–F) Zoom-ins of the respective boxed regions in B containing putative single actin filaments. Localizations are represented as Gaussians with color-coded precision according to the colorbar. All localizations for each single actin filament were found to reside within the overlaid 4-nm wide strips. (G) 2D distribution of localization precision vs. photons per event for the full dataset from A (Fig. S28E). Cumulative distribution values on the right axis give the percentage of events (see Fig. S28F) with precision less than the corresponding values on the left axis. A majority of events were detected at <2 nm precision and 20% at <1 nm (Ångstrom microscopy). (H) Zoom-in of boxed region in Fig. S29. (I) Zoom-in of boxed region in H. The two probes at the lower right are observed to be separated by 4.34 ± 0.55 nm, corresponding to 9 Ångstrom resolution (Materials and Methods). Refer to Figs. S29–31 for enlargements of the other respective boxed regions as well as other regions in A.

Motivated by the substantial increase in localization precision upon background reduction, we sought to determine the exact origin of the background. Surprisingly, control measurements revealed that the predominant background contribution in our phalloidin experiments came from the composite glass used in our microscope setup and *not* from the free probes (Fig. S32; Table S2; Materials and Methods). These results suggest the potentially significant benefit of the use of autofluorescence-free quartz (fused silica) glass for both the coverslip (*8*) and the objective (e.g. 100×/1.35 N.A. Ultrafluar objective, previously available from Carl Zeiss).

## Discussion

Taken together, our results demonstrate a novel and highly accessible approach to single molecule localization microscopy (SMLM), termed Flow Immunofluorescence Localization Microscopy (FILM), which leverages standard widefield microscopy systems and off-the-shelf fluorescent probes to achieve spatial resolution of the probe dyes down to the Ångström scale across the field of view of a widefield fluorescence microscope. FILM supports high-density, serial multiplexed imaging of multiple molecular targets within a single specimen, offering a uniquely powerful platform for spatial molecular biology. Using this method, we reveal several previously uncharacterized nanoscale features in immune cells, within immunological synapses, in human tonsil tissue, and in colorectal cancer samples. Moreover, imaging of individual actin filaments labeled with phalloidin-Atto643 demonstrated <4 nm resolution *in situ*, with closely spaced localization events imaged down to 9 Ångström resolution.

While FILM can already be applied using conventional antibody probes, full exploitation of its molecular resolution requires nm-scale probes (*42*), such as phalloidin or antibody fragments.

FILM is compatible with stable as well as semi-stable probes with binding half-times down to the minute timescale (e.g. lower affinity antibody Fab fragments). Stable binders can even be made semi-stable by applying a chaotropic agent (*43*). The use of semi-stable probes, in particular, has the advantage of much lower label accumulation on the sample over time, minimizing steric hindrance and artefactual sample expansion (*44*).

Here, we have demonstrated FILM based on the non-covalent binding of probes to targets; however, gradual *covalent* labeling of tagged targets with organic dyes flowed at low concentration over the sample should also be possible, for example, for FILM-based versions of SNAP-tag or HaloTag labeling (*45*).

While photon collection from each individual probe molecule is slow with FILM, simultaneous camera-based acquisition of hundreds to thousands of probes across the entire field of view makes it a few times faster than the sequential detection of single probes with MINFLUX (Materials and Methods). We have imaged several challenging, highly abundant cytoskeletal markers to few percent labeling efficiency, amounting to e.g. ∼250,000 localizations of F-actin per HeLa cell over the course of ∼8 hours of image acquisition (Fig. 5A; Materials and Methods; Table S2). For more typically expressed protein targets, having only tens to hundreds of thousands of epitopes per cell, FILM can therefore localize most of the targeted epitopes on a similar or shorter timescale to molecular resolution. For even faster acquisition with FILM, simultaneous acquisition of multiple markers using spectral multiplexing could be employed (*46*, *47*).

There is significant room for improvement of the attainable resolution with FILM. In addition to the above mentioned use of quartz glass for the optics (Fig. S32), resolution could additionally be improved by use of a lower profile flow chamber (e.g. 50 µm instead of 540 µm) to reduce the free probe background and its artefacts (Fig. S8) and by use of a photostabilizing buffer to increase photon collection (e.g. to >10^7^ photons per fluorophore (*23*)).

We have exclusively explored *lateral* resolution in the above; however, FILM’s superb photon collection efficiency allows for robust detection of the PSF shape — even for highly out-of-focus probes displaced out to 5 µm from the focal plane and displaying up to five diffractive rings (Fig. S33), and should therefore provide an excellent basis for axial localization, e.g. using biplane detection (*48*), to enable fully 3D molecular, and even *volumetric*, SMLM. This would, for example, be extremely useful for 3D exploration of immunological synapse substructures and marker co-localization using the probes validated above.

In summary, FILM is a widely accessible, high-resolution imaging technique that achieves down to Ångström-scale localization using standard microscopy and probes, enabling dense, multiplexed molecular mapping in cells and tissues for advanced spatial biology.

## Materials and Methods

### Sample preparation

#### Coating of slides

To ensure later cell attachment, the glass surface of the µ-Slide VI 0.5 (ibidi, #80607, 540 µm channel height) was coated in advance. After a quick washing step in sterile ddH_2_O, each channel was filled with 0.01% Poly-ʟ-Lysine solution (PLL, Sigma-Aldrich, #P4832) and incubated for 1 h at room temperature. The channels were then thoroughly washed thrice with sterile water before cell seeding. For all tissue samples, #1.5H glass coverslips (ibidi, #10812) were thoroughly cleaned and then silanized. Specifically, each coverslip was successively submerged for 30 s in 36% hydrochloric acid (HCl), distilled water, and isopropanol, and then dried at room temperature. The coverslips were then placed in a freshly prepared silane treatment solution within a glass Petri dish and incubated for 1–2 h at room temperature. After incubation, each coverslip was individually rinsed with ddH_2_O and washed twice for 10 min each in a glass Petri dish submerged in ddH_2_O. The coverslips were then dried overnight at 60 °C in a drying cabinet and stored for up to several weeks at 4 °C.

#### Fixation

Cells were washed once with PBS and then fixed with either pre-warmed 4% (vol/vol) paraformaldehyde (PFA) in PBS or cytoskeleton-preserving buffer (CB; 3.7% (wt/vol) PFA, 0.5% (vol/vol) Triton X-100, 10 mM MES (pH 6.1), 90 mM KCl, 3 mM MgCl_2_, 2 mM EGTA) for 15 min at room temperature. Following fixation, cells were washed four times with PBS (30 s, 1 min, 5 min, 10 min). Slides were stored short-term in storage solution (0.02% sodium azide (Sigma-Aldrich, #S8032) in PBS, pH 7.4) at 4 °C in a humidified chamber.

#### Permeabilization

When permeabilization was necessary to access the target epitope within the cell, the sample was treated with 0.1% (vol/vol) Triton X-100 (Sigma-Aldrich, #T8787) in PBS for 15 minutes at room temperature.

#### Blocking

Prior to each imaging session, samples were blocked with 2% (wt/vol) bovine serum albumin (BSA) (Carl Roth, #3737) in PBS for 30 min at room temperature to saturate excess binding sites. Subsequently, the slide was rinsed several times with PBS.

#### Mammalian cells

Human cervical cancer cells (HeLa, American Type Culture Collection, #CCL-2) were cultured in Dulbecco’s Modified Eagle’s medium (DMEM) (Biowest, #L0091) containing 10% (vol/vol) FCS and 2 mM L-glutamine. As a model for the immunological synapse, human B cell lymphoma Raji cells (American Type Culture Collection, #CCL-86) and Jurkat cells (clone E6-1, American Type Culture Collection, #TIB-152) were cultured in RPMI-1640 medium (Biowest, #L0501) supplemented with 10% (vol/vol) fetal bovine serum (Eximus Group, #BS-2022-500) and 2 mM L-glutamine (Thermo Fisher Scientific, #11539876). Cells were passaged every 2–3 days, adherent cells were detached using trypsin-EDTA (Thermo Fisher Scientific, #15400054), while suspension cells were diluted. All cells were maintained at 37 °C in a 5% CO_2_ atmosphere. For cell number and viability assessment, single cell suspensions were diluted 1:1 with 2% (wt/vol) Erythrosin B solution (Sigma-Aldrich, #1159360010) and cells were counted on a Neubauer hematocytometer. HeLa cells were seeded onto freshly PLL-coated µ-Slide VI 0.5 at a density of 10,000 cells per channel and incubated for at least 48 h at 37 °C with 5% CO_2_ to promote cell adherence and spreading before fixation.

#### Immune cells and immunological synapses

For later distinguishment, Raji cells were pre-incubated with CellTracker^®^ Blue CMAC (7-amino-4-chloromethylcoumarin, Thermo Fisher Scientific, #C2110) (10^6^ cells mL^−1^) at 10 µM in Phosphate Buffered Saline (PBS) (pH 7.4) (Sigma-Aldrich, #D8537) for 15 min at 37 °C. After two rinsing steps in complete medium, Raji cells were resuspended at a concentration of 3.3×10^5^ cells mL^−1^. To each channel of the freshly coated µ-Slide VI 0.5, 30 µL of the stained cell suspension was applied (equaling 10,000 cells) by addition of the solution into the connected reservoir well of each channel and removal from the associated well on the other side. This step was repeated three times to ensure that the liquid was exchanged complete. The chamber slide was incubated at 37 °C and 5% CO_2_ for 30-45 min to allow the labeled Raji cells to adhere to the glass bottom. Subsequently, Raji cells were pulsed with 2 µg/mL Staphylococcal enterotoxin E (SEE) (Hölzel Diagnostika Handels GmbH, #CSB-EP320170FKZ) in conjugation medium (CM = RPMI-1640 without phenol red (Biowest, #L0505), 0.5% (vol/vol) FCS, 2 mM L-glutamine) for 30 min. Proper liquid exchange within the channel was ensured using the addition/removal of solution steps described above. Without washing, Jurkat cells were co-seeded onto the slide in a 1:2 ratio (6.6 × 10^5^ cells mL^−1^, 20,000 cells) and incubated for another 40–45 min at 37 °C, 5% CO_2_ to allow synapse formation. At this cell density, cells were sufficiently spaced from each other (20% confluency) for microscopy.

#### Tissue slices

Fresh frozen tissue (ProteoGenex, Sample ID #68330B2(2) tonsil, #66747T2(1) colorectal adenocarcinoma) was sectioned at 4 μm slices and mounted onto silanized coverslips, specifically in the area where the small channel (5 mm × 48.2 mm) was subsequently positioned. Immediately following fixation with PFA or CB and a brief rinsing period, the sample undergoes a 1-hour quenching step in 100 mM glycine at 4 °C, followed by a 30-minute wash in PBS. Once the protective film was removed from the adhesive tape, the sticky-Slide I Luer (ibidi, #80178, 400 µm channel height) was carefully affixed to the bottom glass coverslip holding the fixed sample, applying slight pressure. PBS was then directly added to fill the channel. Slides were stored short-term in storage solution (0.02% sodium azide in PBS, pH 7.4) at 4 °C in a humidified chamber.

### Labeling reagents

#### Commercial antibodies and reagents for super-resolution microscopy and IF staining

α-tubulin (REA1136 REAfinity™, h, APC, #130-119-541, Miltenyi Biotec), CD2 (REA972 REAfinity™, h, APC, #130-116-150, Miltenyi Biotec), CD20 (REA780 REAfinity™, h, APC, #130-111-339, Miltenyi Biotec), CD28 (REA612 REAfinity™, h, APC, #130-118-343, Miltenyi Biotec), CD45 (REA1023 REAfinity™, h/nhp, APC, #130-117-191, Miltenyi Biotec), CD54/ICAM-1 (REA266 REAfinity™, h, APC, #130-120-711, Miltenyi Biotec), CD81 (REA513 REAfinity™, h, APC, #130-119-787, Miltenyi Biotec), CD107a/LAMP1 (REA792 REAfinity™, h, APC, #130-111-847, Miltenyi Biotec), IgD (REA740 REAfinity™, h, APC, #130-110-644, Miltenyi Biotec), IgM (REAL689 REAdye_lease™, h, PE, #130-125-786, Miltenyi Biotec), Pan-Cytokeratin (REA1141 REAfinity™, h, APC, #130-120-096, Miltenyi Biotec), β-catenin (REA480 REAfinity™, h, APC, #130-124-444, Miltenyi Biotec), TCRα/β (REA652 REAfinity™, h, APC, #130-113-535, Miltenyi Biotec), TOM22 (REA1185 REAfinity™, APC, h/ms, #130-122-079, Miltenyi Biotec), DAPI Staining Solution (#130-111-570, Miltenyi Biotec), and ATTO 643 phalloidin (ATT-AD643-81, ATTO-TEC). All commercially available antibodies have been validated by their manufacturer as indicated in their respective datasheet and/or website. (h = human, ms = mouse, nhp = non-human primate)

#### Fab conjugation using NHS-activated dyes for super-resolution microscopy

The lyophilized dye (Vio^®^ 667) were allowed to equilibrate to room temperature before being dissolved in dry DMSO (Thermo Fisher, #D12345) to a concentration of 20 mg mL^−1^. The concentrations of the dye and unconjugated CD54/ICAM-1-Fab (REAL146, REAlease^®^, h) or α-tubulin-Fab (REAL979, REAdye_lease™, h) Antibody were measured immediately before conjugation using absorbance readings at 635 nm and 280 nm, respectively, on a NanoPhotometer^®^ NP80 (Thermo Fisher). The coupling reaction was conducted at a dye-to-protein ratio of 3 to achieve a degree of labeling (DOL) of approximately 1.5. In brief, 15 µmol of Fab was diluted to 2.5 mg mL^−1^ in CliniMACS^®^ PBS/EDTA buffer (Miltenyi Biotec, #200-070-025) supplemented with 50 mM sodium carbonate buffer (pH 9). A 3-fold molar excess of NHS-ester-coupled dye was added to the reaction mixture and incubated for 1 h at room temperature with gentle mixing (800 r.p.m.). Excess unreacted dye was removed by size exclusion chromatography using either Zeba™ Dye Removal Spin columns (Thermo Fisher, #A44299; MWCO 7 kDa) or Amicon filters (Merck/EMD Millipore, #UFC501024; MWCO 10 kDa), following the manufacturer’s instructions. After purification, the conjugate was diluted 1:100 with 100× Storage solution to reach a final concentration of 0.1% (wt/vol) of human serum albumin (HSA) (Octapharma, #5401000) and 5 % (wt/vol) of sodium azide in PBS (pH 7.4). The DOL was calculated using absorbance measurements at 635 nm and 280 nm, taking into account the dye’s correction factor using the formula:

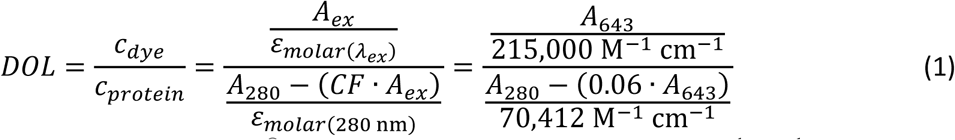

with *CF* the correction factor for Vio^®^ 667 absorption at 280 nm, 215,000 M^−1^ cm^−1^ the molecular extinction coefficient of Vio^®^ 667 at 643 nm, and 70,412 M^−1^ cm^−1^ the molecular extinction coefficient of the Fab conjugate at 280 nm. The purity of the conjugates was confirmed by analytical size-exclusion chromatography, using pure Fab and Vio dye as controls. The Vio^®^ 667-coupled CD54/ICAM-1-Fab displayed an average DOL of 1.5.

### Experimental setup and data acquisition

#### Microscope

Conventional IF imaging and SMLM measurements were performed on a custom widefield setup built on a Zeiss AxioObserver inverted microscope (Carl Zeiss) equipped with a 63× oil immersion objective (63×/1.46 *α* Plan-Apochromat Oil Corr TIRF, Carl Zeiss). For control measurements (Fig. S32), a 63× water immersion objective (63×/1.2 C-apochromat W Corr, Carl Zeiss) was also employed. Fluorescence excitation was achieved using a set of LEDs (Colibri 7, Carl Zeiss) emitting at 631/33 nm, 555/30 nm, 469/38 nm, and 385/30 nm. The size of the illuminated FOV could be controlled by adjusting a pinhole for the excitation light. Samples were mounted on 170 µm thick borosilicate coverslip glass (#1.5H glass coverslips, ibidi, #10812). A quartz (fused silica) coverslip of 150–180 µm thickness (Science Services, #E72256-08) was used for control measurements (Fig. S32). Images were captured by a monochromatic scientific CMOS (sCMOS) camera (Prime BSI, Photometrics) with a resolution of 103.1738 nm px^−1^ for the 63× objective. All excitation, dichroic, and emission filter sets defining the epifluorescence channels that were optimized for the dyes are detailed in Table S1. The setup was mounted on a vibration-isolated optical table. Axial drift was corrected for using a hardware autofocus (Definite Focus.2, Carl Zeiss), which measures the position of the glass coverslip by reflecting weak infrared (∼850 nm) LED light off of it, and then adjusts, if necessary, back to the user-defined focal position in the sample. Lateral drift was later corrected during image analysis by tracking of the ensemble of probes with PROSPERO (see below).

#### Illumination intensity at the focal plane

For the assessment of photon counts under different intensities shown in Fig. S2, a 631/33 nm LED (Colibri7, Carl Zeiss) and a 400 mW 640 nm laser (Rapp OptoElectronic) were employed, with corresponding filter combinations provided in Table S1. In order to measure the applied power density at the sample, we measured the laser and LED illumination light by placing a power meter directly on top of the objective using a restricted excitation pinhole to create an illuminated FOV smaller than the camera chip to measure the area of illumination for precise determination of the intensity (measured power divided by illumination area) at the sample plane.

#### Microfluidic handling

Liquid handling was controlled using an ARIA fluidics device (Fluigent, #CB-SY-AR-01), which is equipped with 10 connected reservoirs for precise pressure-based delivery of distinct liquid solutions to the sample. All solutions introduced into the system were first passed through a 0.22 μm syringe filter. The selected solution was then directed from its designated ARIA reservoir through the internal PTFE tubing (1/16” OD + 0.010’’ ID), and subsequently through a compatible tubing adapter (Darwin Microfluidics, #CIL-P-757) into silicone tubing (3.2 mm OD + 1.6 mm ID silicone; ibidi, #10842). Following a tubing prefilling step with PBS, the inlet and outlet silicone tubing were securely attached to the µ-Slide via an Elbow Male Luer Connector (ibidi, #108025), while carefully avoiding the introduction of air bubbles. The inlet of the chip receives liquid from the selected ARIA reservoir, with the outlet directing the flow into a waste bottle. Proper cleaning of the tubing was carried out after each imaging session or when switching probes, following the manufacturer’s instructions. The sample chamber (Fig. 1A) used for the cultured cells corresponded to a µ-Slide VI 0.5 Glass Bottom (Ibidi) with a height of 0.54 mm, width of 3.8 mm and length of 17 mm. For the tissue slices, the sticky-Slide I Luer from Ibidi had a height of 0.4 mm, width of 5 mm and length of 50 mm. The antibody probe was flowed through the sample chamber at a continuous flow rate of 25–100 μL min^−1^ using the microfluidic pump of the ARIA. The flow velocity within the chamber was then *V* = 203‒ 812 µm s^−1^ for both slides due to their similar cross-sections. The length, *L*, of the field of view was 211.3 µm, implying a transit time over the full extent of the field of view of *T*_transit_ = *L*⁄*V* = 0.26 − 0.96 s. After some hours, bubbles often formed in the tubing leading up to the sample chamber, which upon their transport to the sample chamber, interrupted the acquisition of several early runs, including datasets shown here. Insertion of a degasser (IDEX, #0001-6682), directly downstream of the outlet from the Aria device, along with a bubble trap (Darwin Microfluidics, #LVF-KBT-S-A), directly upstream of the sample chamber, reduced bubble formation substantially. Vibrations from the active degasser were prevented from reaching the sample by rigid attachment of the flexible tubing downstream of the degasser to the floating optical table. The degasser was e.g. employed during acquisition of the highest resolved dataset presented here in Fig. 5, which demonstrated down to <4 nm precision and thereby successful prevention of the transmission of vibrations from the degasser to the sample.

#### SMLM image acquisition

Prior to each FILM experiment, candidate probes were optimized for probe concentration to generate a high number of sparse events across the field of view for efficient SMLM acquisition. For the tissue samples, ROIs with interesting morphology were selected based on prior DAPI staining of the samples. Automated image acquisition was performed using the Experiment Designer module of the ZEN blue software as follows (Fig. S4):

Block 1: autofocusing and acquisition of a brightfield image (not used).

Block 2: delay of 5 s to allow the system to fully equilibrate following the autofocusing.

Block 3: acquisition of a brightfield image later used for registration (drift correction).

Block 4: acquisition of a 1-min fluorescence movie with frames of 5 s.

Imaging of APC and Vio^®^ 667 was performed at 100% LED (631/33 nm), with a power density of 5.3 W cm^−2^. For the PE conjugate depicted in Figure 6, 20% LED (555/30 nm) and a power density of 0.7 W cm^−2^ was utilized. All imaging was performed using the High-Dynamic-Range (HDR) Combined Gain 16-bit mode for the PrimeBSI camera (Photometrics). Both experiment blocks were repeated hundreds of times to collect roughly 2000–21000 frames (each 5 s). Acquisition specifics, including applied concentration, were adjusted for an optimal number of events in each frame in trial runs, which e.g. depended on the binder association rate with the target and target abundance on the sample. Additional conversion of the ZEN software *.czi files to the TIFF format required for PROSPERO were made using ImageJ (*49*). A comprehensive list of all samples, probes, and instrumental settings for each FILM acquisition is provided in Table S2.

#### IF staining

To confirm specificity of the FILM acquisitions, cells were labeled at a dilution of 1:50 with the corresponding primary antibody in autoMACS Running Buffer (Miltenyi Biotec, #130-091-221) at room temperature following FILM imaging. Samples were typically also stained with DAPI Staining Solution (#130-111-570, Miltenyi Biotec) at a dilution of 1:100.

Following the removal of the staining reagents, cells were washed three times with PBS before imaging.

### FILM analysis with PROSPERO

PROSPERO is a completely general, custom-written, Python-based analysis software for FILM datasets with a user-friendly GUI that only requires as input: (1) separate directories containing the brightfield images and fluorescence movies in TIFF format; (2) the relative acquisition times and exposure times for each image; and (3) specific camera values corresponding to the pixel size at the sample (103.1738 nm), readout offset (100 I.U.), gain (1.67 I. U. per e^−^), and, optionally, the readout noise (1 e^−^) and dark current (0.5 e^−^ per s). The camera values in parentheses were the specific ones pertaining to our device (PrimeBSI, Photometrics). Optional parameters are the readout noise, which is typically insignificant compared to Poissonian errors for FILM, and a homogeneous dark current contribution, which is completely degenerate with the flat background model component used for event fitting and therefore plays no role. A user-chosen threshold is required before starting the full analysis; the user can test different threshold values for event detection for a specific frame from the movie. Once the optimal threshold has been specified (along with other optional parameter settings in the GUI), the full analysis consisting of following 13 steps can proceed:

1. Registration of the brightfield images is performed with the “Tanialign” subroutine from the Tanitracer software suite (*50*), which is based on the robust AKAZE algorithm for image alignment (*51*). A list of the relative shifts in *x* and *y* for each brightfield image relative to the very first brightfield image is thereby obtained.
2. (2) The original raw intensity image, image_1_, is first subtracted of the readout offset and the dark current to obtain image_2_. A locally determined background map for each frame is obtained by sliding a (2*n*+1)×(2*n*+1) window of radius *n* pixels (we used *n* = 4) over image_2_ and assigning the minimum value over the window to the position of the central pixel to obtain background_1_, which is then smoothed by averaging over a larger (2*m*+1)×(2*m*+1) pixel window of radius *m* pixels (we used *m* = 10) to create a final smoothed background map, background_2_. background_2_ is subtracted from image_2_ to obtain image_3_. A (2*n*+1)×(2*n*+1) window (*n* = 4) is then slide over image_3_ to determine the integrated intensity over the window for each window position divided by the propagated noise to obtain a signal-to-noise ratio (SNR); for positions of the window having an SNR that exceeds the user-defined threshold, the position of the local peak intensity value is recorded. Filtering of this list of peak intensity positions for *unique* entries generates a final list corresponding to peak-centered positions of all thresholded events in the image. A model consisting of a 2D Gaussian (pixel-integrated) plus flat background is fit within a (2*n*+1)×(2*n*+1) window of radius *n* pixels (with, again, *n* = 4) centered at each event position for the raw intensity image, image_1_, by nonlinear maximum likelihood (ML) estimation based on the Levenberg-Marquardt algorithm (*52*, *53*), with the uncertainty for each pixel intensity determined by the standard Gaussian approximation to the Poissonian photon shot noise along with the Gaussian camera noise model. Uncertainties in the parameters were determined in the standard way by approximating the likelihood function as a multidimensional Gaussian, defined by assessment of the curvature of the likelihood function (at its peak value) along each parameter axis (*54*). The position of radial symmetry is also calculated in this step over the (2*n*+1)×(2*n*+1) window of radius *n* pixels (with, again, *n* = 4) according to the method of (*55*), but with the weight terms only dependent on the squared intensity gradients and not additionally on inverse distance (see Eq. 1.4 in (*55*)).
3. Events are first filtered for a positive normalization of the 2D Gaussian fit and, optionally, for additional parameters having to do with other aspects of their fit values. For the latter, only filtering based on the difference of the positions of radial symmetry and the Gaussian center were used for the indicated datasets in Table S2, for which we used a difference in distances of no greater than the pixel dimension (1 pixel = 103.1738 nm). Starting from the first frame, events are assigned successive integer IDs. Cycling over the next *q* frames (we used *q* = 3), all events in each frame within a certain radius (we use a radius of 3 times the pixel size) of an event in the first frame are assigned the same ID, with all other novel events assigned successively higher IDs. The algorithm then steps to the second frame and repeats the process (assigning the same ID to any events in the next *q* frames that coincide with an event in the second frame). This is repeated for each frame until reaching the final frame.
4. A directory containing color-coded tracked images can optionally be created (see e.g. Fig. S5B).
5. For each unique localization ID, one can optionally filter out the rows pertaining to its first and last frames to remove potential temporal bias for the registration based on tracked particle information performed in the next step. (As this did not significantly improve the registration results, this was not performed for the analysis presented in this manuscript.)
6. For each pair of two successive frames, the relative displacement of the second with respect to the first frame is calculated by a weighted averaging of the shifts in fitted positions of all tracked events (typically hundreds to thousands). Proper statistical weighting is determined by error propagation of the uncertainties for the subtracted *x* (and, separately, *y*) positions based on the Gaussian uncertainties for these parameters obtained in step 2. In the end, a list of the relative *x* and *y* shifts for each frame with respect to the very first frame. Roughly speaking, assuming a typical localization precision of individual events based on single frame information of 20 nm, statistical combination of the uncertainties of >400 such tracked events between the two frames would imply an ensemble precision of 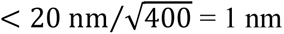 uncertainty for determination of the relative frame drift (drift correction at sub-nm precision).
7. “Registration” step: The brightfield image registration (defined at the times corresponding to the acquisition of each brightfield image, specifically, the middle timepoint for the exposure) from step 1 is combined with the tracked probe registration (again, defined at the middle timepoint for each fluorescence exposure) as follows. The brightfield images are considered to provide an absolute reference for the registration; however, their sampling is too crude, making linear interpolation for the registration information between successive brightfield images only valid to ∼10–20 nm accuracy: Compare the final estimate for the drift correction (cyan curves) with the brightfield registration estimate (black curves) in Fig. S7. The tracked probes provide better accuracy at shorter timescales (less than a few minutes), but display significant drift over few-minute-to-hour timescales (note gradual deviation of the magenta curves from the black curves in Fig. S7). We correct the tracked probe registration values (magenta curves) for this long-term drift by adding a spline correction (orange curves) that brings the linearly interpolated tracked probe registration (magenta curves) into perfect alignment with the brightfield registration (black curves) at the respective timepoints of the latter to create the final interpolated drift correction estimates of each frame for *x* and *y* (cyan curves).
8. For each tracked probe (unique integer ID), the estimates of its position can now be corrected for drift. The final position of the probe with respect to the initial image coordinate system can then be determined by a weighted average of its estimates in each frame, with proper statistical weighting corresponding to the relevant positional uncertainties obtained in step 2. Error propagation of the Gaussian positional uncertainties yields final “averaged” uncertainty values for the *x* and *y* positions of each event.
9. Events are optionally filtered for user-defined ranges of specific fit parameters. A super-resolution image is then created for a default (5 nm) or user-defined pixel size for the averaged positions from step 8. This allows for a “quick view” of the super-resolved image based on *averaged* localizations over multiple frames for each event from step 8. More rigorous joint analysis of the multiple frames for which a given probe appeared is carried out in the next step, corresponding to the principal result of PROSPERO.
10. “Optimization” step: Based on the tracking information from step 7, each tracked probe (unique integer ID) can now be jointly fit across each of its relevant frames using global parameters for the *x* and *y* positions and width of its 2D Gaussian, and local parameters for the Gaussian normalization and flat background level in each frame. Uncertainties in the parameters were determined in the standard way by approximating the likelihood function as a multidimensional Gaussian, based on assessment of the curvature of the likelihood function (at its peak value) along each parameter axis (*54*).
11. Events are optionally filtered for user-defined ranges of specific fit parameters (see Table S2). A super-resolution image is created for a default (5 nm) or user-defined pixel size for the fitted positions from step 10.
12. Several plots are created to display the analysis results. These include a histogram of photons collected per probe (e.g. Fig. S10A), a cumulative distribution for localization precision in *x* and *y* (e.g. Fig. S10B), a cumulative distribution for the probe lifetime (e.g. Fig. S10C), and a 2D histogram of localization precision vs. photons collected per probe (e.g. Fig. 2G).
13. Events are optionally filtered for user-defined ranges of specific fit parameters (see Table S2). Zoomed-in views of a user-defined rectangular ROI are then created, e.g. using peak-normalized (or integral normalized) 2D Gaussians (Fig. 2E) or error bars (Fig. 2F) to represent the localizations, and with and without color-coding for localization precision.

For most datasets, events with high radial asymmetry were excluded to remove off-center detections of diffractive rings of highly out-of-focus probes (Fig. S6). For the immune cell and tissue slice datasets, events with either low counts (e.g. <2000) or low localization precision (e.g. >20 nm) were excluded due to noticeable artefacts in these datasets (Fig. S8). See Table S2 for the specific filtering of events used for each dataset.

### Additional image analysis

#### Analysis of microtubule transverse profiles

Transverse profiles of microtubule filaments were obtained from reconstructed FILM images. Specifically, the intensity profile of 10 filament segments of 3 µm length was extracted from the full FOV (Fig. S3) by means of rotated rectangles in ImageJ (*49*) for a total analyzed length of 30 µm. The label intensity was normalized to unity, and the average line profile was fit with a Gaussian function (Fig. 2C). The FWHM from the Gaussian fit was compared to the theoretical outer diameter of ∼55 nm expected for an APC-labeled antibody-decorated filament (Fig. 2A).

#### General image preparation and analysis

ImageJ (*49*) was used for image exploration, basic image analysis (intensity), and image preparation for the main and supplemental figures (e.g. Gaussian blurring, image cropping and contrasting). For the 2- or 3-pixel Gaussian blurring mentioned throughout the figure captions, this corresponded to application of the ImageJ tool “Process/Filters/Gaussian blur…” with “Sigma (radius)” set to 2 or 3, respectively.

#### IF image alignment

The multi-color IF image shown on the left side of Fig. 4B was generated by aligning the individual images based on their concomitantly acquired brightfield images using the ImageJ (*49*) plugin Align_Slice (Landini, G. Code at https://github.com/landinig/IJ-Align_Slice).

### Photon collection and localization precision for SMLM

#### Theoretically achievable precision based on photon counts

The PSF of a microscope can be modeled as an Airy disc, which, for simplicity, is often modeled with a Gaussian function.

Fitting of an Airy disc with a Gaussian yields a FWHM (full width at half maximum) of size:

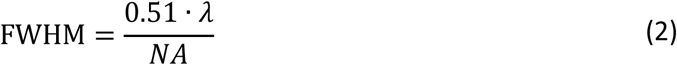

Converting to the standard deviation of the Gaussian (FWHM = 2.35*σ*_PSF_) gives:

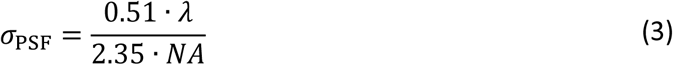

for the achieved precision. The limit on lateral localization precision, *σ*_SEM_, obtained for a single emitter depends on the standard error of the mean (SEM) of its separate photon distributions in *x* and *y*. For a single photon, the precision will at best equal the optical resolution of the microscope, or *σ* ≥ *σ*_PSF_. More generally, however, for *N* detected photons, this theoretical lower limit, corresponding to *σ*_SEM_, is reduced by the square root of *N*:

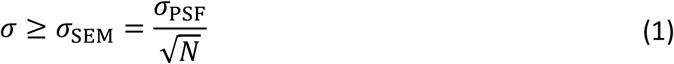

Rearranging for *N* and substituting in *NA* = 1.46 (for our oil immersion 63× oil objective, Materials and Methods), we find that the minimum number of photons needed to achieve a certain precision *σ* is:

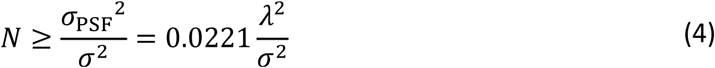

Based on the peak emission wavelengths of the fluorophores used in this manuscript, the theoretical minimum number of photons required to reach down to Ångström precision (<1 nm) would be:

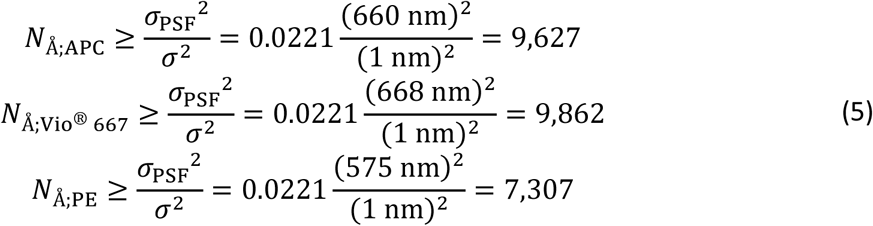

### Photobleaching of probes adsorbed to coverslip glass

In order to compare photon counts per localization under different excitation intensities (Fig. S2), the probes were applied to separate channels of a µ-Slide VI 0.5 (ibidi, #80607, 540 µm channel height) at low concentration for 1 min and then washed three times with PBS. For photobleaching of APC, a dilution of the α-tubulin-REA1136 antibody (DOL of 1) at a ratio of 1:50,000 was employed, while for Vio^®^ 667, 25 pM of an α-tubulin-Fab (DOL of 1.5; based on REAdye_lease™ reagent REAL979, Miltenyi Biotec) was applied. For each distinct ROI, the glass surface was brought into focus with the hardware autofocus and a time-lapse image series was acquired at a specific illumination intensity (e.g. 100% LED, 10% laser, etc.) until all probes were photobleached. Cross-comparison of the results from these distinct ROIs testing different illumination intensities was possible by restriction of the analysis to only the same number, *N*, of well-detected events in each ROI for a given fluorophore. Analysis with PROSPERO (see Materials and Methods) revealed much longer fluorophore lifetimes (Fig. S2B–C) and much higher photons collected per probe (Fig. S2D) at lower intensities. Higher photon collection per probe at lower intensities can be largely attributed to the highly nonlinear dependence of the photobleaching half-time on the applied intensity, with, e.g. a reduction of only 3× in illumination intensity (from 16 W cm^−2^ at 5% laser to 5.3 W cm^−2^ at 100% LED) generating a 150× longer photobleaching half-time for the brightest APC events (from 0.2 s to 30 s, Fig. S2C). For a linear relationship between applied illumination intensity and photobleaching rate (equal to the inverse of the half-time), an unchanging photon collection distribution for Fig. S2D would have been predicted, as photon collection should scale with the product of the illumination intensity and the half-time, which would be constant. Our results are highly discrepant with a linear relationship, proving that the probes are in the nonlinear regime over the assayed intensity range. To address localization precision, we redisplayed the photon distributions from Fig. S2D on the *y*-axis in the top panel of Fig. S2E, with corresponding distributions for the localization precision determined with PROSPERO, using drift correction based solely on the ensemble of tracked probes as no brightfield images were available (Materials and Methods). The circles give the median value for the distributions, with the “X” symbols in the lower panel giving the minimum theoretically achievable precision (see previous section) expected for the respective median photon count event from the upper panel. The median observed precisions in the lower panel were up to three times worse than those expected from the theoretical limit based solely on photon number, due largely to a significant flat background component required for the 2D Gaussian fits with PROSPERO. The level of background in these experiments was consistent with that created by autofluorescence from the coverslip glass and optical elements in our microscope setup (see the “Control coverslip measurements” section below).

### Actin labeling with phalloidin-Atto643

For our resolution of phalloidin-Atto643 with FILM, we used CB-fixed HeLa cells as described above. In the analysis of our other experiments, a significant flat background component was necessary for the fitting of isolated probe events, which limited the achievable precision (shifted the distribution for precision away from the theoretical limit, as in Fig. 2G). We conjectured that this background came from out-of-focus elements in our system. Restricting the total light put into the system, by limiting the illuminated FOV, might reduce this homogeneous background. We therefore restricted the area of illumination in the focal plane to 20% (excitation pinhole set to 20%, illuminating ∼27% of the camera FOV), which allowed for a four-fold reduction in background and consequent two-fold increase in localization precision (Fig. S28). Instead of 15% detected with precision <2 nm and 2% in the Ångström regime (Figs. S28B–C), now fully 50% could be detected with precision <2 nm and 20% in the Ångström regime (Figs. S28E–F). The average spacing between labeled actin subunits in the isolated filaments shown in Figs. 5C–F and S30 was ∼50 nm, which, accounting for the 2.75 nm distance between adjacent actin subunits and their associated phalloidin binders (*39*, *40*), implies a labeling efficiency of roughly 5%. As 1475520 separate probes were detected over the six relatively large HeLa cells (six separate nuclei were observed in the brightfield image, not shown), or ∼250,000 localizations per cell, this implies an amount of F-actin per HeLa cell of ∼5×10^6^, which should be considered a *minimum* value due to the selection effect arising from easier identification of the analyzed single filaments (Figs. 5C–F) that were likely labeled higher than average. For the two highly precise localizations in Fig. 5I separated by 4.34 ± 0.55 nm, the error bar of *σ* = 0.55 nm = 5.5 Å was determined by standard propagation of error, i.e. 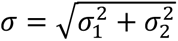, based on the separate localization precisions of 4.5 Å and 3.1 Å. The standard Gaussian-based definition of resolution of FWHM = 2.35 *σ*_0_ applies for objects localized with *identical* precisions, *σ*_0_, with the above propagated error calculation for the uncertainty of their displacement becoming 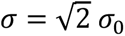, or 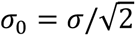. Plugging this in for the FWHM resolution formula, gives a generalized expression for the resolution that we obtained in terms of FWHM of 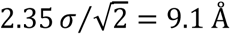.

### Autofluorescence background due to composite glass in the microscope setup

To better characterize the background for FILM, we performed imaging of just the coverslip glass (Fig. S32A). We assessed the background of the standard 170 µm thick borosilicate coverslips used for all FILM experiments described here. As a control, we also measured the background of a 150–180 µm thick quartz (fused silica) coverslip. Coverslip glasses were soaked in isopropanol for several minutes, removed, and then allowed to dry. The borosilicate coverslip was mounted on the microscope for observation with either the 63× oil immersion objective or the 63× water immersion objective on our setup and with, respectively, oil or distilled water immersion (see Materials and Methods for further details of the microscope setup). 5 s exposures at 100% LED for the 631/33 nm excitation channel (Table S2) were acquired with excitation pinholes of either 100% or 20%, as indicated in Fig. S32. The quartz coverslip was observed only with the 63× water immersion objective. The background levels for the borosilicate coverslip were nearly identical to that observed in the phalloidin datasets (see Fig. S28) for excitation pinholes corresponding to 100% (Fig. S32B) and 20% (Fig. S32C–D). For the 20% pinhole, we observe a flat background and a plateau, with the latter due to autofluorescence from the near-focus portion of the coverslip. To rule out any possible autofluorescence signal from the immersion oil, we switched to a water immersion objective. Here, the background was slightly reduced (Fig. S32E–G) most likely solely due to the lower N.A. of the objective (no need to assume significant autofluorescence from the oil). We then switched from the standard borosilicate coverslip glass to a quartz coverslip glass (Fig. S32H–J), which should have significantly less autofluorescence (*56*, *57*). Use of the quartz coverslip glass reduced the background by half. For the 20% FOV, as well, no circular profile of the restricted FOV (illumination aperture) could be observed, indicating that the quartz coverslip contributes no significant autofluorescence. Nevertheless, roughly half of the background remained, most likely due to the roughly ten or so lenses (also made of glass composites similar to borosilicate) of the objective.

### Speed of acquisition with FILM

To better appreciate the speed of FILM for acquisition of events at molecular resolution, we compare it with MINFLUX (*58*). For the full α-tubulin acquisition in HeLa cells with FILM (Fig. S3), 15% of the total 14,176,305 probe localizations acquired over 9,564 frames (5 s each) were localized with σ<2 nm (45 localizations/s). For the reduced FOV acquisition of phalloidin-Atto643 binding to F-actin, 50% of the total 1,475,520 events acquired over 5,712 frames (5 s each) were localized with σ<2 nm (26 localizations/s). For a comparable MINFLUX run (*58*), 8 localizations/s at σ<2 nm was claimed for super-resolution of the nuclear pore channel component, Nup96 (19,400 localizations over 40 min for the dataset shown in their Fig. 3d). For their experiment, nuclear pore channels containing Nup96 were expressed in a homogeneous manner over a 10×15 nm FOV that sampled a portion of the nuclear membrane within a large U2OS cell, providing a comparison tilted slightly in favor of MINFLUX due to target homogeneity over their *entire* FOV. Based on the two FILM runs described above, FILM was therefore at least 5.6-fold and 3.2-fold faster than MINFLUX at generating σ<2 nm localizations.

### Ethics statements

For all studies using human primary tonsil or colon cancer tissue, written informed consent was obtained following the guidelines of the approved Universitätsmedizin Göttingen, and Cologne Review Board protocol, respectively. All procedures were carried out in accordance with the European Communities Council Directive European Communities Council (86/609/EEC) and (2010/63/EU).

## Acknowledgments

We thank Anthony de Vries, Werner Müller, Robin Diekmann, Michael Wagner and Michael Knop for their invaluable technical assistance and helpful discussions.

## Author contributions

Conceptualization: AK

Formal analysis: JB, AK

Funding acquisition: MS, AK

Investigation: JB, AL, TF, AK

Methodology: JB, AK

Project administration: MS, AK

Resources: AL, TF

Software: AK Supervision: MS, AK

Visualization: JB, AK

Writing – original draft: JB, AK

Writing – review & editing: JB, MS, AK

---

**Fig. S1.**
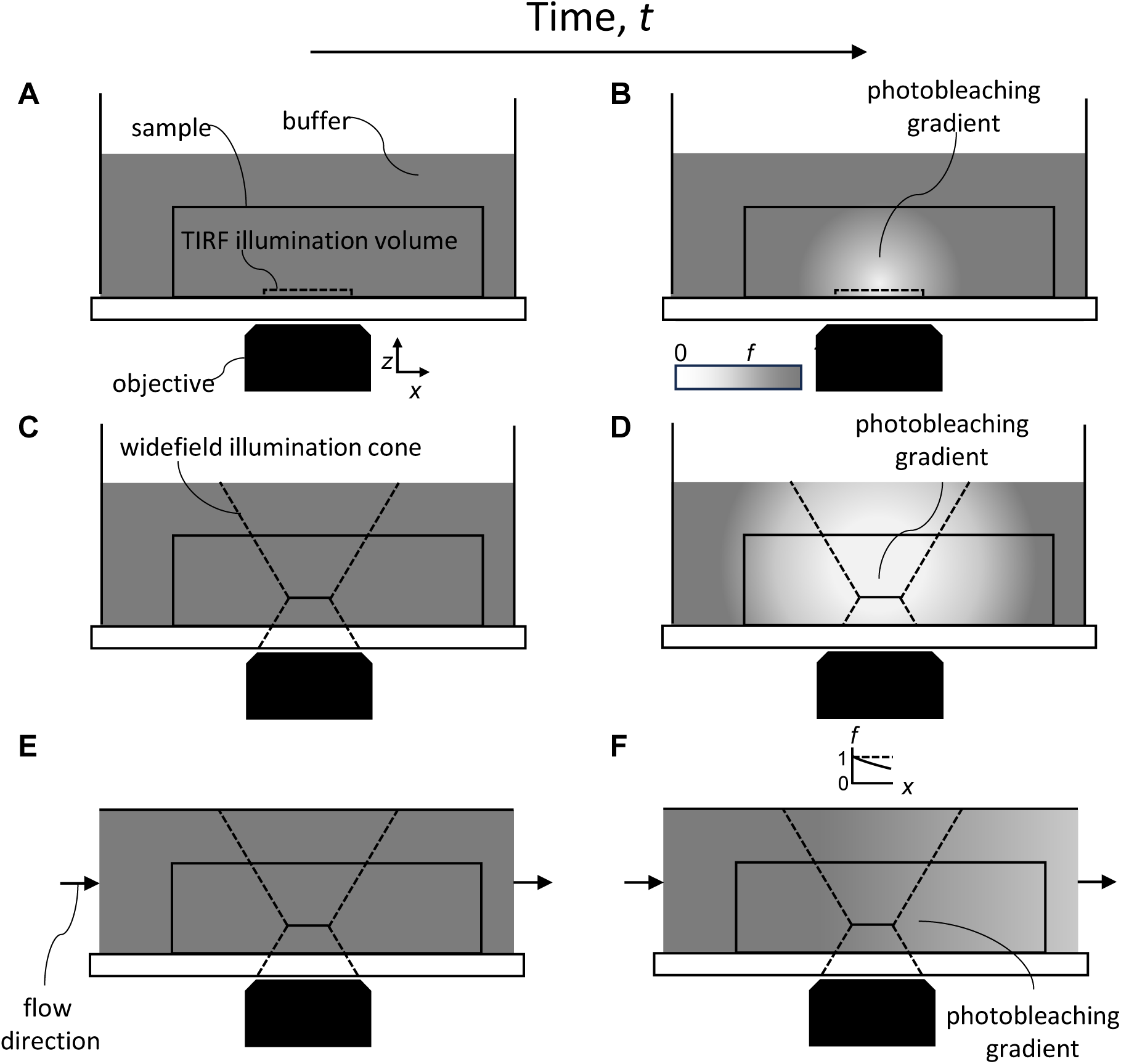
Photobleaching of free probes for static incubation vs. continuous flow. (A) Initial condition of no photobleached fraction of free fluorescent probes for SMLM based using static fluid conditions and TIRF microscopy. (B) Later timepoint for the setup in A showing the presence of a bleached population (lighter shading) of free probes both within and surrounding the illuminated region. (C) Initial condition for the fraction of free fluorescent probe for SMLM using static fluid conditions and widefield microscopy. (D) Later timepoint for the setup in C showing an even larger bleached region than displayed for TIRF in B. (E) Initial condition for the fraction of free fluorescent probe using FILM based on continuous transport of fluorescent probes over the sample (now mounted inside a microfluidic sample chamber) and for acquisition with widefield microscopy. (F) Later timepoint for the setup shown in E. Due to the continuous flow-based delivery of fresh probe from left to right over the sample, a shallow, steady-state gradient of bleached free probe quickly forms and persists throughout the entire acquisition described by *f*(*x*) = exp[−*x* ln(2)/(*VT*_1/2_)], with *x* the distance along the flow within the FOV, *V* the flow speed, and *T*_1/2_ the photobleaching half-time (see also Fig. 1C). For standard dyes with conventional photobleaching kinetics and fast enough flow speed, this gradient can be limited to less than a few percent over the full length, *L*, of the FOV.

**Fig. S2.**
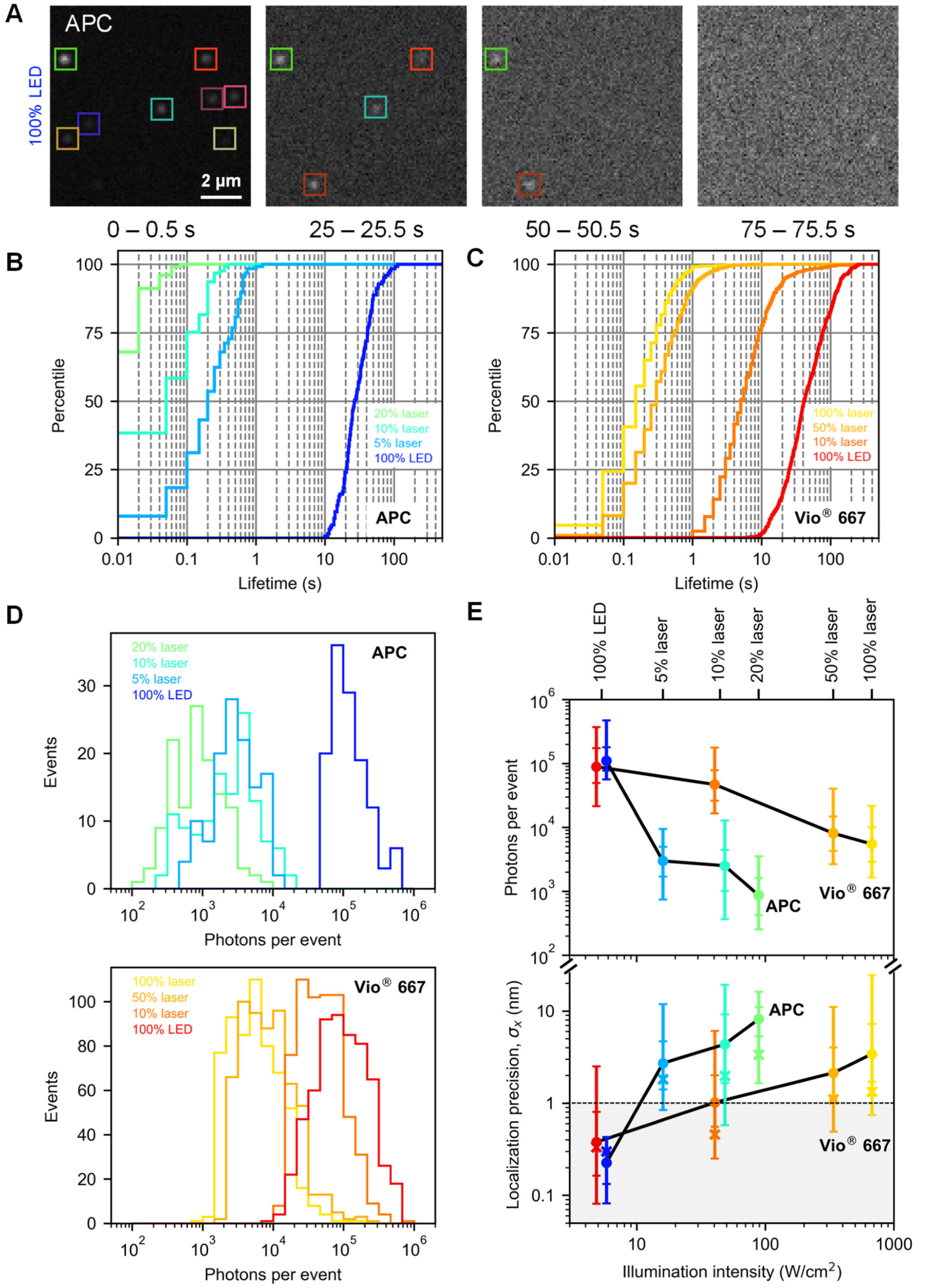
Photon collection efficiency and localization precision for probes adsorbed on glass. (A) A microfluidic chamber was incubated for 1 min with a buffer containing a probe conjugated to APC, allowing the probe to attach non-specifically to the glass. The chamber was washed multiple times. Widely spaced ROIs were then imaged with different intensities of illumination provided by an LED or a laser. Shown are multiple timepoints (0.5 s frames) of a zoomed-in portion of a region that was illuminated with 100% LED. Events were analyzed with PROSPERO to determine the total number of detected photons for each probe (tracked across its relevant frames) as well as its position and localization precision. Same color-coding of the bounding boxes across the different frames indicates the successful tracking of each detected probe. A fixed subset, *N*, of well-detected probes in each ROI (*N* = 125 for APC, *N* = 600 for Vio^®^ 667) was analyzed at each illumination intensity to allow direct comparison of photon collection efficiency vs. illumination power. (B, C) Cumulative lifetime distributions of the *N* well-detected probes for APC (degree of labeling, DOL = 1) and Vio^®^ 667 (DOL = 1‒2), respectively, at the different illumination intensities. (D) Histograms of total photons collected per probe for the *N* well-detected probes at the different illumination intensities. For both probes, a strong inverse dependence of total photons per probe on illumination intensity was observed. (E) Statistical distributions of photons per event (top panel) and 1-σ localization precision (bottom panel). Horizontal ticks for each distribution indicate the 5%, 25%, 75%, and 95% cumulative values, with the filled circle indicating the median value at 50%. The “X” symbols in the lower panel indicates the best possible theoretical precision one would obtain based purely on total photons collected (ignoring other non-Poissonian noise sources and background) for the event with median photon number from the upper panel. For all datasets, the actual median localization precision (filled circle) lies within a factor of 1‒3× of this theoretical limit.

**Fig. S3.**
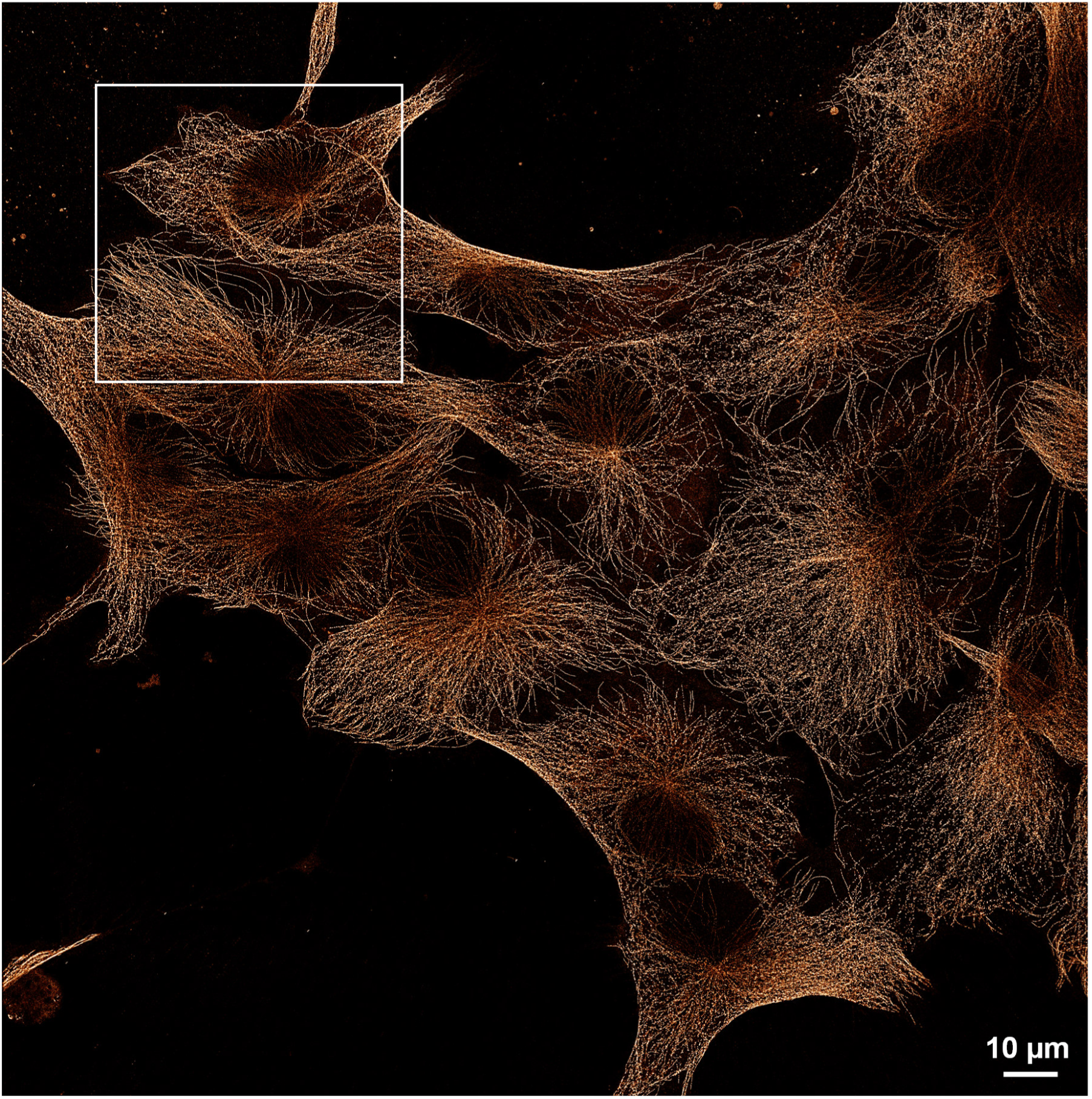
Full camera field of view for FILM-resolved α-tubulin in HeLa cells. Full FOV (211.3 x 211.3 µm) for the α-tubulin staining of PFA-fixed and permeabilized HeLa cells presented in Fig. 2B (boxed region here). Image displayed using the “Red Hot” Lookup Table (LUT) in ImageJ (*49*).

**Fig. S4.**
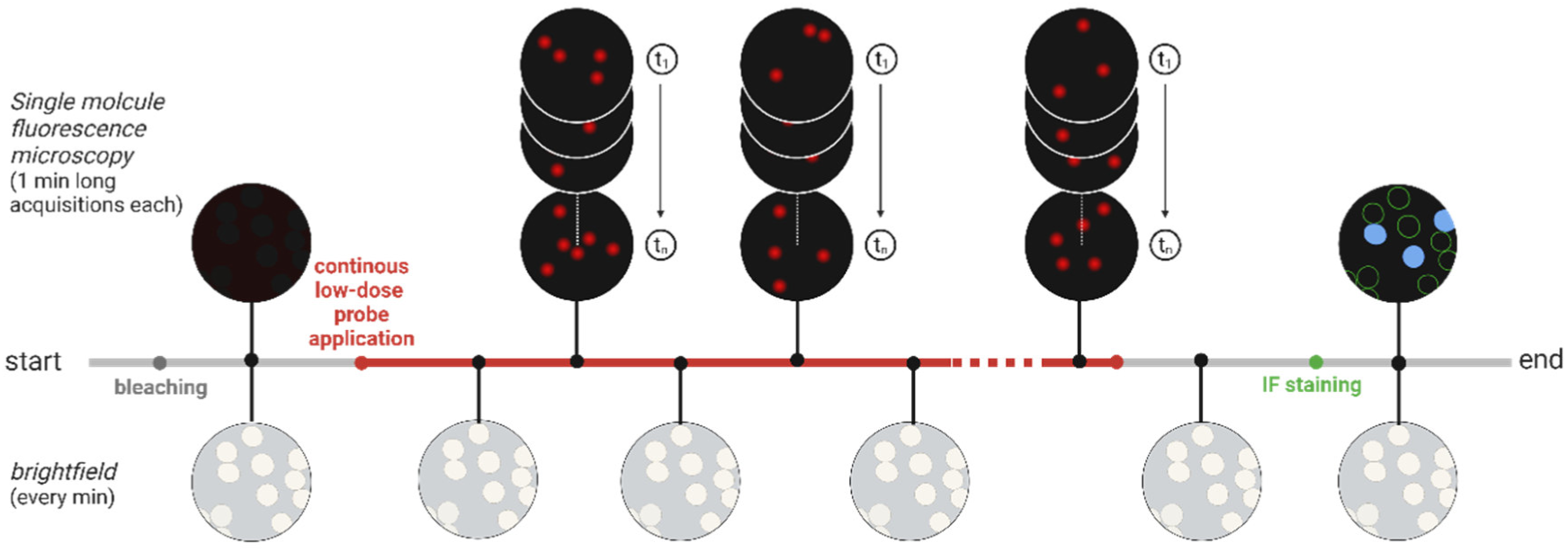
Experimental design for FILM. An initial bleaching of the sample FOV at 100% LED was carried out and confirmed by imaging. A low concentration of the probe was then flowed over the sample in a continuous fashion (red line) and the sample was continuously imaged with single brightfield images alternated with short fluorescence movies of 1 min duration (with frames of typically 5 s). For the analysis, registration information using the brightfield images and, separately, probe tracking across the fluorescence movies was combined for high accuracy registration at both short and long timescales with our analysis code, PROSPERO (see Materials and Methods). For most datasets, a conventional IF image (e.g. to compare specificity) was also obtained at the end of the FILM acquisition by a final incubation with the probe at high concentration (“IF staining”) followed by washing and imaging. Created in BioRender. *Bartmuß, J. (2024)* https://BioRender.com/c70r146

**Fig. S5.**
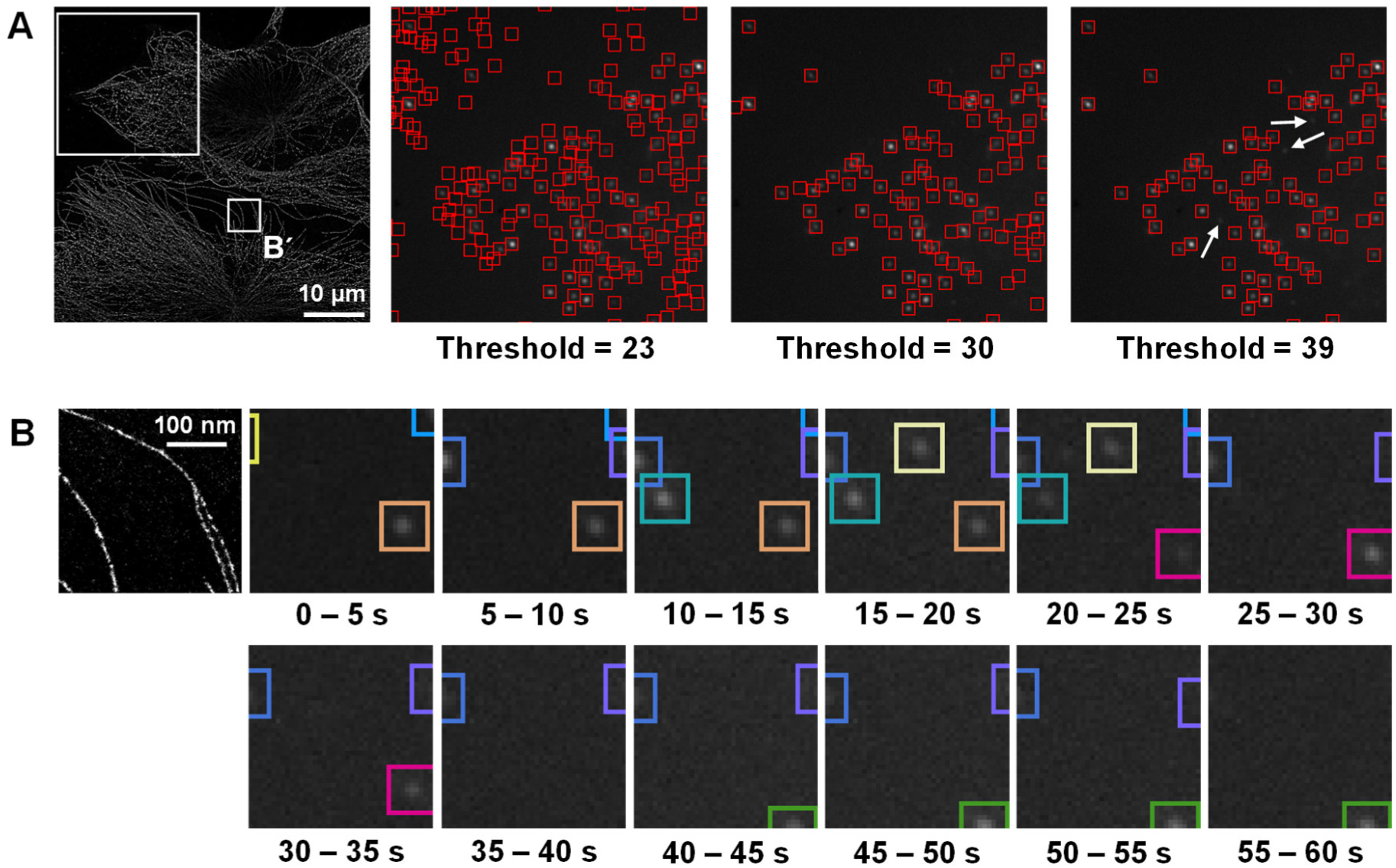
Probe detection and tracking with PROSPERO. (A) FILM image of tubulin in HeLa cells from Fig. 2B is redisplayed. For the large ROI in the upper left, different thresholds for event detection are shown in the frames to the right (events are indicated by red squares consisting of 9×9 pixels), allowing determination of the optimal threshold. A single threshold must be chosen by the user, which is then applied to all frames of the dataset, generating a list of events for each frame. (B) Zoom-in of the small boxed region in A (identical to Fig. 2C). In the panels to the right, the individual frames over a 1-min fluorescence movie are shown. In each frame, a 2D Gaussian plus flat background was fit to each thresholded event to localize it. This localization information was then used to track events across frames, with constant color coding from frame-to-frame demonstrating successful tracking.

**Fig. S6.**
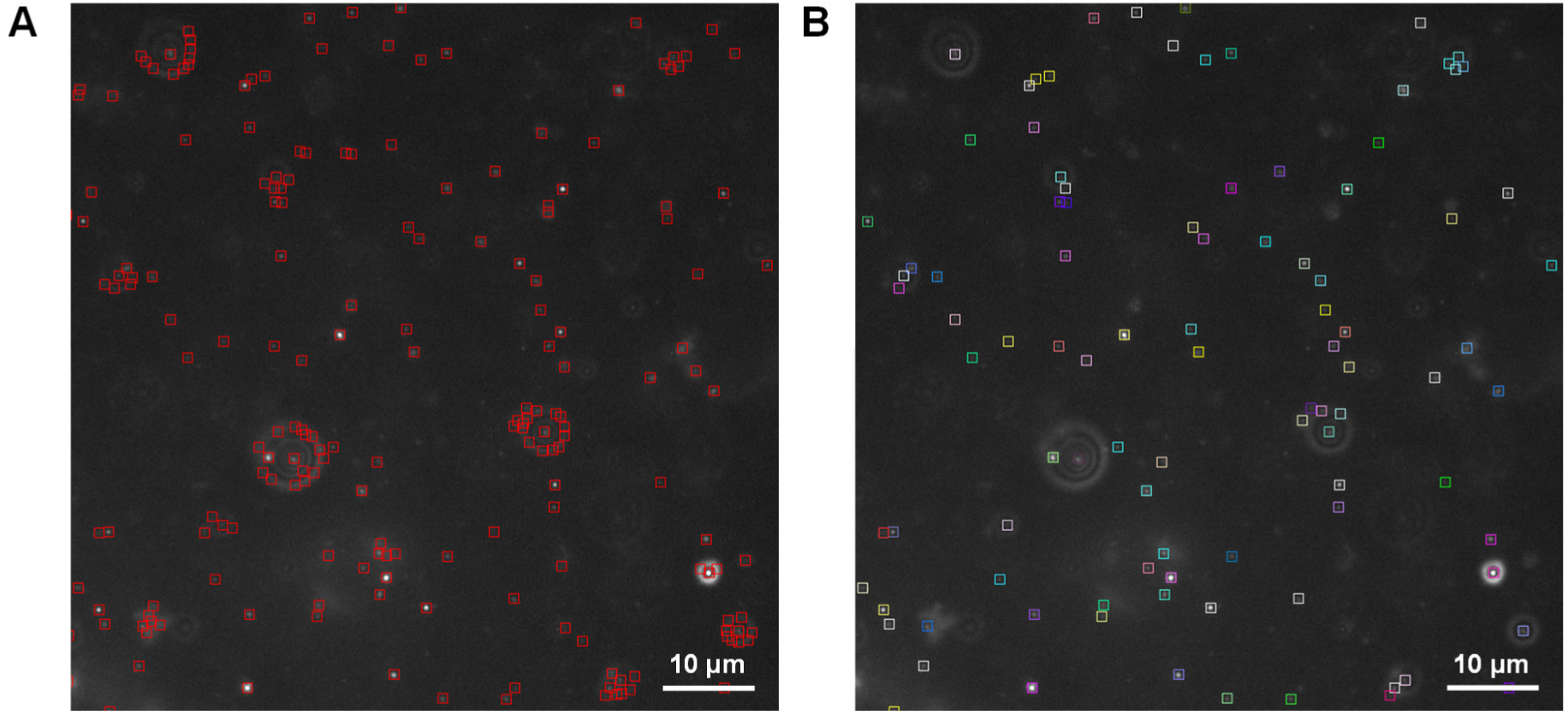
Filtering of thresholded events by radial symmetry with PROSPERO. (A) Due to the extremely high collection efficiency of FILM, out-of-focus events are also robustly detected, generating images of a central peak surrounded by multiple concentric diffractive rings. Simple threshold detection over the small red detection windows (9×9 pixels) will therefore pick up the same particle multiple times, as observed for several out-of-focus events in the displayed 5 s frame taken from the CD81 staining of healthy tonsil tissue (Fig. 4). For most datasets, it was therefore necessary to filter the thresholded events for radial symmetry by determining the center of radial symmetry for each event within the detection window (see Materials and Methods) and comparing it with the center of its 2D Gaussian fit. (B) Removal of events with more than 1-pixel (103.17 nm) deviation of their radial and Gaussian centers, followed by probe tracking over the entire acquisition, resulted in the displayed, color-coded, tracked events for the frame shown in A. Note the absence here of most of the spurious detections on the ring-like structures in A.

**Fig. S7.**
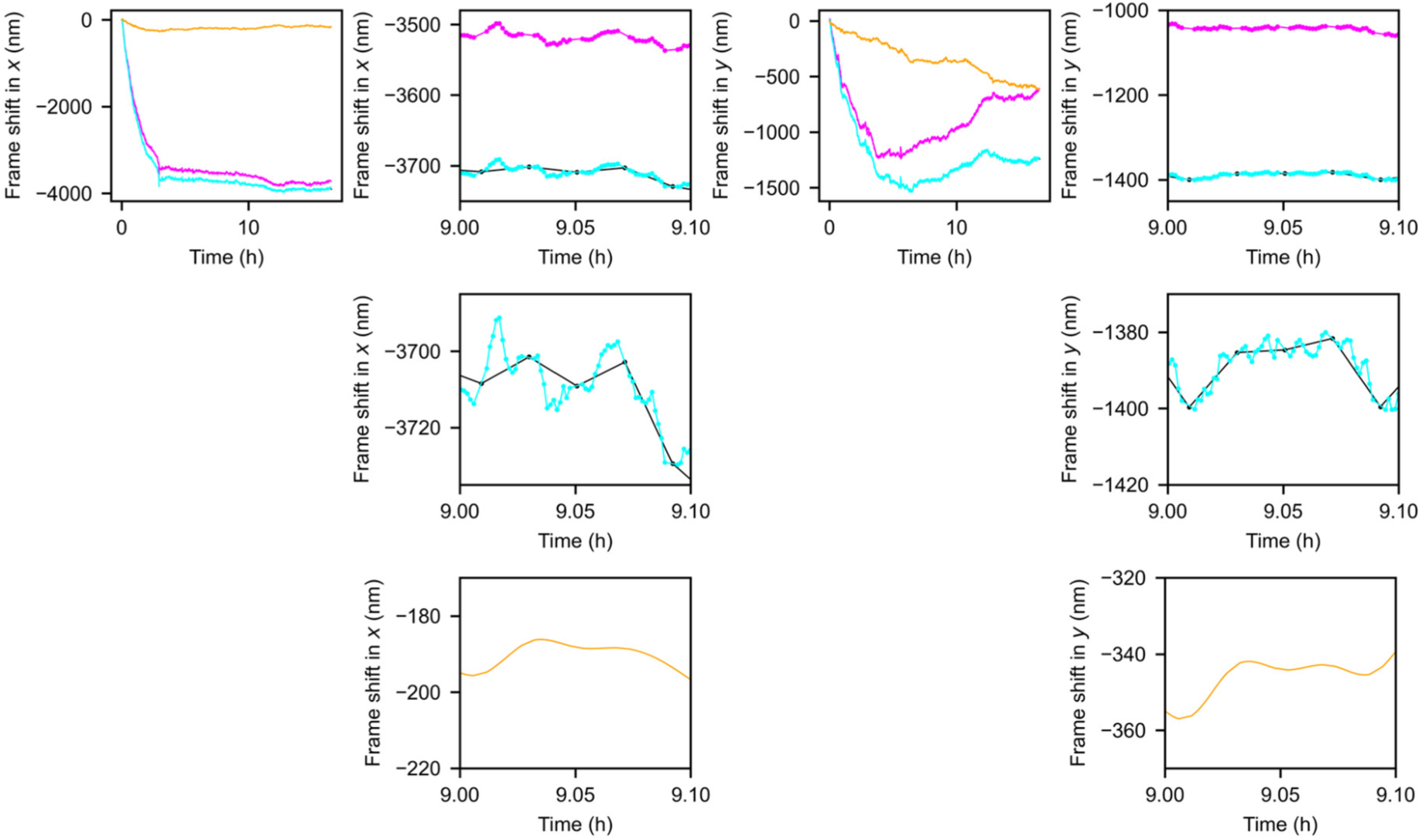
Sub-nm drift correction with PROSPERO by combining brightfield image registration with probe tracking information. In the upper-left panel, estimates of the shift of the sample in *x* over the entire acquisition based solely on the brightfield images (black, obscured by the overlaid cyan curve) or the tracked-probe information (magenta) are shown, with the former estimate accurate on long timescales and the latter accurate on short timescales. To bring the magenta line into concordance with the black line, the displayed spline-fitted function (orange) is added to generate the final estimate of the frame shift (cyan) now accurate on both short and long timescales. Immediately to the right is a zoom-in of the abscissa from 9.00 to 9.10 hours. Immediately below is a further contrasting of the ordinate to show the high discrepancy of the black line (linearly interpolated over the 1-min gap in brightfield images) from the cyan curve, reflecting the additional information contained in the probe-tracking magenta curve. In the bottom panel, the spline-fitted function (orange) is redisplayed. Note that the cyan curve wanders in relatively smooth trajectories, implying that the drift estimates based on averaged tracked probe shifts between successive frames are correlated on short timescales and not merely “noise”, which would have instead appeared as sharper ± deviations from one time point to the next. The deviations of the cyan curve with respect to the black linearly interpolated curve are out to ±15 nm, specifying the level of inaccuracy of registration based solely on the intermittent brightfield images. The panels on the right provide the same information for drift estimation along the *y* axis. Due to the longer frame acquisition time of FILM (5 s per frame) only sample drift must be corrected for and not higher frequency vibrations, which for conventional SMLM at higher frame speeds (e.g. 50 ms per frame) pose severe challenges for accurate registration (*59*).

**Fig. S8.**
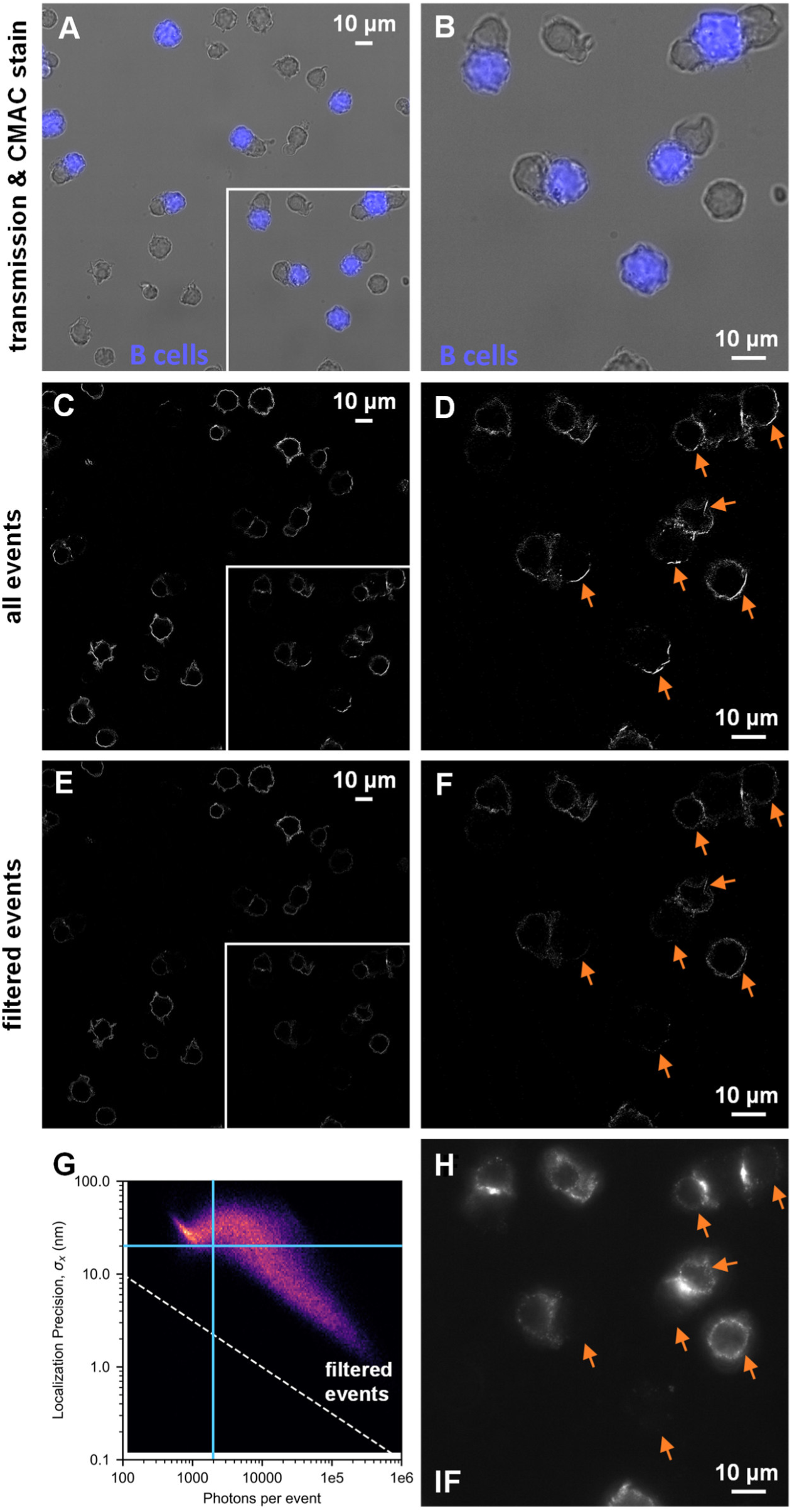
Filtering out of false localizations arising from sharp optical features backlit by the free probe background. (A) Overlay of image of CMAC pre-staining of B cells with a brightfield image. (B) Zoom-in of boxed region in A. (C) All detected events for FILM acquisition of anti-CD28-APC on T and B cells. (D) Zoom-in of boxed region in C. Orange arrows indicate regions containing many false positive detections. (E) Filtered events. (F) Zoom-in of boxed region in E. Orange arrows (same as in D) show the successful removal of excess intensity due to false positives by the filtering. (G) Filter settings. Only events with >2000 photons and < 20 nm localization precision were selected for display in panels E and F. (H) Conventional IF staining (103.17 nm pixels) generated at the end of the FILM acquisition by incubating with the antibody at the standard concentration and time recommended by the manufacturer. Orange arrows show both the absence of excess intensity for the regions in D (containing false positive events). Good overall correlation is found with the image of filtered events in F. Panels C–F and H use 10 nm pixels and 3-pixel Gaussian blurring.

**Fig. S9.**
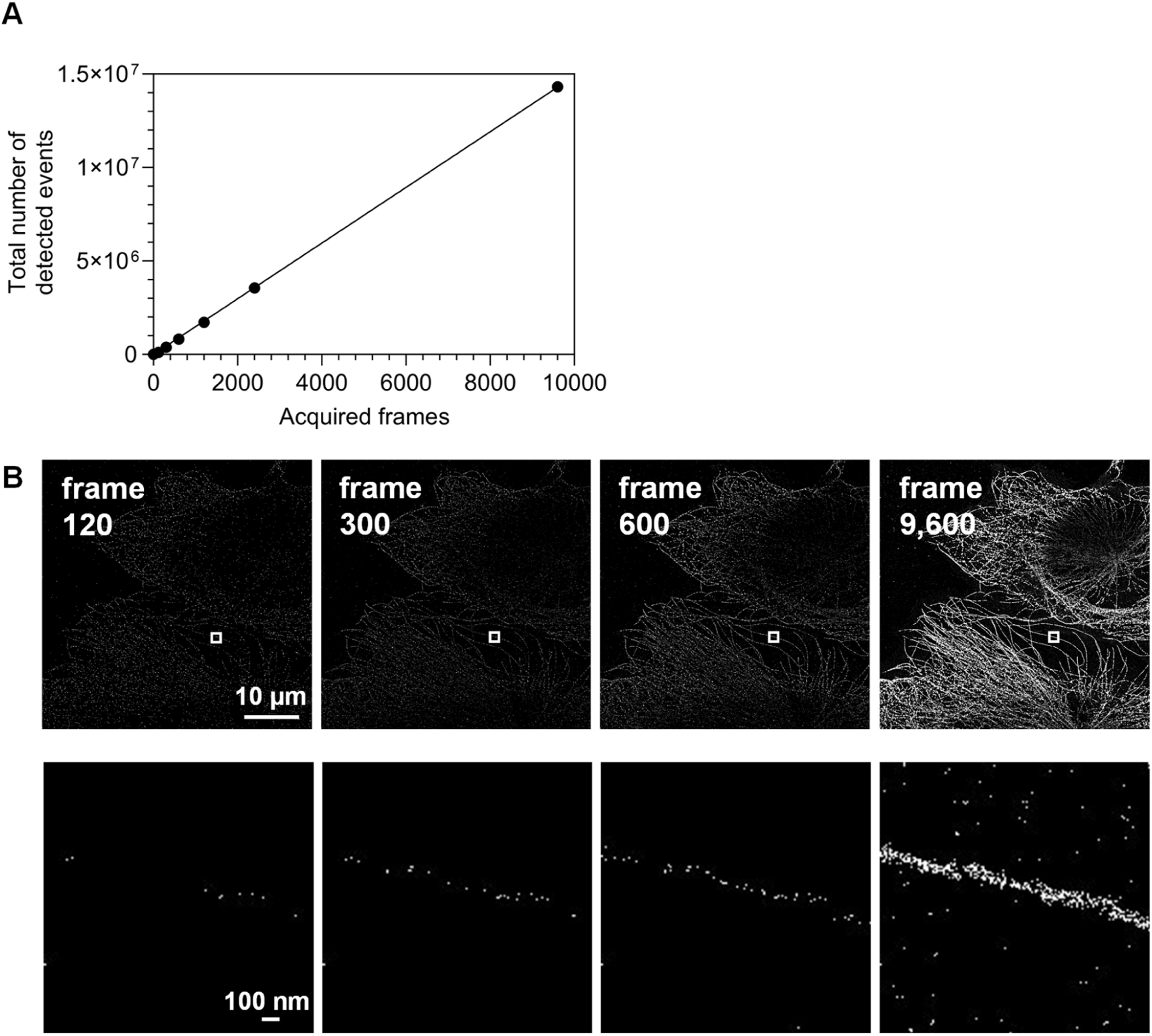
Temporal acquisition of events for acquisition of α-tubulin in HeLa cells with FILM. (A) A steady, unchanging acquisition rate for the localizations was observed for the FILM staining of α-tubulin in PFA-fixed and permeabilized HeLa cells (full FOV shown in Fig. S3) over the entire duration of the experiment. (B) Top row: Zoom-in corresponding to the boxed region of Fig. S3 for the specific indicated 5 s frames (103.17 nm pixels). Bottom row: Zoom-in of the boxed region in the top row to show the individual localizations for an isolated microtubule (10 nm pixels with 2-pixel Gaussian blurring).

**Fig. S10.**
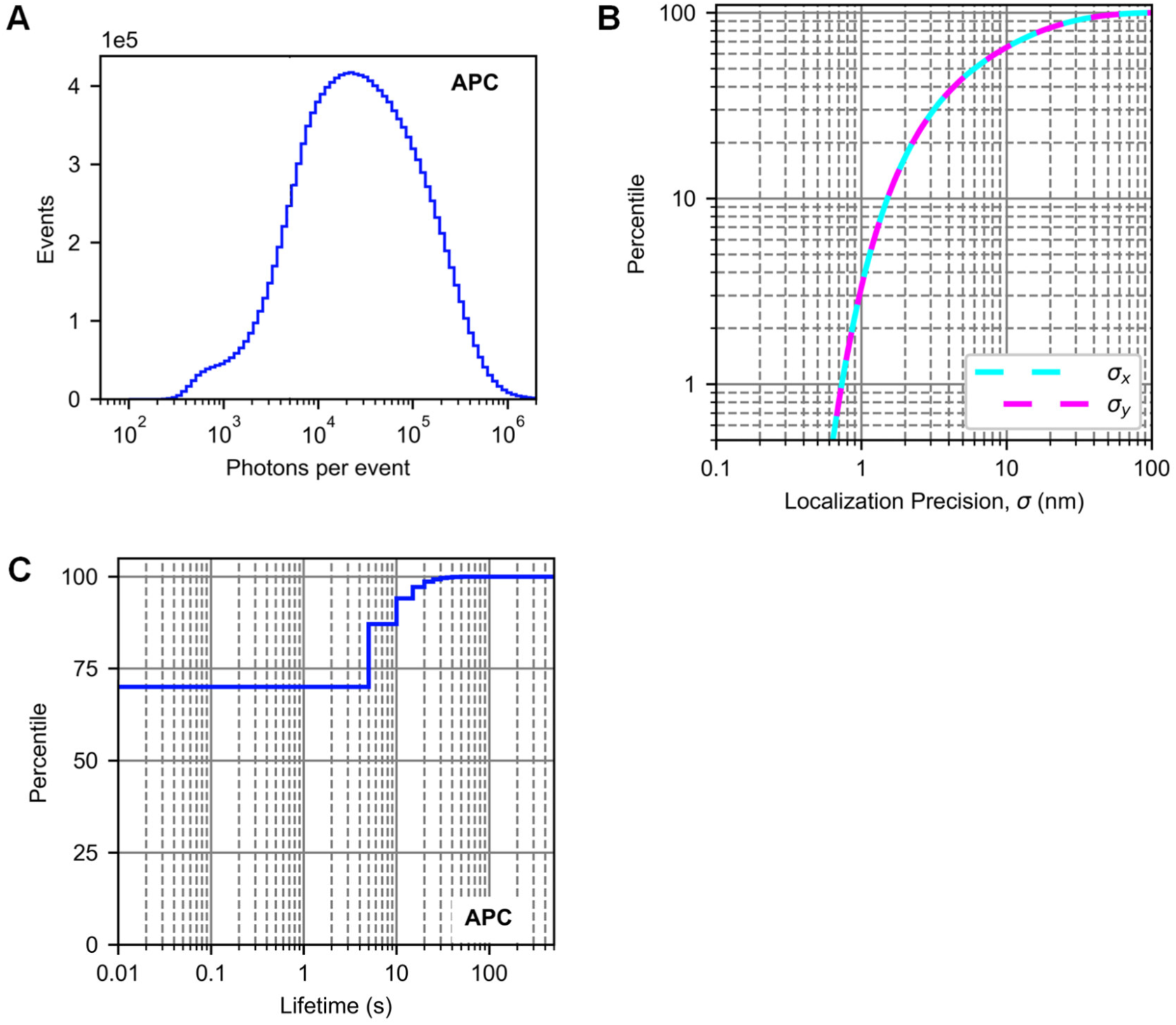
Photons per event, localization precision, and probe lifetimes for the FILM acquisition of α-tubulin in HeLa cells. (A) Histogram giving the distribution of events with different numbers of photons for the 14,176,305 total events observed for the anti-α-tubulin-APC antibody probes over the full camera FOV (Fig. S3). (B) Cumulative distribution of the 1-*σ* localization precision of the probes along the *x* and *y* directions. (C) Cumulative distribution of probe lifetimes. The distribution truncates at the frame exposure of 5 s at a value of ∼70%. Based on the curvature of the distribution, the apparent *in situ* photobleaching half-time of APC for FILM can be inferred to be on the timescale of a few seconds.

**Fig. S11.**
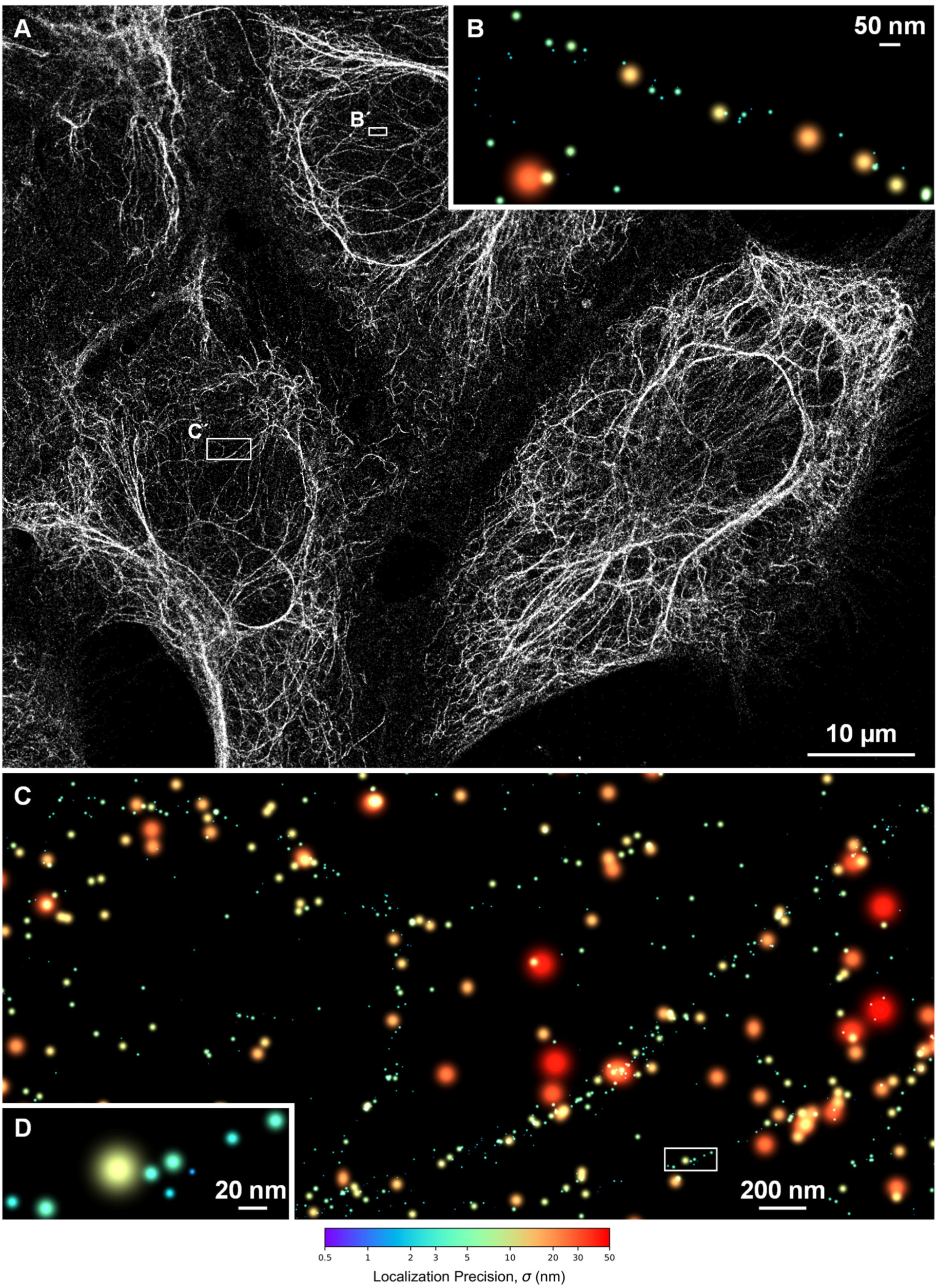
FILM image of cytokeratin in HeLa cells. (A) Resolution of cytokeratin in PFA-fixed and permeabilized HeLa cells using a pan-cytokeratin antibody conjugated to APC with FILM (10 nm pixels with 3-pixel Gaussian blurring). (B) Zoom-in of the boxed region B’ from A. (C) Zoom-in of the boxed region Ć from A. (D) Zoom-in of the boxed region in C. For panels B–D, localizations were represented as peak-normalized Gaussians with color-coded precision according to the colorbar.

**Fig. S12.**
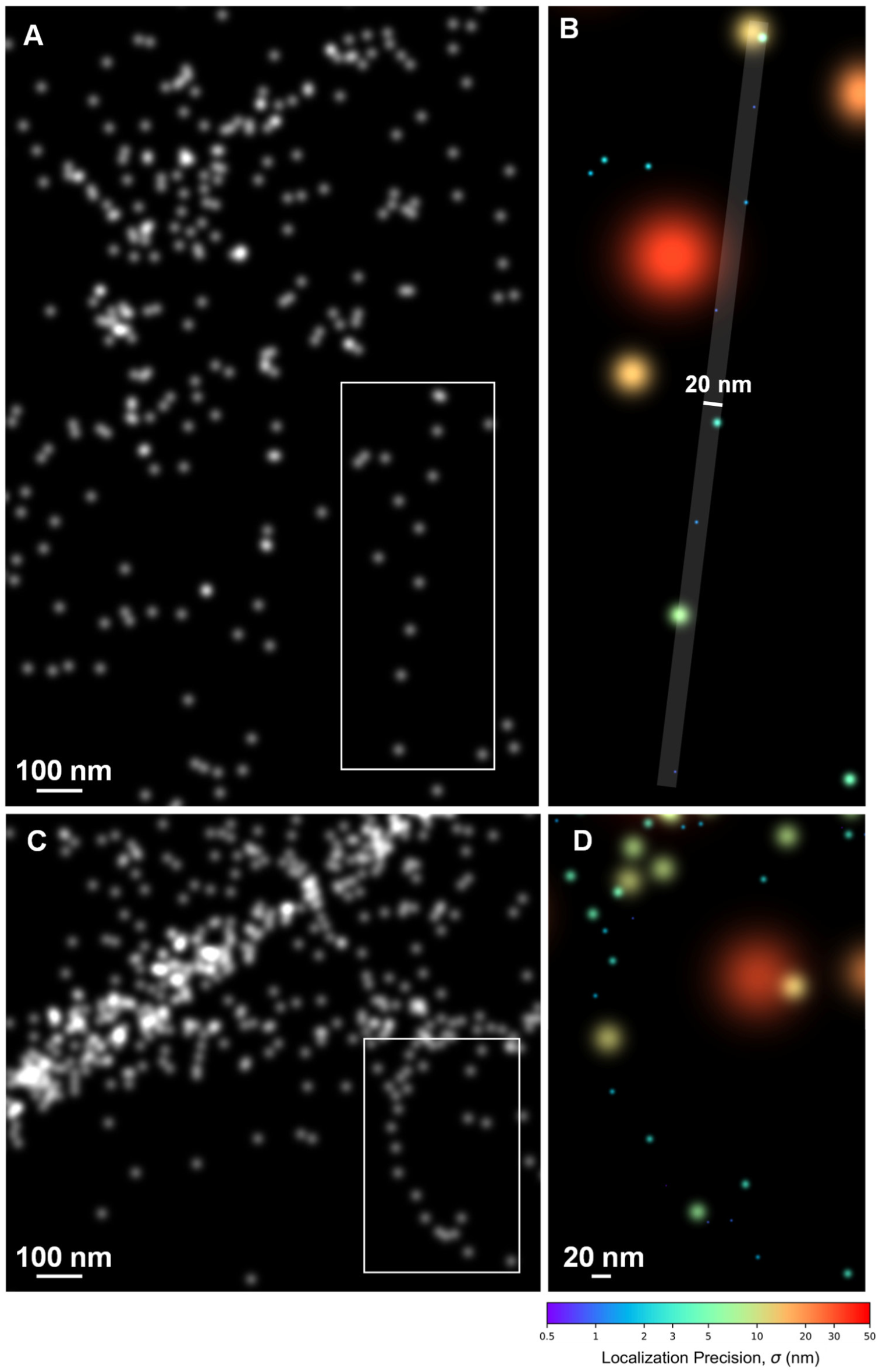
Isolated cytokeratin filaments in HeLa cells. (A) ROI from the full cytokeratin dataset (Fig. S11) containing a straight, sparsely labeled filament (5 nm pixels with 2-pixel Gaussian blurring). (B) Zoom-in of the boxed region in A. All localized positions (centers of Gaussians) were contained within the overlaid strip of width 20 nm. (C) ROI from the full cytokeratin dataset (Fig. S11) containing a curving, sparsely labeled filament (5 nm pixels with 2-pixel Gaussian blurring). (D) Zoom-in of the boxed region in C. For panels B and D, localizations were represented as peak-normalized Gaussians with color-coded precision according to the colorbar.

**Fig. S13.**
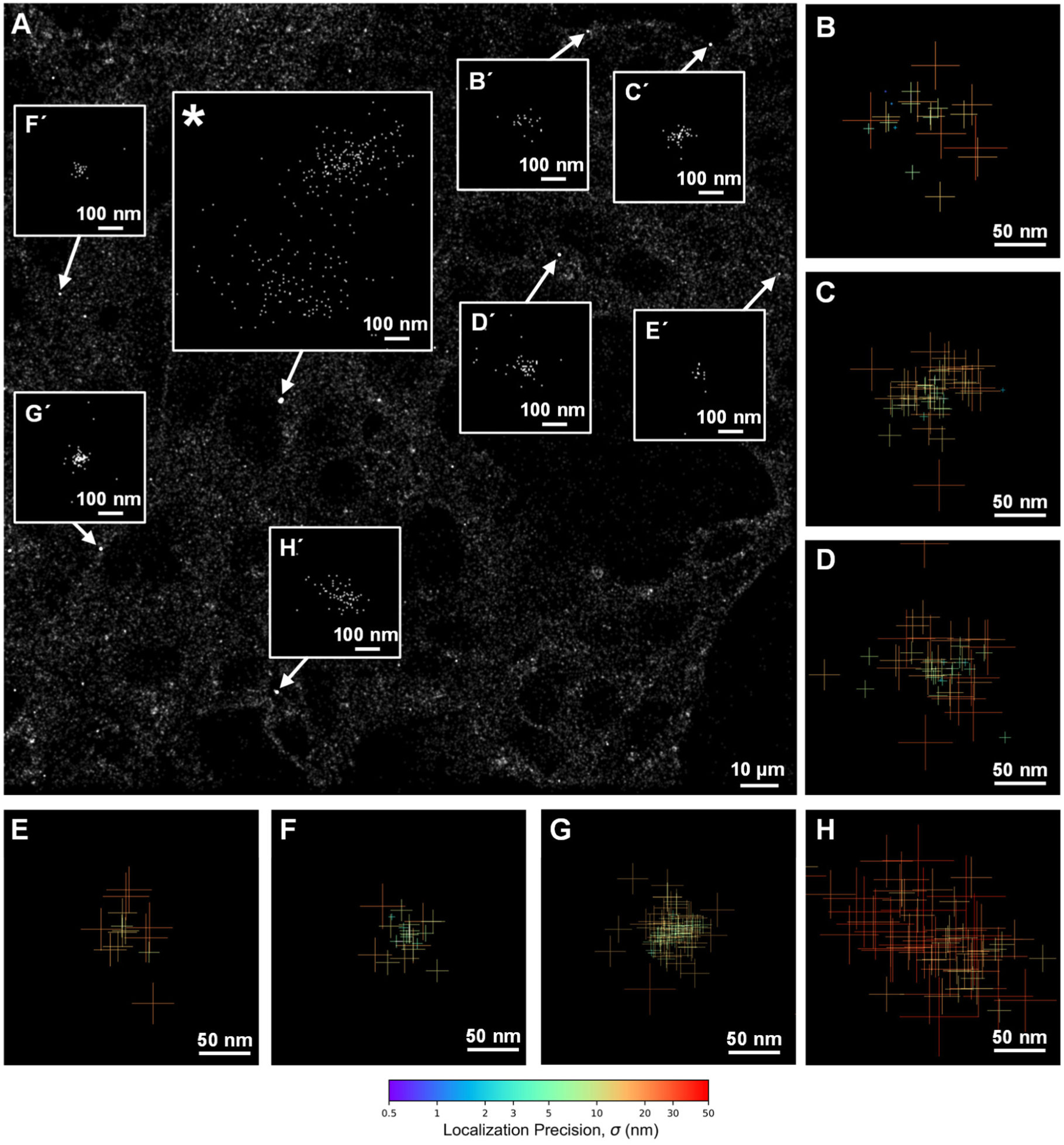
FILM acquisition of LAMP1 vesicles in HeLa cells. (A) FILM acquisition of an anti-LAMP1-APC antibody in PFA-fixed and permeabilized HeLa cells (103.17 nm pixels with 2-pixel Gaussian blurring). Boxed regions show zoomed-in views (5 nm pixels) of the brightest concentrations from the image, which were consistent with single, round LAMP1-rich vesicles of sizes 50–200 nm (aside from the *-labeled boxed region, which appears to contain two significantly larger vesicles). (B–H) Zoom-ins of respective boxed regions in A. Localizations are represented as crosses (indicating the ±1-σ localization along *x* and *y*) with color-coded precision according to the colorbar.

**Fig. S14.**
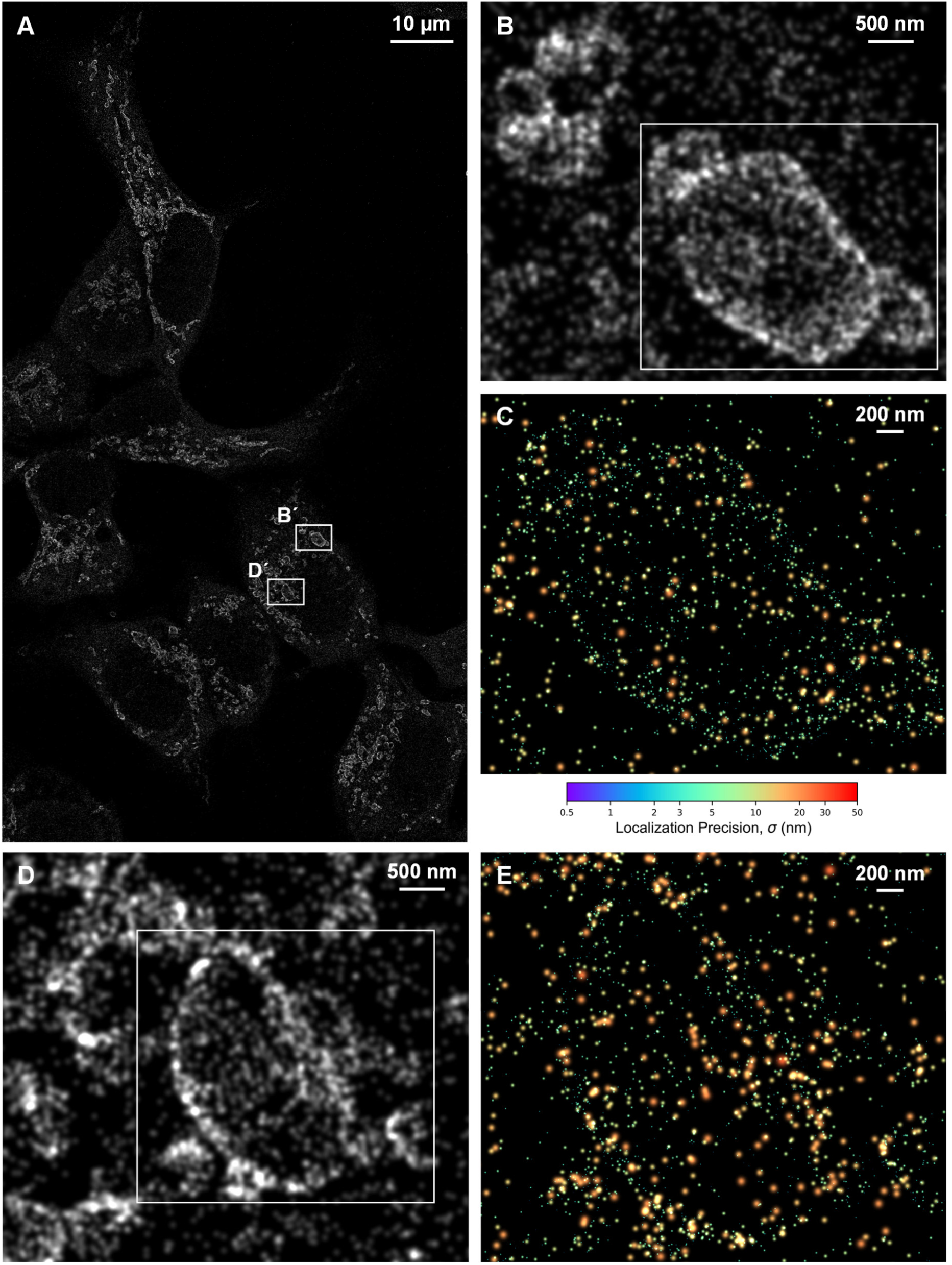
Resolution of mitochondrial marker TOM22 in HeLa cells with FILM. (A) FILM acquisition of anti-TOM22-APC in PFA-fixed and permeabilized HeLa cells. (B) Zoom-in of respective boxed region in A. (C) Zoom-in of boxed region in B. (D) Zoom-in of respective boxed region in A. (E) Zoom-in of boxed region in D. Panels A, B, and D use 10 nm pixels with 3-pixel Gaussian blurring. In panels C and E, localizations are represented as peak-normalized Gaussians with color-coded precision according to the colorbar.

**Fig. S15.**
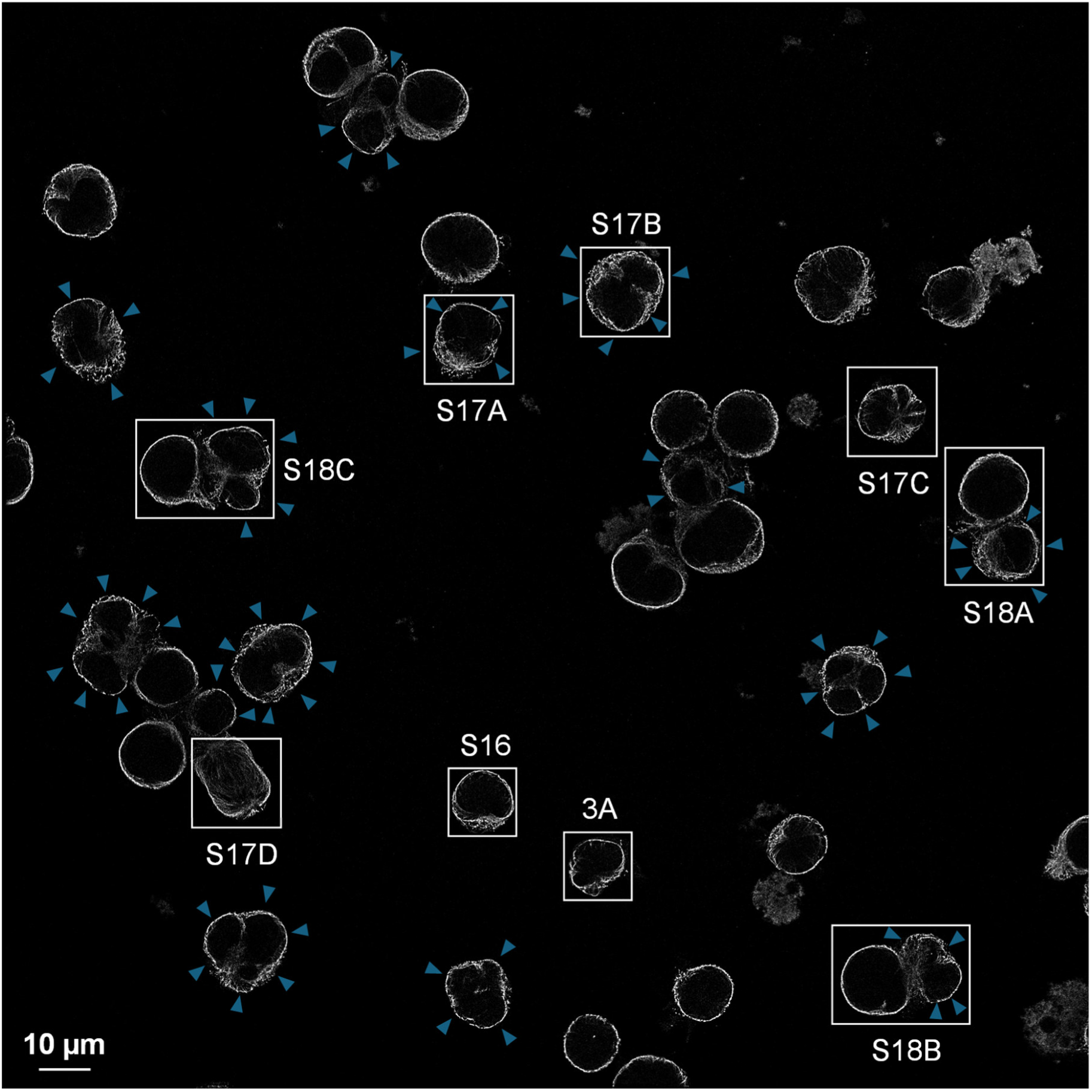
Full FOV for FILM-resolved α-tubulin in T and B cells. FILM acquisition of an anti-tubulin-APC antibody applied to Jurkat T cells (unlabeled cells) and Raji B cells (blue arrowheads) co-cultured on glass and then PFA-fixed and permeabilized. Blue arrowhead labeling is based on an image (not shown) of CMAC staining for pre-labeling of the Raji B cells before setting up the co-culture. Boxed regions are further analyzed in the respective figures or figure panels specified by the labels.

**Fig. S16.**
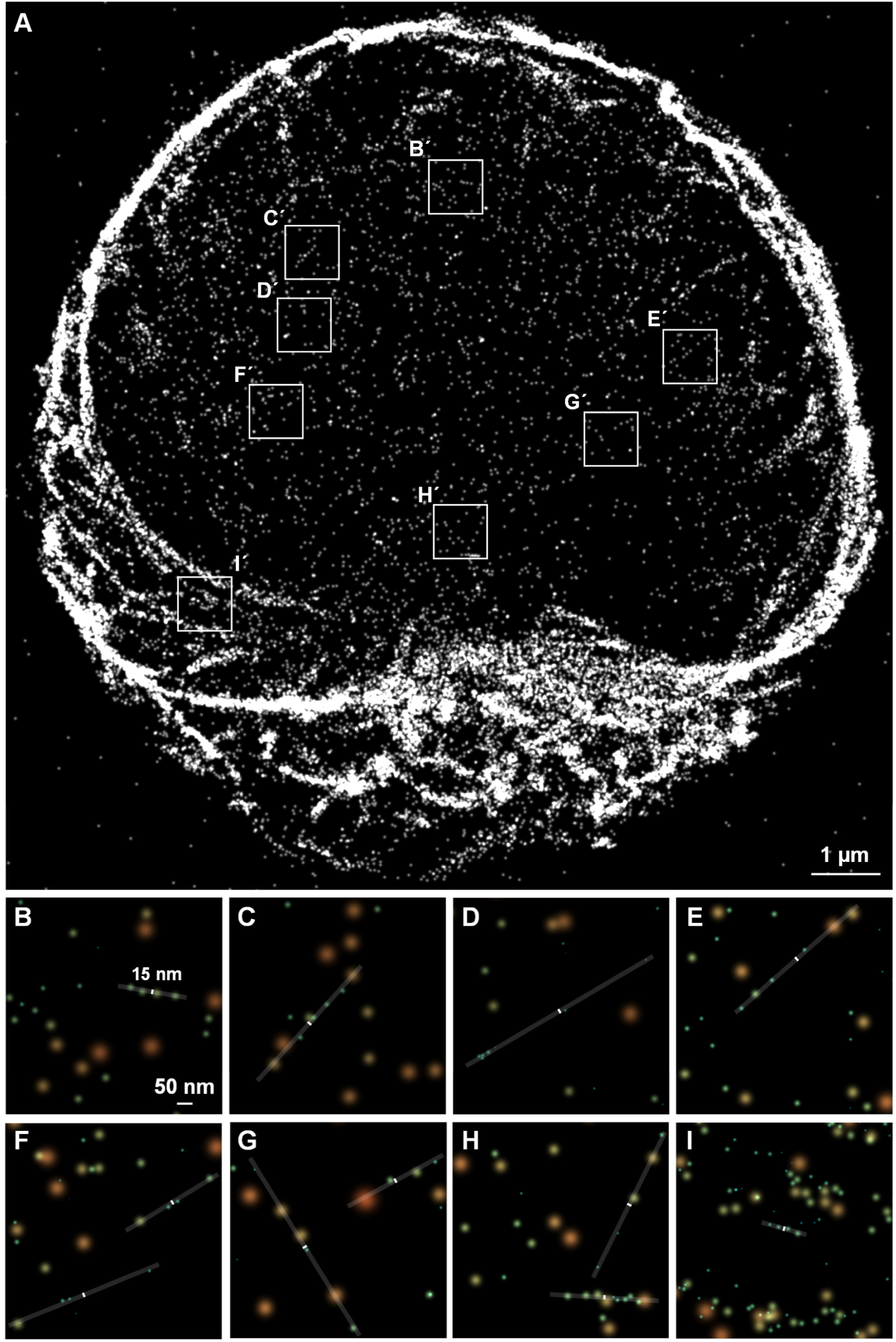
FILM-resolved α-tubulin staining of an isolated T cell. (A) FILM staining of tubulin in an isolated Jurkat T cell (see respective boxed region in Fig. S15). (B–I) Zoom-ins of respective boxed regions in A revealing isolated sets of aligned localizations. Localizations are represented as peak-normalized 2D Gaussians with color-coded precision according to the colorbar shown in Fig. 2E. Almost all of the localized positions (centers of the Gaussians) belonging to each set of nearby aligned localizations could be contained within the overlaid reference strips of width 15 µm.

**Fig. S17.**
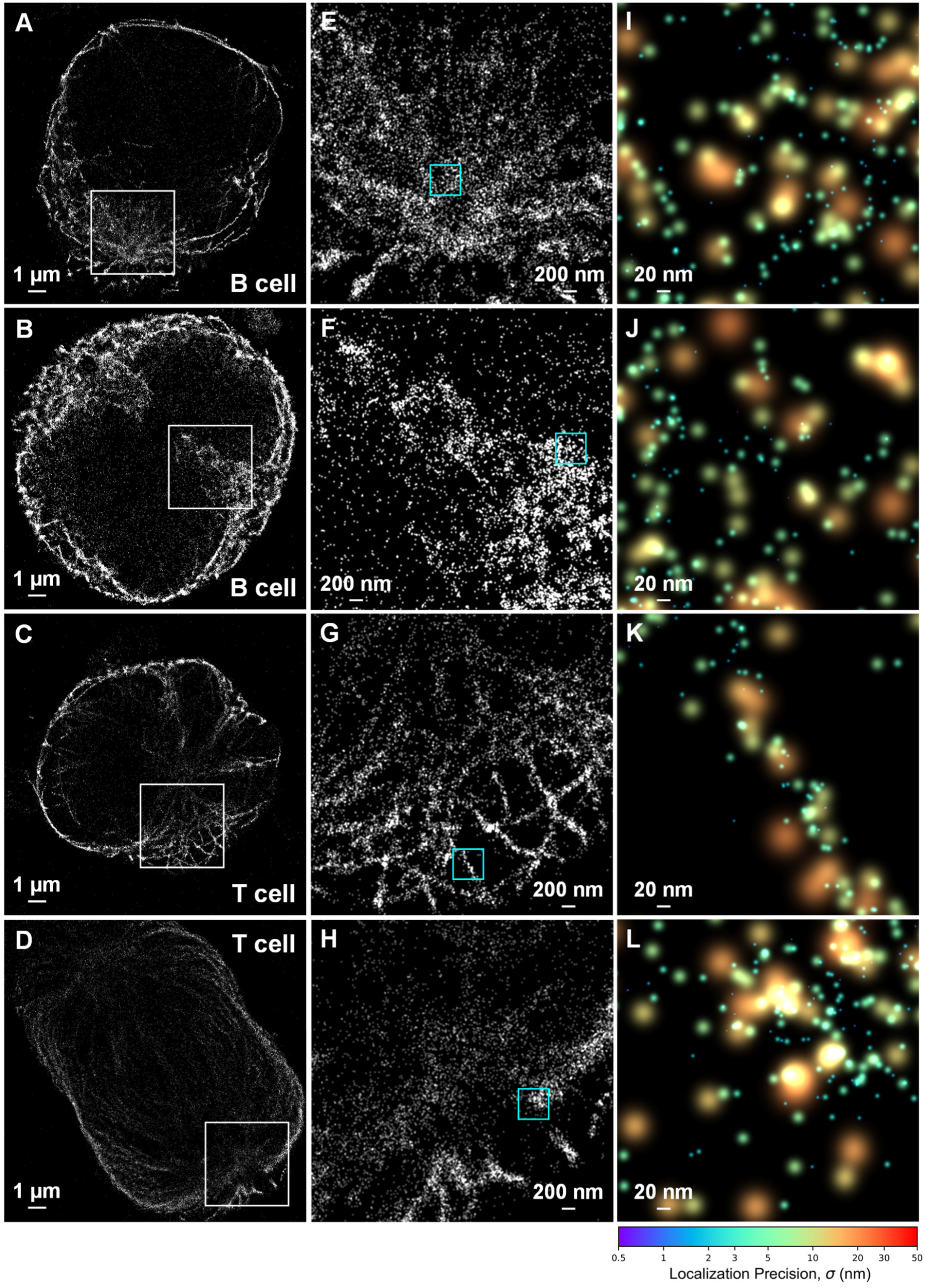
Resolution of α-tubulin in dividing immune cells with FILM. (A–D) Zoom-in of the respectively labeled boxed regions of Fig. S15 containing dividing T and B cells (5 nm pixels with 2-pixel Gaussian blurring). (E–H) Zoom-ins of the respective boxed regions from the frames to the left (5 nm pixels with 2-pixel Gaussian blurring). (I–L) Zoom-ins of the respective cyan boxed regions from the frames to the left. Localizations are represented as peak-normalized Gaussians with color-coded precision according to the colorbar.

**Fig. S18.**
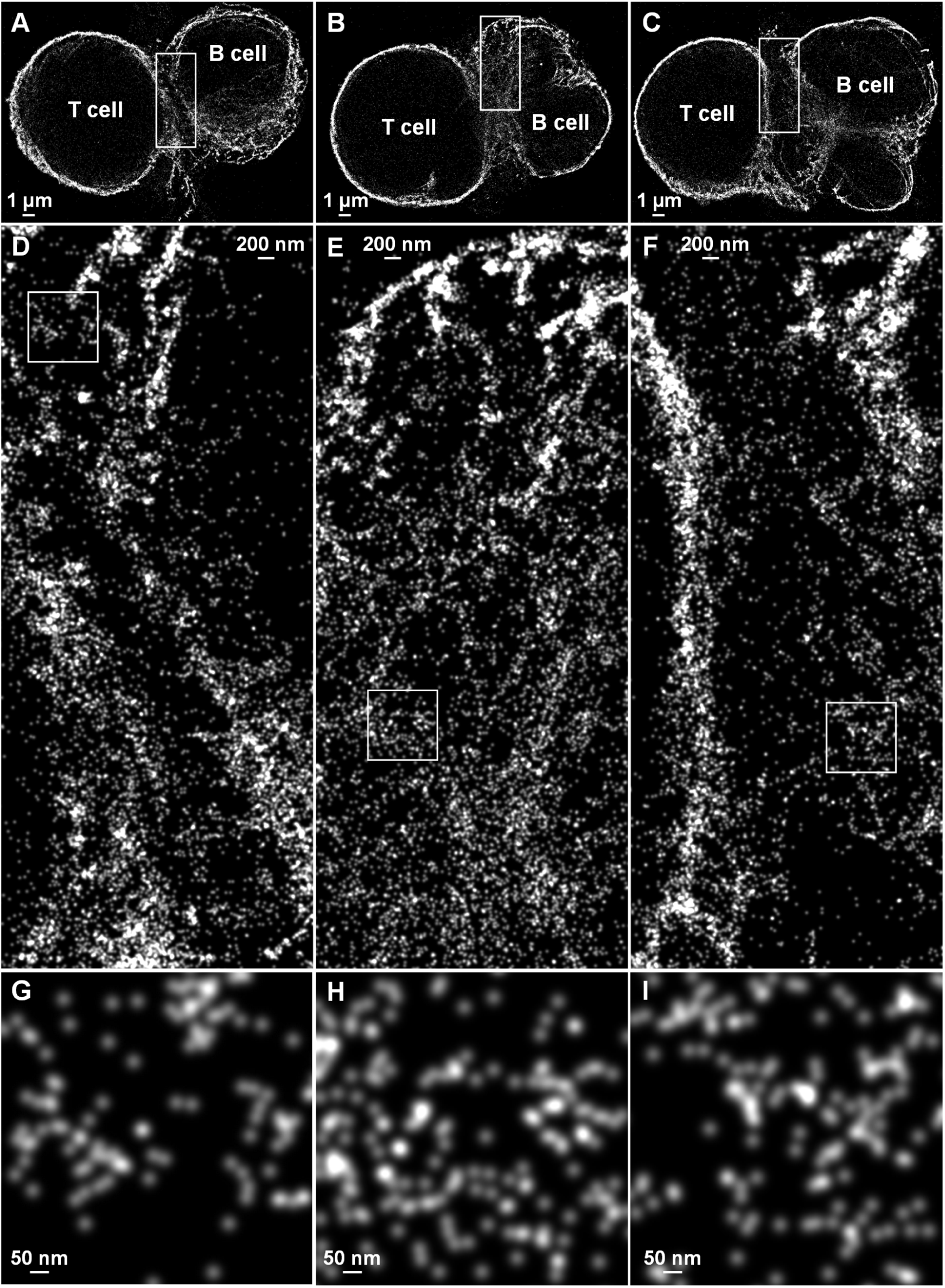
Resolution of α-tubulin in immunological synapses formed between T and B cells with FILM. (A–C) Zoom-in of the respective boxed regions in Fig. S15 containing immunological synapses formed between co-cultured T and B cells. (D–F) Zoom-ins of the respective boxed regions from the frames immediately above. (G–I) Zoom-ins of the respective boxed regions from the frames immediately above. All images have 5 nm pixels and 3-pixel Gaussian blurring to optimally display the continuous cytoskeletal network down to single protofilaments.

**Fig. S19.**
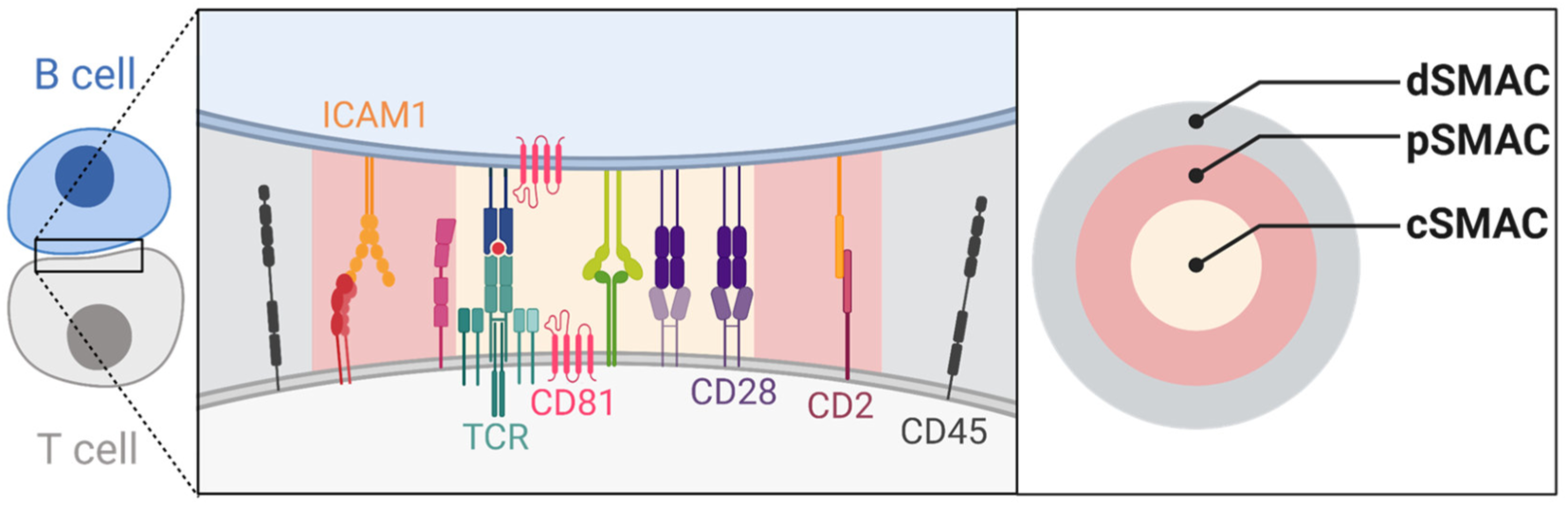
Bullseye structure of a canonical immunological synapse with distinct SMAC zones. Bullseye structure expected for a mature immunological synapse formed between a T cell and a B cell consisting of distinct SMAC zones: distal (dSMAC), peripheral (pSMAC), and central (cSMAC). Proteins known to preferentially localize to each of the zones are indicated. Due to the orientation of the cells on the glass (left and middle panels display the relevant *xy*-perspective for our microscope), the different SMAC zones will appear projected on top of each other. In the right panel, the underlying bullseye structure is revealed using an *xz*-perspective.

**Fig. S20.**
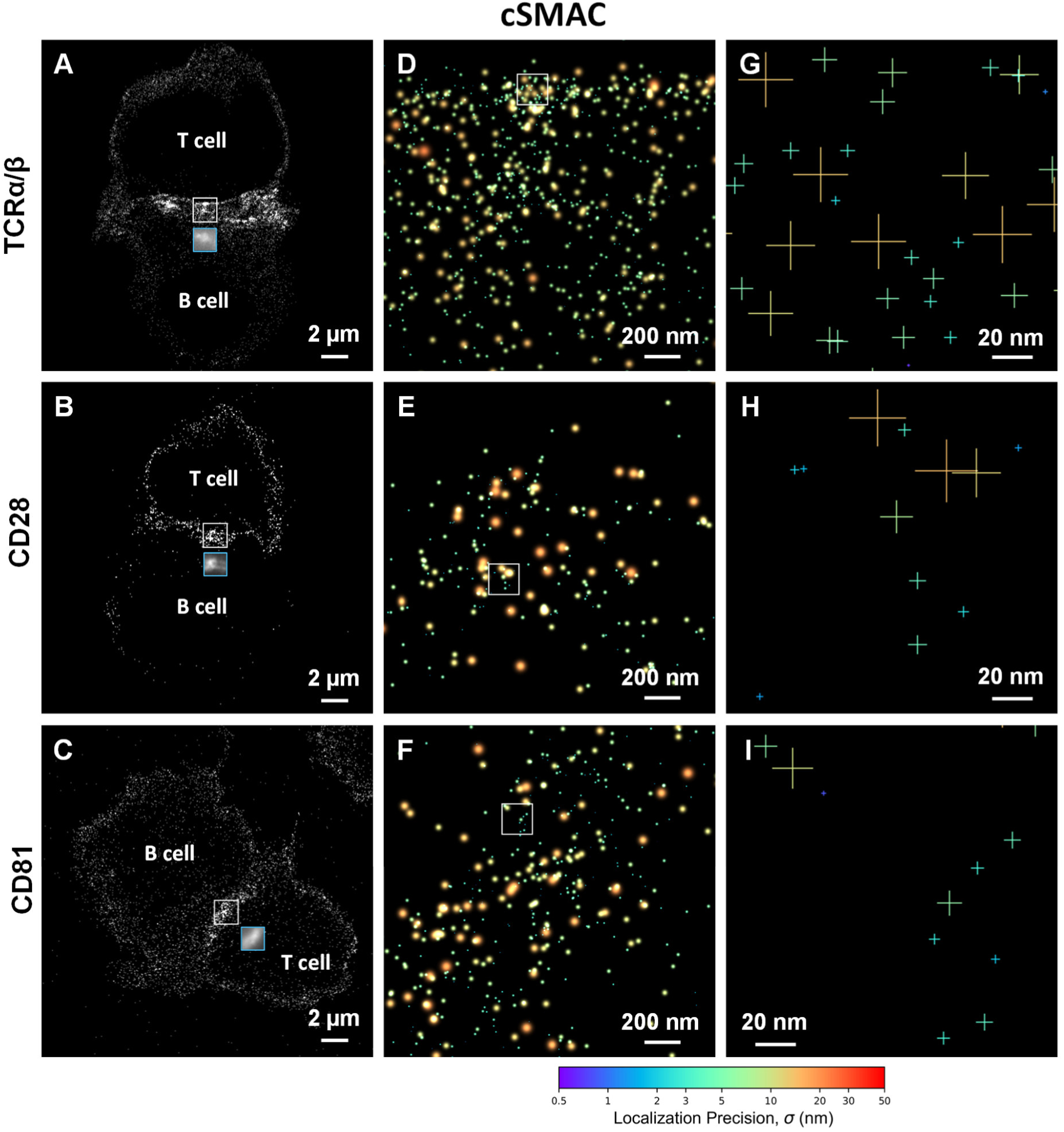
Resolution of markers for the cSMAC zone of an immunological synapse with FILM. (A–C) FILM acquisition of antibodies labeled with APC that recognize TCRα/β, CD28, and CD81 in the cSMAC (10 nm pixels with 3-pixel Gaussian blurring). Cyan boxed region shows the IF image (103.17 nm pixels) for the corresponding white boxed region. (D–F) Zoom-in of the respective boxed regions for the panels to the left. Localizations are shown as peak-normalized 2D Gaussians with color-coded precision according to the colorbar. (G–I) Zoom-in of the respective boxed regions for the panels to the immediate left. Localizations are shown as crosses (indicating ±1-σ uncertainty in *x* and *y*) with color-coded precision according to the colorbar.

**Fig. S21.**
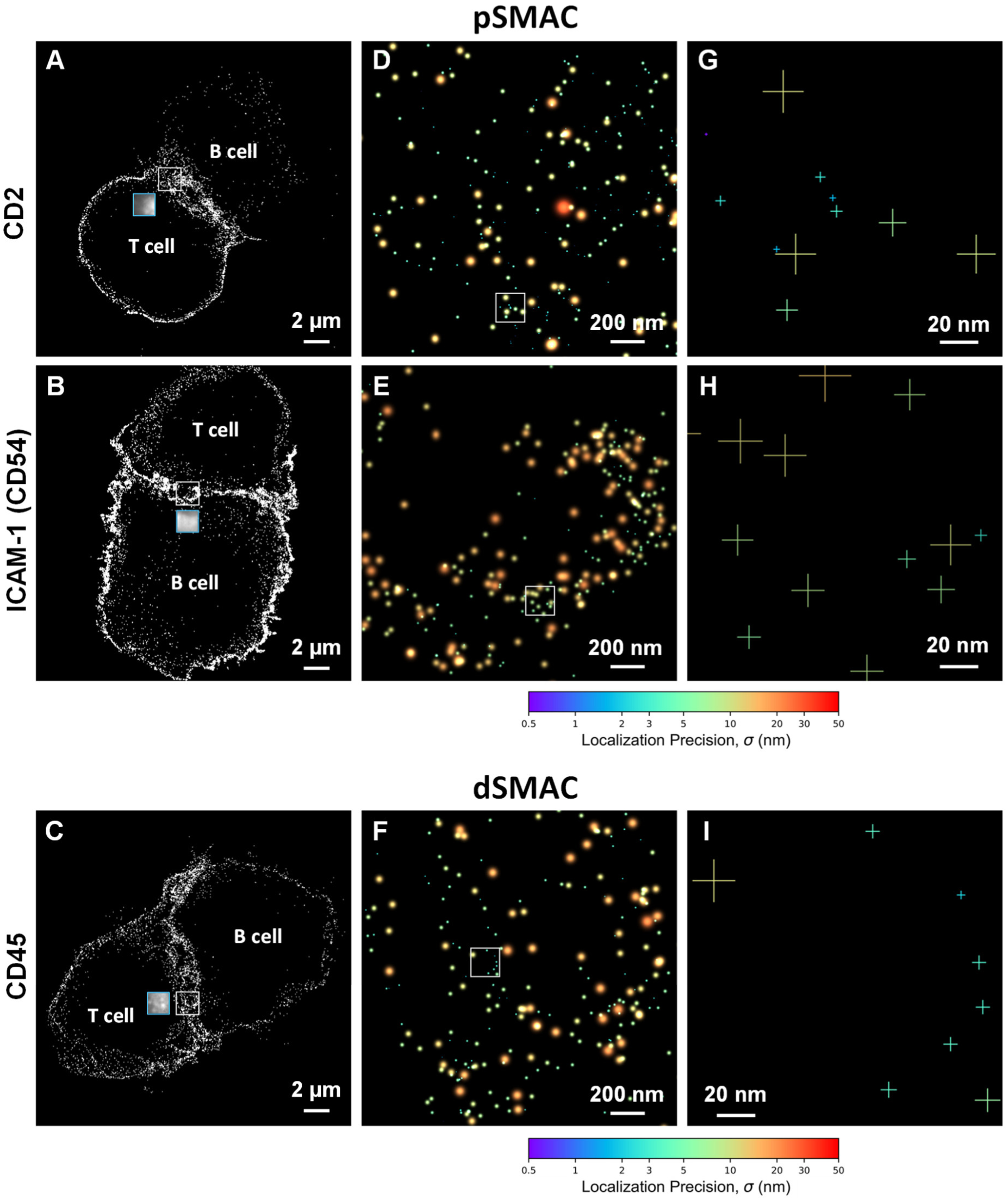
FILM imaging of immunological synapse markers of the pSMAC and dSMAC zones. (A–C) FILM acquisition of binder probes that recognize CD2, ICAM-1 (CD54) in the pSMAC, and CD45 in the dSMAC zones (10 nm pixels with 3-pixel Gaussian blurring). For CD2 and CD45, APC-conjugated antibodies were used; for ICAM-1, a nanobody Fab conjugated with Vio^®^ 667 was used. Cyan boxed region shows the IF image (103.17 nm pixels) corresponding to the white boxed region. (D–F) Zoom-in of the respective boxed regions for the panels to the left. Localizations are shown as peak-normalized 2D Gaussians with color-coded precision according to the colorbar. (G–I) Zoom-in of the respective boxed regions for the panels to the immediate left. Localizations are shown as crosses (indicating ±1-σ uncertainty in *x* and *y*) with color-coded precision according to the colorbar.

**Fig. S22.**
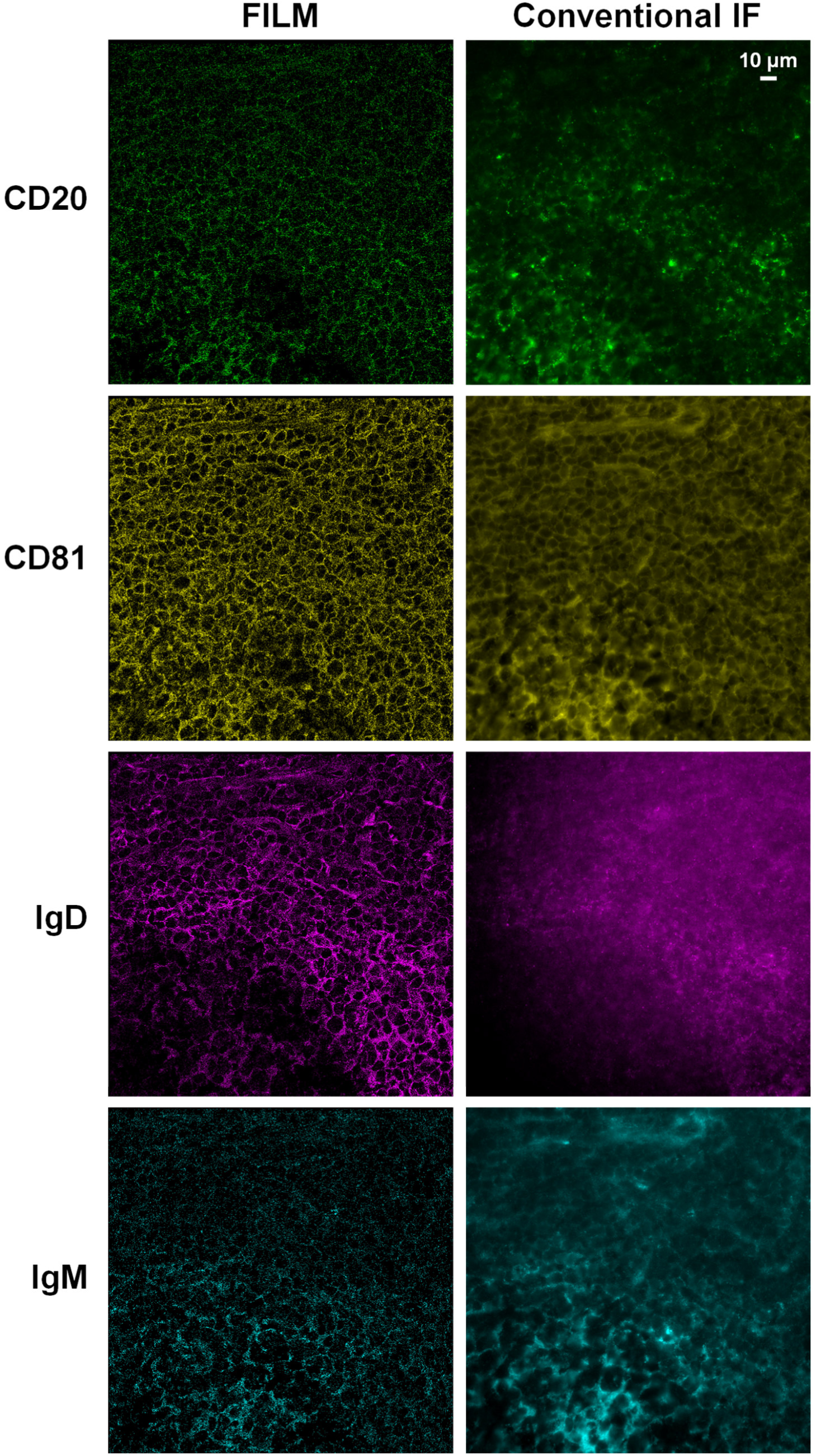
Serial FILM acquisition of four markers in healthy tonsil tissue. Individual stainings for the overlaid FILM and IF images shown in Fig. 4B. To better delineate the stained features over the entire camera field of view, we used for the FILM representation a pixel size corresponding to twice the camera pixel size (206.34 nm). The IF image is displayed with pixels corresponding to the camera pixel size (103.17 nm).

**Fig. S23.**
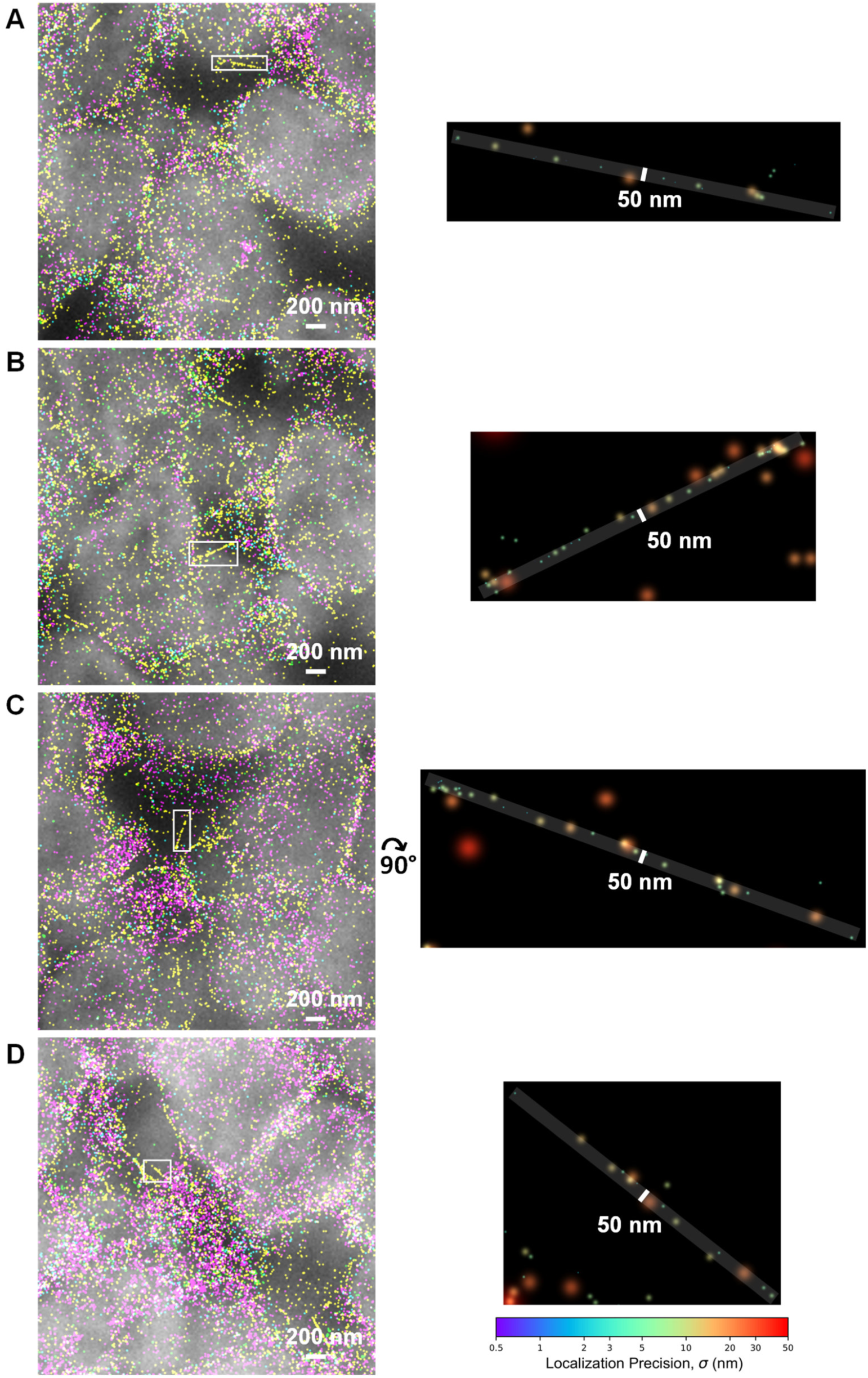
Narrow linear distributions of CD81 in healthy tonsil tissue. (A–D) On the left are further examples of linear distributions of CD81 (see Fig. 4F) that extend out into regions of the tissue with significantly lower nuclear staining. On the right are zoom-ins of the boxed regions indicated in the respective left panels. These depict only the CD81 localizations as peak-normalized 2D Gaussians with color-coded precision given by the colorbar. Almost all of the localizations (centers of the Gaussians) lie within the overlaid 50 nm reference strips. Localizations with significant offsets from the strip were consistent with the otherwise diffuse CD81 staining.

**Fig. S24.**
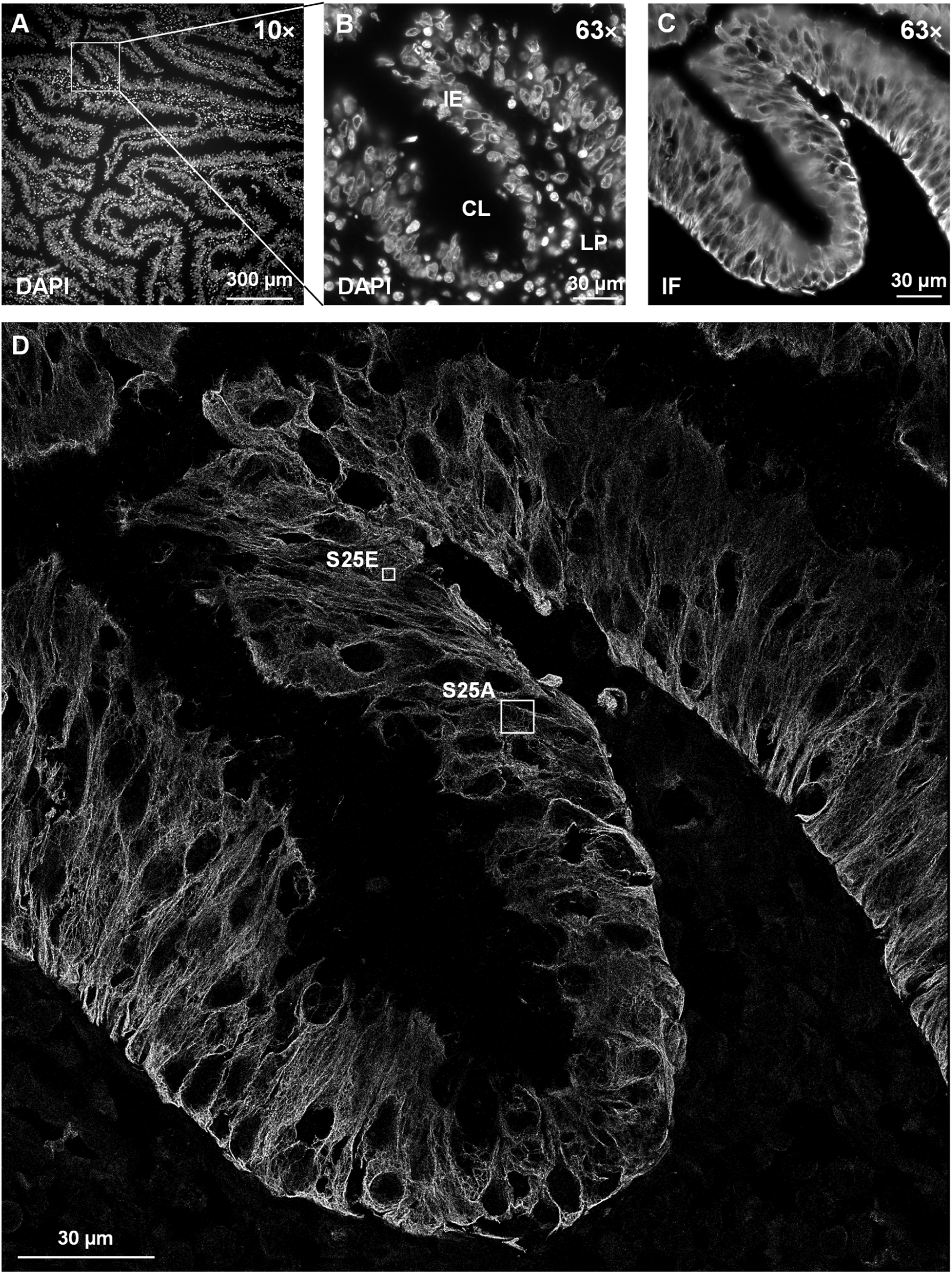
FILM imaging of cytokeratin in colorectal cancer tissue. (A) Overview DAPI image (10× objective) of a CB-fixed, quenched, and permeabilized human colorectal adenocarcinoma tissue slice (4 µm thick). (B) DAPI image (63× objective) of the boxed region in A that includes infiltrative colonic glands with the crypt lumen (CL), intestinal epithelium (IE) and the lamina propria (LP). (C) Conventional IF image of an anti-pan-cytokeratin-APC antibody obtained *after* the FILM acquisition. (D) FILM image of anti-pan-cytokeratin-APC. Panels A–C use 103.17 nm pixels; panel D uses 10 nm pixels with 3-pixel Gaussian blurring.

**Fig. S25.**
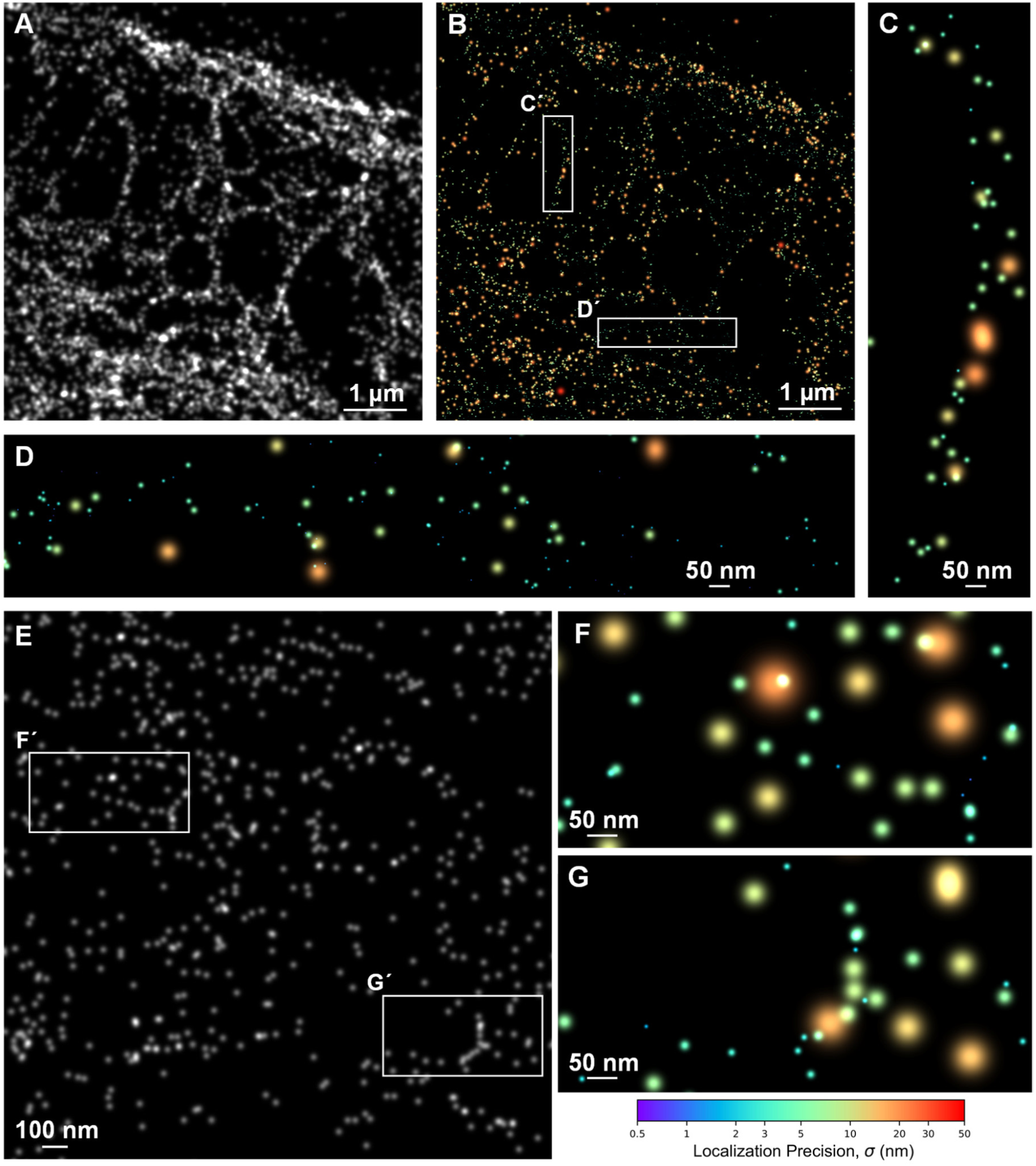
FILM imaging of single cytokeratin filaments in colorectal cancer tissue. (A) Zoom-in of respective boxed region from the FILM image of anti-pan-cytokeratin-APC in colorectal cancer tissue shown in Fig. S24D (10 nm pixels with 3-pixel Gaussian blurring). (B) Region shown in A as peak-normalized 2D Gaussians with color-coded precision according to the colorbar. (C,D) Zoom-ins of boxed regions Ć and D’, respectively, from B showing cytokeratin bundles down to roughly 50–100 nm widths. (E) Zoom-in of respective boxed region from Fig. S24D (5 nm pixels with 2-pixel Gaussian blurring). (F, G) Zoom-ins of the respective boxed regions from E showing narrow branching and curving structures that are sparsely labeled and may correspond to single cytokeratin filaments. For panels C, D, F and G, representation of the events is as described for B.

**Fig. S26.**
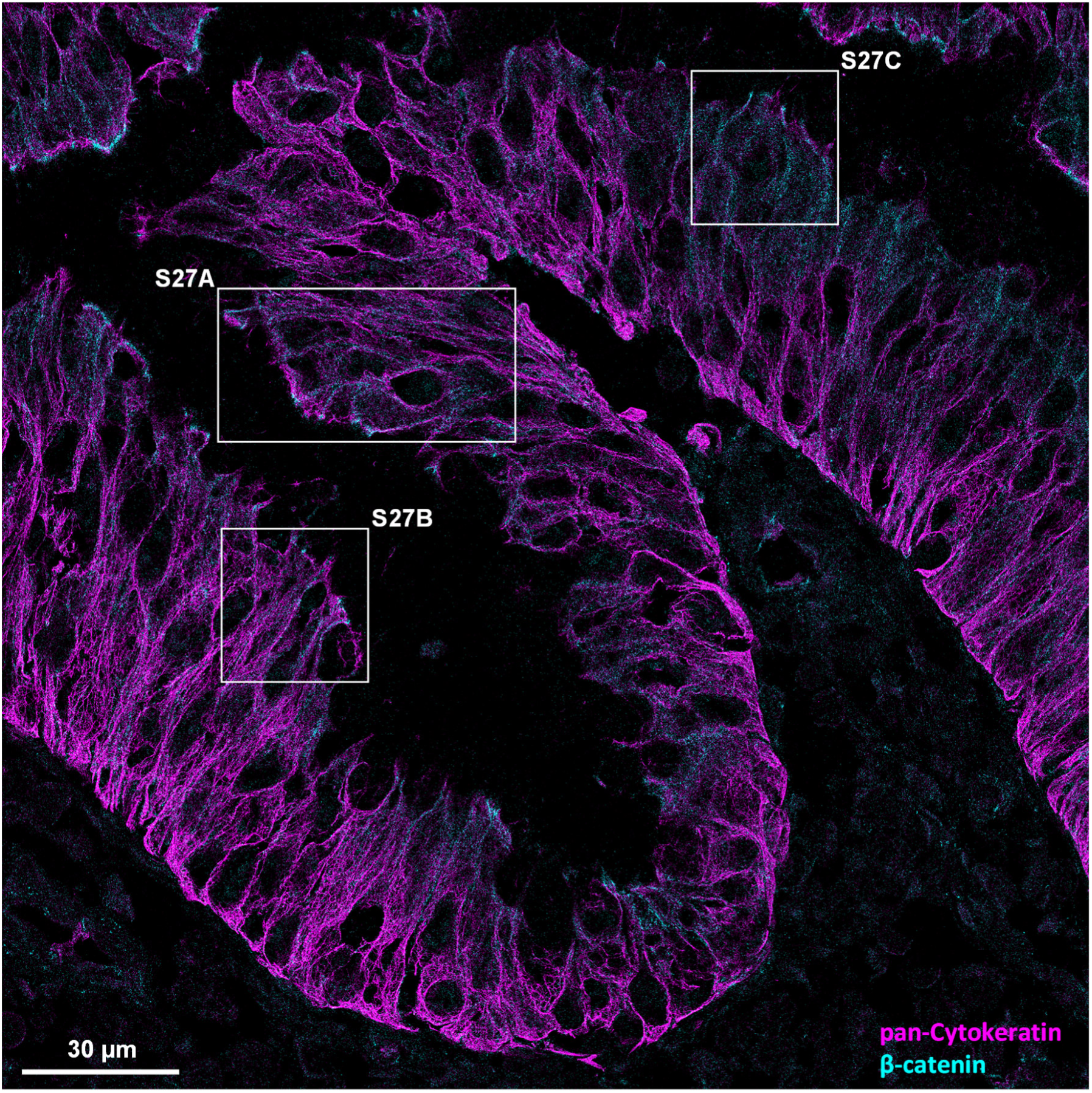
Serial FILM staining of cytokeratin and β-catenin in colorectal cancer tissue. Following cytokeratin staining of the colorectal cancer tissue slice shown in Fig. 24D (shown here in magenta), the tissue was washed and photobleached extensively before initiating a sequential FILM acquisition with an APC-labeled antibody recognizing β-catenin (cyan) (10 nm pixels with 3-pixel Gaussian blurring).

**Fig. S27.**
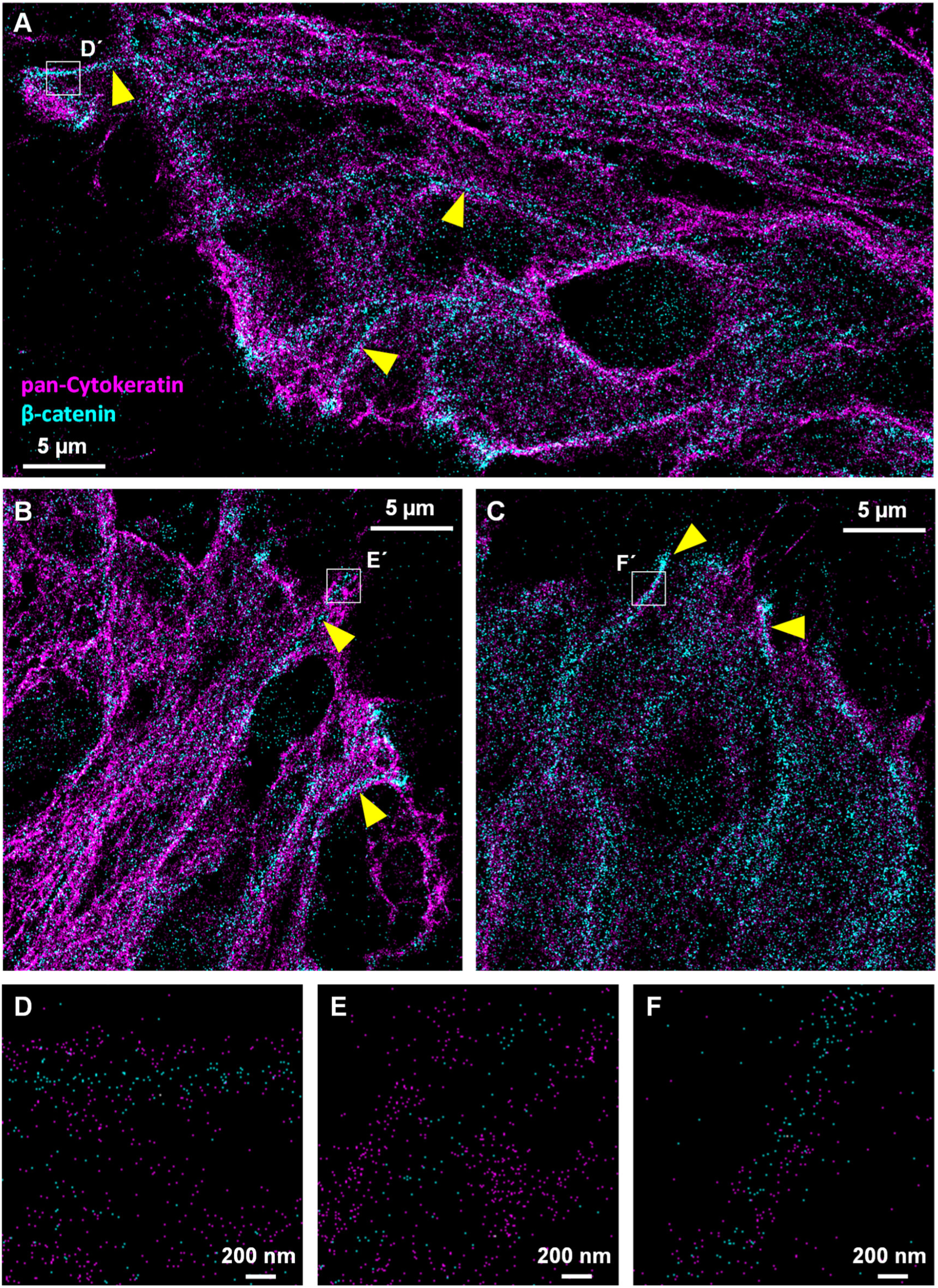
Cytokeratin-flanked channels of β-catenin in colorectal cancer tissue. (A–C) Zoom-in of the respective boxed regions in Fig. S26 showing narrow cytokeratin-free channels of *β*-catenin (∼200 nm in width) flanked by filamentous cytokeratin that terminate at the cell periphery (10 nm pixels with 3-pixel Gaussian blurring). (D–F) Zoom-ins of the respective boxed regions in panels A–C (10 nm pixels).

**Fig. S28.**
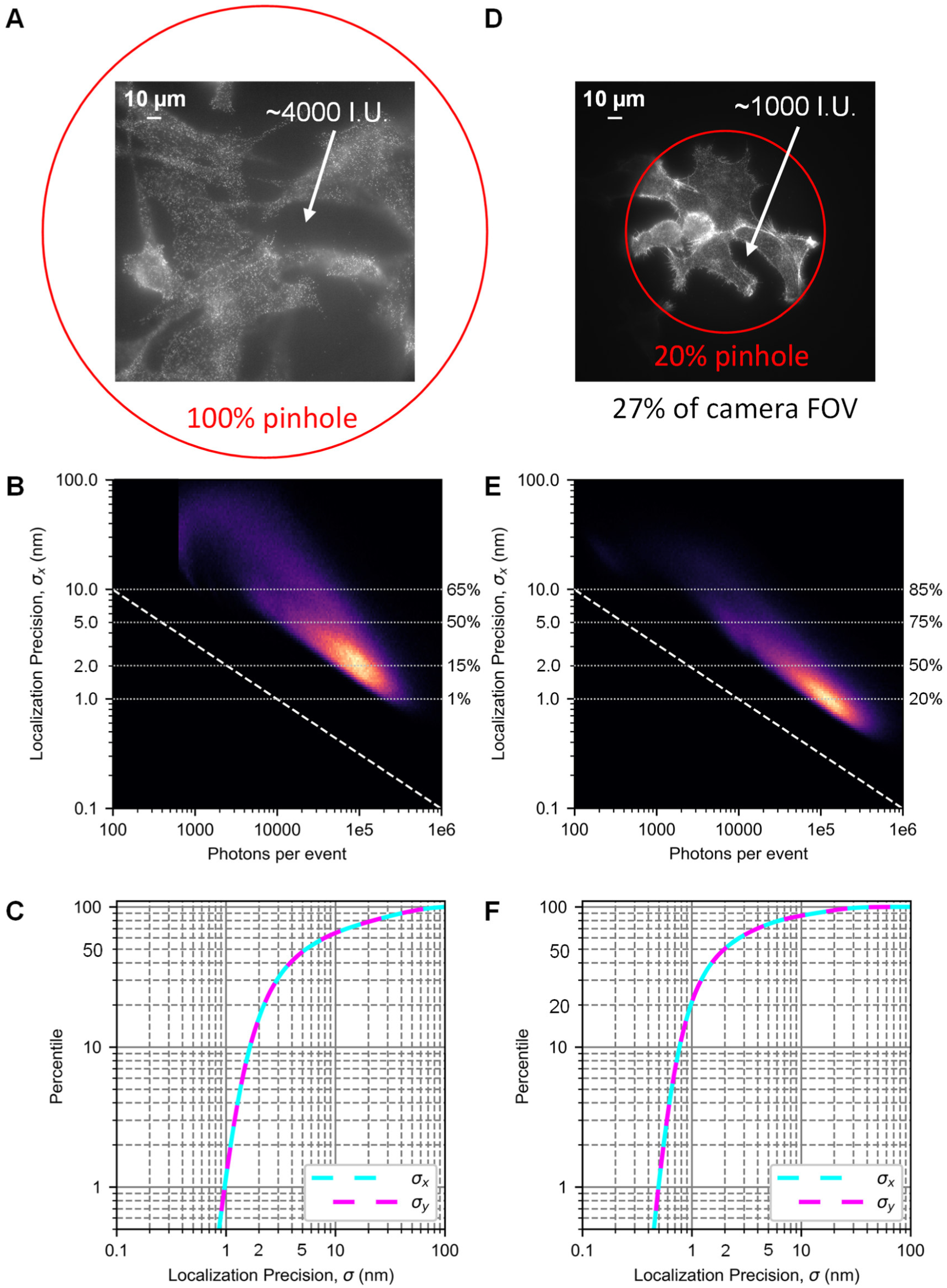
Background reduction using a smaller illuminated FOV allows for higher localization precision. (A) Single 5 s frame from a FILM acquisition of phalloidin-Atto643 labeling fixed HeLa cells using a fully opened excitation pinhole (red circle) (103.17 nm pixels). (B) 2D distribution of localization precision vs. photon counts per probe over the entire acquisition. (C) Cumulative distributions of the localization precision in *x* and *y*. (D–F) Same plots as for A–C, but for a separate experiment on cells in a different microfluidic chamber and using a reduced excitation pinhole of 20%, corresponding to illumination of ∼27% of the total camera FOV.

**Fig. S29.**
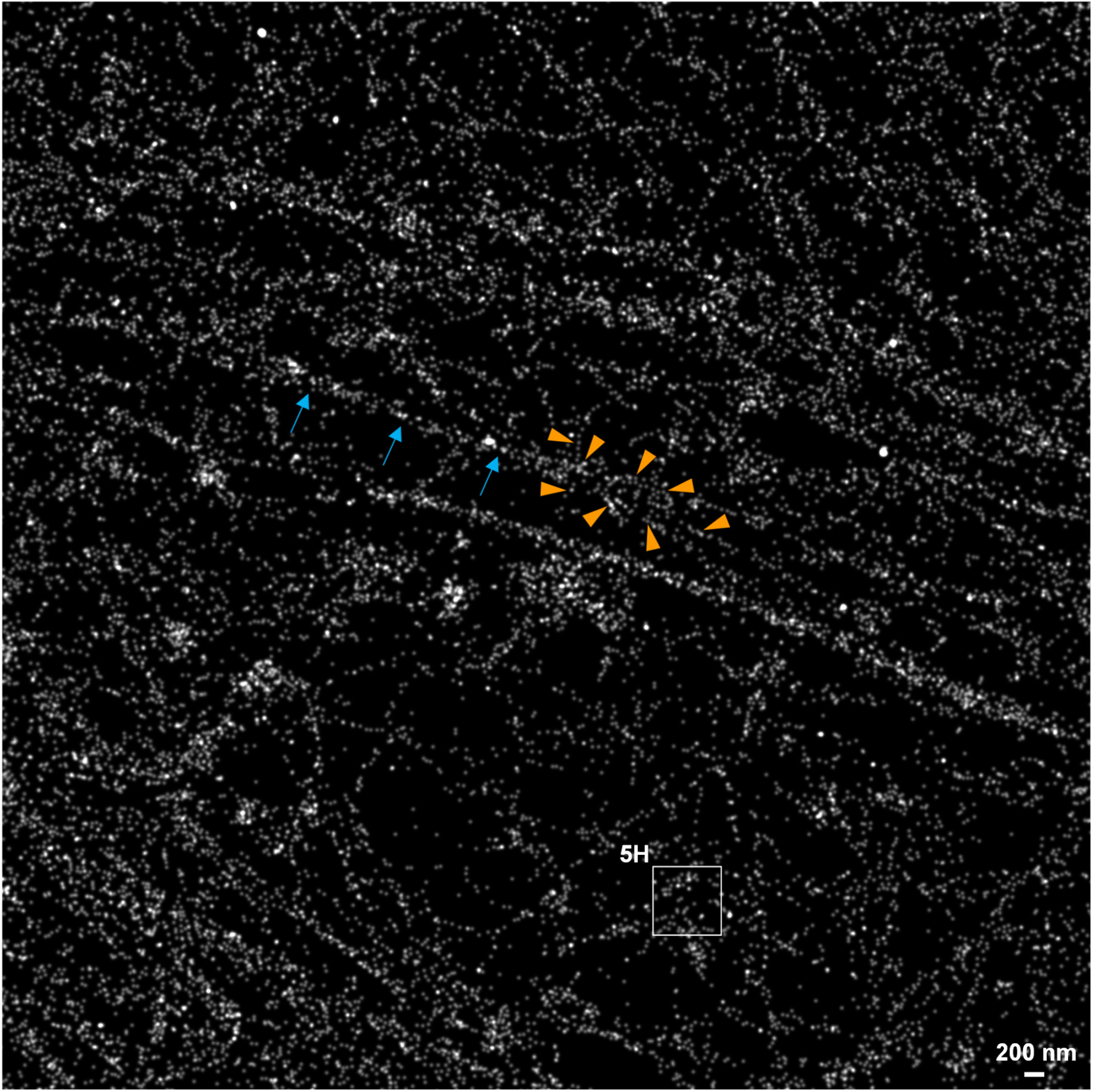
FILM image of the actin network in a HeLa cell. Zoom-in of the respective boxed region in Fig. 5A (5 nm pixels with 3-pixel Gaussian blurring). The indicated actin bundle (blue arrows) is shown to branch off into at least eight thinner actin filaments (orange arrowheads). For the indicated boxed region, containing a highly resolved single actin filament, a zoom-in is given in Fig. 5H.

**Fig. S30.**
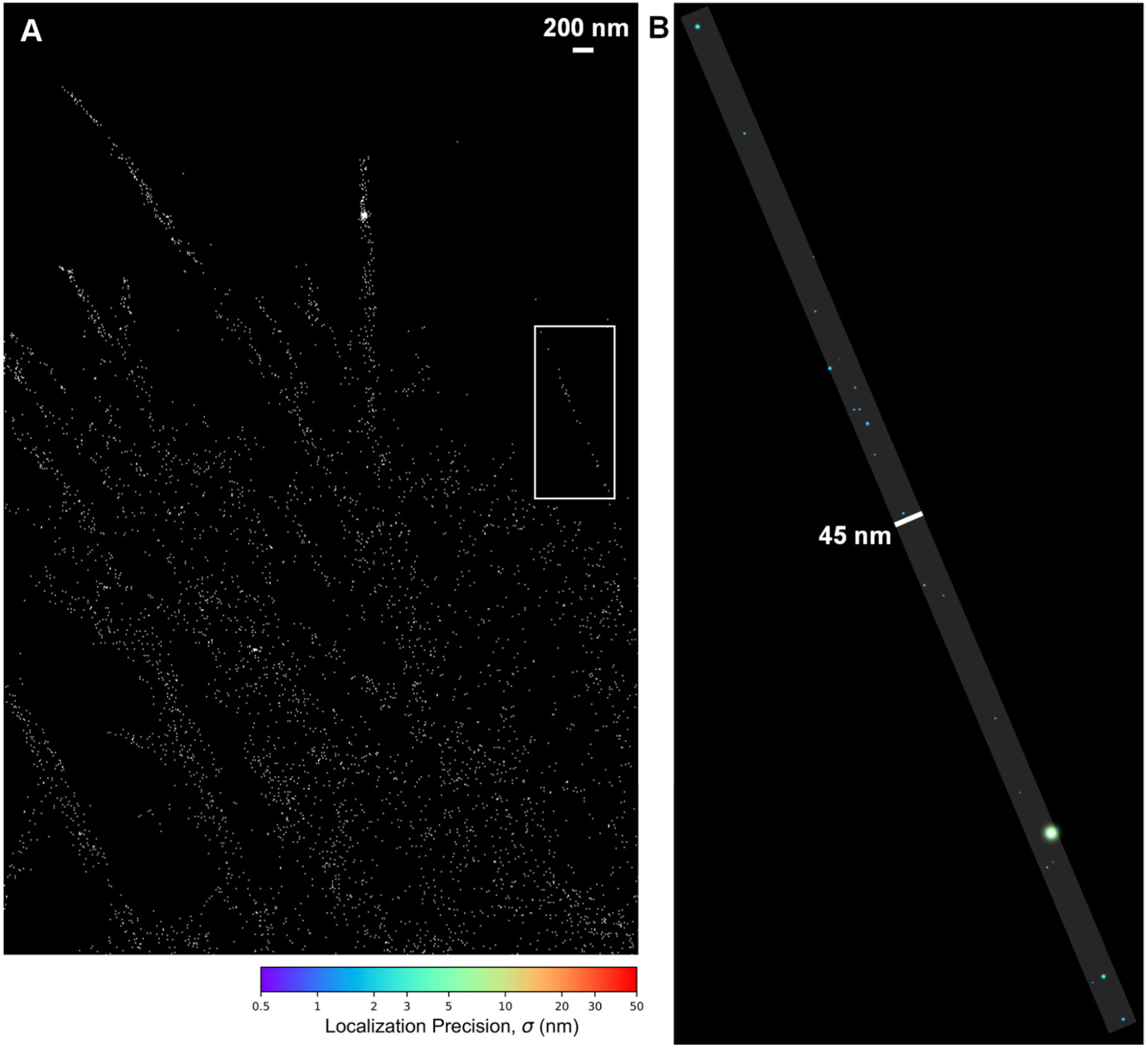
Single actin bundle resolved with phalloidin-Atto643 in a HeLa cell. (A) Zoom-in of the respective boxed region in Fig. 5A (10 nm pixels). (B) Zoom-in of the boxed region in A. Localizations are represented as peak-normalized 2D Gaussians with color-coded precision given by the colorbar. All 22 localizations could be contained within the overlaid 45 nm wide reference strip.

**Fig. S31.**
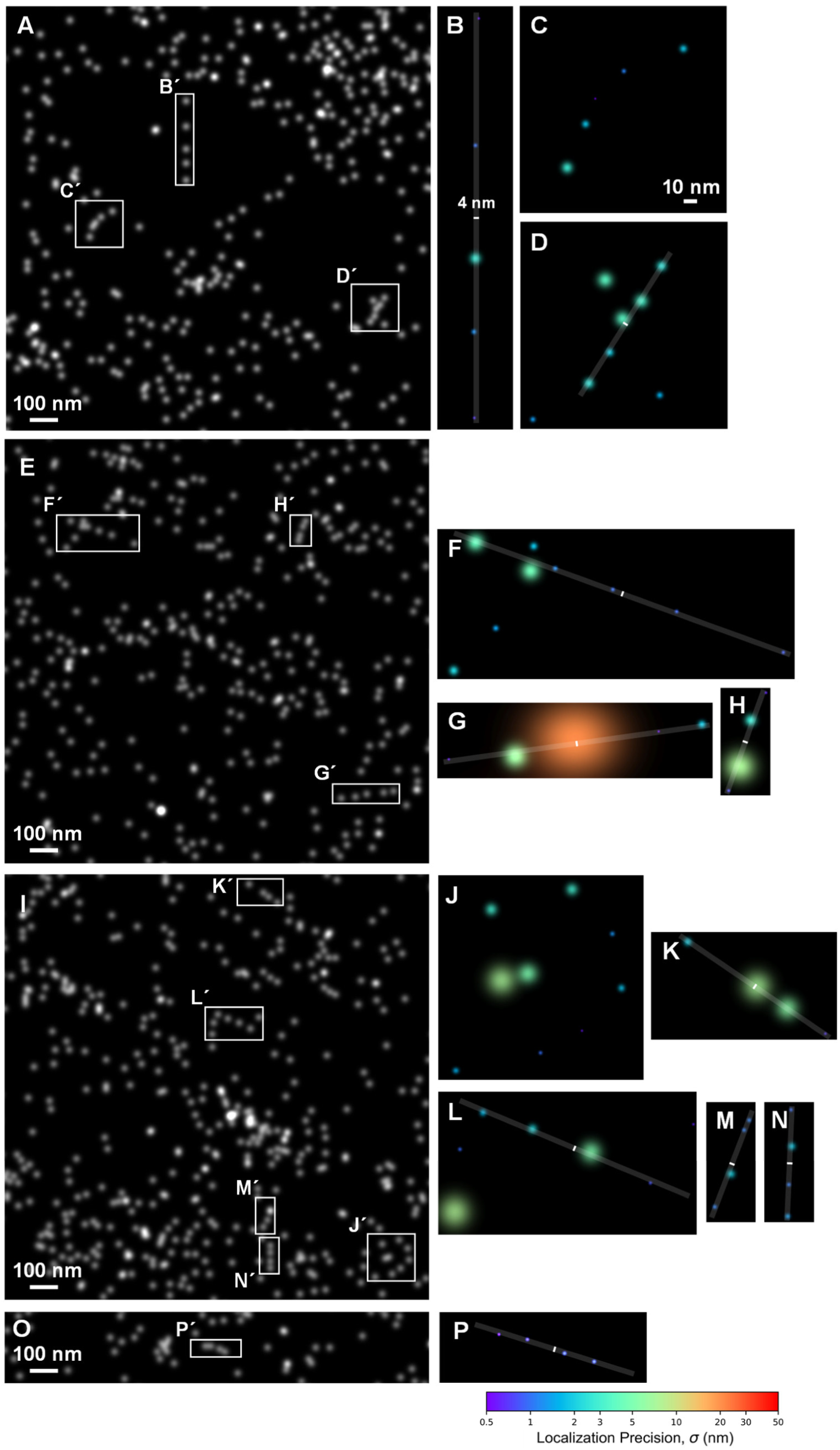
Single actin filaments in HeLa cells resolved with FILM. (A, E, I, O) Zoom-ins of subregions of Fig. 5A (5 nm pixels with 2-pixel Gaussian blurring). (B– D, F–H, J–N, P) Zoom-ins of the respective boxed regions in the panels to the left. Localizations are shown as peak-normalized 2D Gaussians with color-coded precision given by the colorbar. The overlaid strip of 4 nm width in B along with the 10 nm scale bar in C apply generically to this set of panels. Curving filaments are shown in panels C and J (no overlaid strip). For the straight filaments, almost all localizations reside within the overlaid strips. A few less precise localizations (panels G, H, and L) lie slightly outside of the strip (within a few nm of the edge of the strip), consistent with their greater localization uncertainty.

**Fig. S32.**
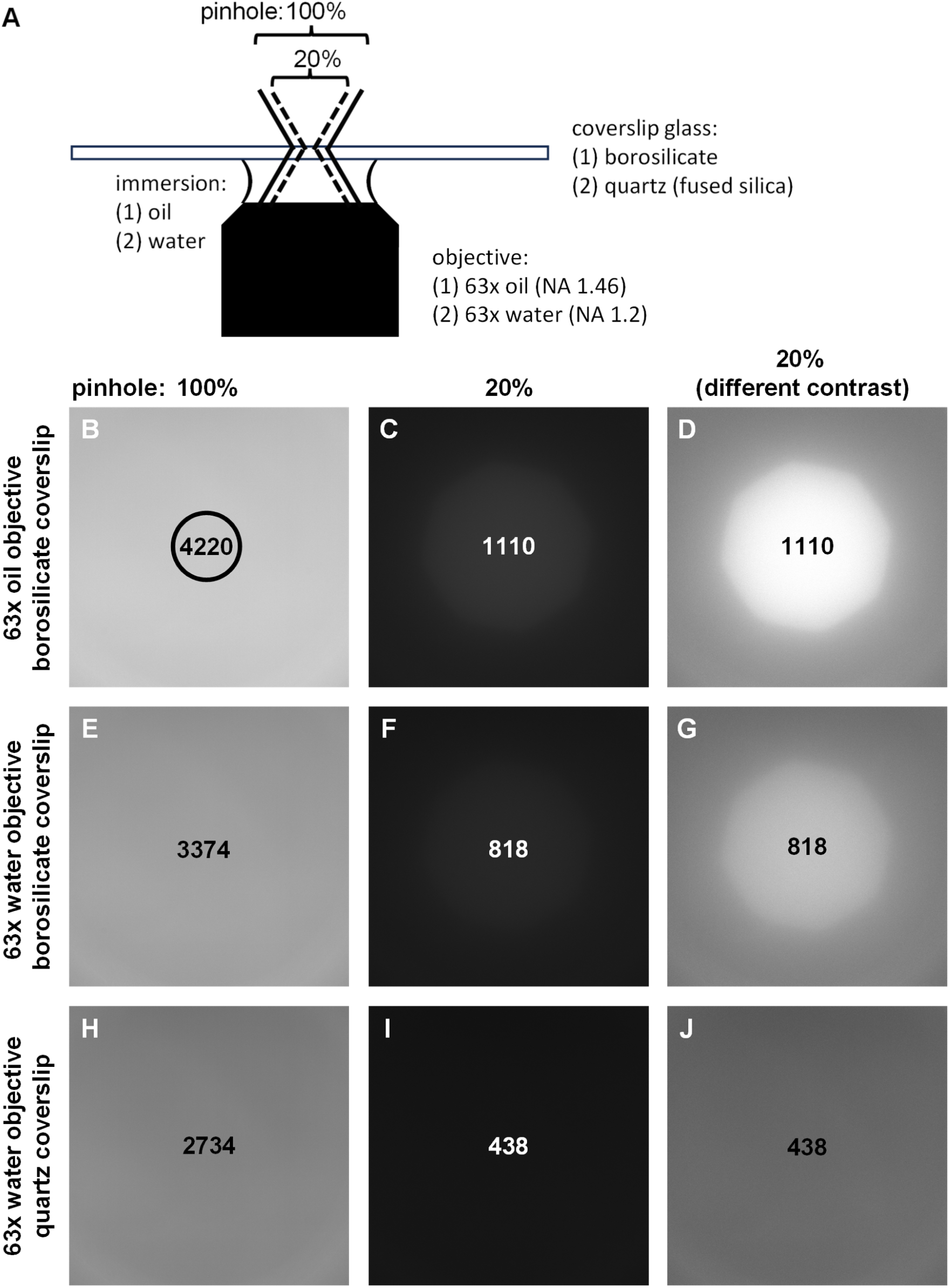
Autofluorescence from the microscope setup. (A) Control measurements were performed to determine the level of homogeneous background on our microscope setup when using different coverslip glasses (either borosilicate glass or quartz glass), objectives (oil- or water-immersion), or illuminated FOVs (excitation pinhole size). (B) Background image (full camera FOV, 211.3×211.3 µm) obtained with the borosilicate coverslip glass and oil-immersion objective for a 100% pinhole. (C) Same as B except for the use of a 20% pinhole. (D) Same as C except with different image contrast. (E) Background obtained with the borosilicate glass and water-immersion objective for a 100% pinhole. (F) Same as E expect for a 20% pinhole. (G) Same as F except with different image contrast. Note the sharp circular profile reflecting a significant autofluorescence contribution from the near-focus portion of the coverslip glass. (H) Background obtained with the quartz coverslip glass and water-immersion objective for a 100% pinhole. (I) Same as H except for the use of a 20% pinhole. (J) Same as I except with different image contrast. Note the absence of the circular profile observed in G for the borosilicate glass, implying the complete absence of autofluorescence signal emanating from the quartz coverslip. All measurements were performed with 100% LED and 5 s exposure. Average integrated intensities were determined in a similar manner for all panels over the circular ROI displayed in B. Oil immersion was used for the oil objective; distilled water was used for the water immersion objective. Image contrast was identical for panels B, C, E, F, H, and I. A different image contrast was applied uniformly for panels D, G, and J. See Materials and Methods and Materials and Methods for further details.

**Fig. S33.**
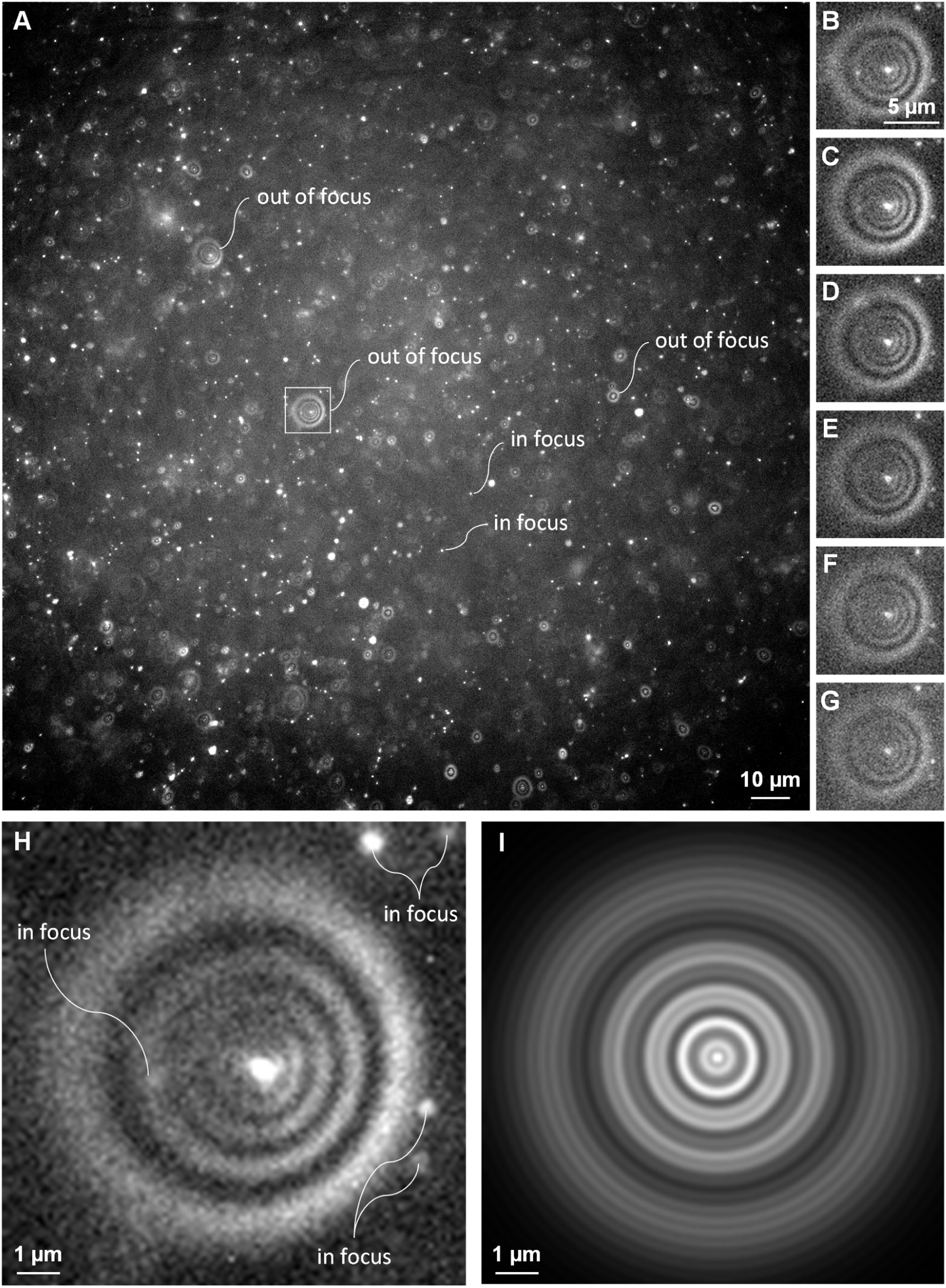
Simultaneous detection of both in-focus and out-of-focus probes with FILM. (A) Representative fluorescence image (5 s exposure; 103.17 nm pixels) from the CD81 dataset (see Fig. 4) showing simultaneous detection of both in-focus and out-of-focus antibody probes conjugated to a single APC. (B–G) Zoom-in of the boxed region in A for the six consecutive frames over which the out-of-focus probe appeared (30 s lifetime). Note the appearance and disappearance of other in-focus localizations. (H) Summed image of all frames from panels B–G, with five diffractive rings clearly discernible. In-focus localizations are indicated that do not belong to the PSF of the out-of-focus probe. Roughly 5.4 million photons were detected for this out-of-focus probe. (I) A Gibson-Lanni model (*60*) for the PSF of a probe located 4.6 µm below the focal plane (with focal plane positioned at 5 µm above the coverslip glass) exhibited a highly similar pattern as that found for the probe in H. The model was calculated using the plugin PSFgenerator (*61*), with values for the positions of the focal plane and probe chosen empirically to give a good match to H. Further values required for the model were the coverslip thickness (170 µm) and the refractive indices of the oil immersion (1.518), borosilicate coverslip glass (1.53) and water-based buffer (1.33).

**Table S1.** Filter combinations used for the different channels referenced to the illumination source.

| **Illumination source** | **Fluorescent dye** | **Excitation filter** | **Dichroic mirror** | **Emission filter** | **Power density at 100%** |
| --- | --- | --- | --- | --- | --- |
| **Colibri 7 LEDs (Carl Zeiss)** |  |  |  |  |  |
| 385/30 nm (UV) | DAPI, CellTracker Blue CMAC | SB 377/50 nm (#FF01-377/50) | SE 409 nm (#FF409-Di03) | SB 447/60 nm (#FF02-447/60) | 14.2 W cm<sup>-2</sup> |
| 555/30 nm (G) | PE | SB 549/17 nm (#FF01-549/17) | SE 561 nm (#Di02-R561) | SB 582/15 nm (#FF01-582/15) | 3.55 W cm<sup>-2</sup> |
| 631/33 nm (R) | APC, Vio® 667 | SB 630/20 nm (#FF01-630/20) | SE 650 nm (#FF650-Di01) | SB 679/41 nm (#FF01-679/41) | 5.33 W cm<sup>-2</sup> |
| **Laser (Rapp OptoElectronic)** |  |  |  |  |  |
| 640 nm | APC, Vio® 667 | SB 631/36 nm (FF01-631/36) | SE 650 nm (FF650-Di01) | LP 665 nm (BLP01-647R) | 676 W cm<sup>-2</sup> |

