## Supplementary material for "Ångström Resolution with Flow Immunofluorescence Localization Microscopy (FILM)": Table S2

Summary of sample preparation, imaging details and analysis parameters for FILM.

| **Sample preparation** | | | **Probe** | | | **Imaging** | | | | | **Analysis with PROSPERO** | | | |
|---|---|---|---|---|---|---|---|---|---|---|---|---|---|---|
| Specimen | Coating | Fixation & perm. | Name | Dilution/ Concentration | Imaging buffer | Flow rate (µL/min) | Total imaging time (h) | Frames | Background estimate (I.U.)[2] | Localizations | Threshold | Maximum radial position deviation (pixels) | Minimum photons | Maximum localization uncertainty (nm) |
| **Figures 2B–G, S3, S5, S7, S9–S10** | | | | | | | | | | | | | | |
| HeLa | PLL | CB + perm. | α-Tubulin-REA1136-APC | 1:5,000 | PBS + 0.1% BSA | 40 | 13.3 | 9,564 | 9,600 | 14,176,305 | 30 | | | |
| **Figures S11–S12** | | | | | | | | | | | | | | |
| HeLa | PLL | CB + perm. | Pan-Cytokeratin-REA1141-APC | 1:5,000 | PBS + 0.1% BSA | 50 | 10.3 | 7,392 | 8,200 | 2,567,779 | 30 | 1 | | |
| **Figure S13** | | | | | | | | | | | | | | |
| HeLa | PLL | PFA + perm. | CD107a/LAMP1-REA792-APC | 1:10,000 | PBS + 0.1% BSA | 35 | 4.2 | 3,000 | 4,300 | 80,200 | 35 | 1 | | |
| **Figure S14** | | | | | | | | | | | | | | |
| HeLa | PLL | PFA + perm. | TOM22-REA1185-APC | 1:4,000 | PBS + 0.1% BSA | 25 | 16.6 | 6,337 | 11,964 | 1,217,485 | 30 | 1 | | |
| **Figure 3, S15–S18** | | | | | | | | | | | | | | |
| T/B cells | PLL | CB + perm. | α-Tubulin-REA1136-APC | 1:10,000 | PBS + 0.1% BSA | 25 | 15.9 | 11,412 | 6,900 | 1,660,723 | 30 | 1 | 2,000 | 20 |
| **Figures S8, S20B, E, H** | | | | | | | | | | | | | | |
| T/B cells | PLL | PFA | CD28-REAL105-APC | 1:6,000 | PBS | 50 | 5.6 | 4,008 | 5,600 | 120,195 | 30 | 1 | 2,000 | 20 |
| **Figures S20A, D, G** | | | | | | | | | | | | | | |
| T/B cells | PLL | PFA | TCRα/β-REA652-APC | 1:5,000 | PBS + 0.1% BSA | 25 | 16.8 | 12,084 | 13,800 | 169,931 | 32 | 1 | 5,000 | 15 |
| **Figures S20C, F, I** | | | | | | | | | | | | | | |
| T/B cells | PLL | PFA + perm. | CD81-REA513-APC | 1:5,000 | PBS + 0.1% BSA | 50 | 15.1 | 10,860 | 16,200 | 361,068 | 32 | 1 | 5,000 | 20 |
| **Figures S21A, D, G** | | | | | | | | | | | | | | |
| T/B cells | PLL | PFA | CD2-REA972-APC | 1:5,000 | PBS | 100 | 7.9 | 5,661 | 6,000 | 71,297 | 30 | 1 | 2,000 | 20 |
| **Figures S21B, E, H** | | | | | | | | | | | | | | |
| T/B cells | PLL | PFA | Fab_CD54-REA266-Vio® R667 | 500 pM | PBS | 25 | 3.4 | 2,421 | 24,300 | 97,727 | 30 | 1 | 2,000 | 20 |
| **Figures S21C, F, I** | | | | | | | | | | | | | | |
| T/B cells | PLL | PFA | CD45-REAL1023-APC | 1:5,000 | PBS | 50 | 4.6 | 3,312 | 9,600 | 44,194 | 30 | 1 | 2,000 | 20 |
| **Figures 4, S6, S22, S23, S33** | | | | | | | | | | | | | | |
| Tonsil tissue | Silane | PFA | IgM-REAL689-PE[1] | 1:5,000 | PBS + 0.1% BSA | 50 | 5.8 | 4,200 | 10,500 | 79,785 | 32 | 1 | 3,000 | 20 |
| | | | IgD-REA740-APC | 1:5,000 | PBS + 0.1% BSA | 50 | 10.3 | 7,404 | 17,100 | 565,910 | 31 | 1 | 4,000 | 20 |
| | | | CD81-REA513-APC | 1:10,000 | PBS + 0.1% BSA | 50 | 3.2 | 2,268 | 8,500 | 454,641 | 35 | 1 | 4,000 | |
| | | | CD20-REA780-APC | 1:10,000 | PBS + 0.1% BSA | 50 | 3.5 | 2,527 | 27,500 | 95,078 | 30 | 1 | 4,000 | 20 |
| **Figures S24–S27** | | | | | | | | | | | | | | |
| Colorectal cancer tissue | Silane | CB + perm. | Pan-Cytokeratin-REA1141-APC | 1:10,000 | PBS + 0.1% BSA | 50 | 10.9 | 7,848 | 6,500 | 3,072,762 | 30 | 1 | 2,000 | 20 |
| | | | β-catenin-REA480-APC | 1:10,000 | PBS + 0.1% BSA | 50 | 7.5 | 5,364 | 8,200 | 451,742 | 30 | 1 | 2,000 | 20 |
| **Figures 5, S28D–F, S29–S31** | | | | | | | | | | | | | | |
| HeLa | PLL | PFA | Phalloidin-Atto643 | 20 pM | PBS | 40 | 7.9 | 5,712 | 1,000[3] | 1,475,520 | 30 | | | |
| **Figures S28A–C** | | | | | | | | | | | | | | |
| HeLa | PLL | PFA | Phalloidin-Atto643 | 20 pM | PBS | 40 | 29.5 | 21,275 | 4,000 | 10,993,681 | 30 | | | |

BSA = bovine serum albumin, CB = cytoskeleton-preserving buffer, PBS = Phosphate Buffered Saline (pH 7.4) , perm. = permeabilization with 0.1 % (v/v) Triton X-100 in PBS, PFA = paraformaldehyde, PLL = poly-L-lysine, T/B = T/B cell co-culture (Jurkat T cells with Raji B cells), REA = REAfinity™ clone, REAL = REAdye_lease™ clone, I.U. = camera intensity units, [1]Data taken with the PE channel settings and 20% LED power, with all other data using the APC channel and 100% LED power (see Table S1). [2]Background from imaging the borosilicate coverslip glass alone was ~4000 (100% excitation pinhole) and ~1000 (20% excitation pinhole). [3]Reduced-background dataset using a 20% excitation pinhole (all other datasets employed a 100% excitation pinhole).
